# KAT2-mediated acetylation switches the mode of PALB2 chromatin association to safeguard genome integrity

**DOI:** 10.1101/735811

**Authors:** Marjorie Fournier, Jean-Yves Bleuyard, Anthony M. Couturier, Jessica Ellins, Svenja Hester, Stephen J. Smerdon, László Tora, Fumiko Esashi

## Abstract

The tumour suppressor PALB2 stimulates error-free repair of DNA breaks, whilst its steady-state chromatin association protects active genes from genotoxic stress. Here, we report that the lysine acetyltransferases 2A and 2B (KAT2A/B), commonly known to promote transcriptional activation, acetylate the PALB2 chromatin association motif (ChAM), providing a dynamic regulatory mechanism for PALB2. ChAM acetylation within a cluster of seven lysine residues (7K), detected in the chromatin-enriched fraction in undamaged cells, enhanced its association with nucleosomes while decreasing its non-specific binding to naked DNA. DNA damage triggered a rapid deacetylation of ChAM and a concomitant increase in PALB2 mobility. Significantly, a 7K-null mutation, which hindered ChAM binding to both nucleosomes and DNA, conferred deficiency in DNA repair and hypersensitivity to the anti-cancer drug olaparib. Thus, our study reveals a unique mechanism mediated by KAT2A/B-dependent acetylation of a non-histone protein, which fine-tunes the DNA damage response and hence promotes genome stability.

## Introduction

PALB2, the partner and localizer of the breast cancer susceptibility 2 protein (BRCA2) (Xia et al, 2006), plays essential roles in the maintenance of cellular homeostasis and disease prevention in humans. Biallelic mutations in *PALB2* cause Fanconi anaemia (FA), a rare genetic disorder characterised by bone marrow failure, developmental abnormalities and increased incidence of childhood cancers (Reid et al, 2007; Xia et al, 2007). Hereditary monoallelic *PALB2* mutations also increase the risk of breast and pancreatic cancer (Jones et al, 2009; Rahman et al, 2007), similarly to inherited *BRCA1* and *BRCA2* mutations (O’Donovan & Livingston, 2010). PALB2 works together with BRCA1 and BRCA2 to promote the error-free repair of highly genotoxic double-strand DNA breaks (DSBs) by homologous recombination (HR). Current models propose that BRCA1 acts as a DNA damage sensor, which in turn recruits PALB2 and BRCA2 to sites of DNA damage. The essential RAD51 recombinase is subsequently recruited and catalyses the strand invasion and homology search phases of HR repair (Sy et al, 2009b; Xia et al, 2006).

We have recently revealed a repair-independent role of PALB2 in protecting transcriptionally active chromatin from DNA damage arising from replicative stress (Bleuyard et al, 2017b). This role of PALB2 is mediated through its high-affinity binding partner the MORF-related gene 15 protein (MRG15), which recognises an epigenetic marker of active genes, histone H3 trimethylated at lysine 36 (H3K36me^3^), *via* its N-terminal chromodomain (Bleuyard et al, 2017b; Hayakawa et al, 2010; Sy et al, 2009a). Moreover, PALB2 intrinsic chromatin association is reinforced by the chromatin-association motif (ChAM), an evolutionarily conserved domain uniquely found in PALB2 orthologues, which directly binds to nucleosomes (Bleuyard et al, 2012). Notably, our locus-specific chromatin immunoprecipitation (ChIP) analyses revealed a decrease in PALB2 association with active genes following an exposure to a topoisomerase inhibitor camptothecin (CPT), suggesting that the mode of PALB2 chromatin association is actively regulated (Bleuyard et al, 2017b). Despite these observations, the regulatory mechanism by which PALB2 is switched between different modes of chromatin association (i.e. association with damaged DNA to promote HR repair and with actively transcribed genes to assist in their protection) remains elusive.

Numerous studies in recent decades have provided evidence that reversible post-translational modifications (PTMs), such as phosphorylation, ubiquitylation, SUMOylation, poly(ADP-ribosyl)ation, methylation and acetylation, are orchestrated to promote genome stability, including the DNA damage response (DDR) (Dantuma & van Attikum, 2016). For example, the damage-responsive ATM and ATR kinases mediate phosphorylation of PALB2 at residues S59, S177 and S376, which in turn facilitate PALB2 interaction with BRCA1, RAD51 foci formation, and hence HR repair of DSBs (Ahlskog et al, 2016; Buisson et al, 2017; Guo et al, 2015). Conversely, in G1 phase, the E3 ligase KEAP1-CUL3-RBX1 ubiquitylates PALB2 at K25, a key residue involved in BRCA1 interaction, and in this way, suppresses PALB2-BRCA1 interaction and HR activation (Orthwein et al, 2015). Furthermore, our recent work identified PALB2 as a key substrate of the lysine acetyltransferases 2A (GCN5/KAT2A) and 2B (PCAF/KAT2B) (Fournier et al, 2016), two well-known transcriptional regulators (reviewed in Nagy et al, 2010), in undamaged cells. However, the physiological role of these modifications is as yet unknown. Notably, KAT2A/B use the metabolite acetyl coenzyme A (acetyl-CoA) as a cofactor (Tanner et al, 2000), and hence are proposed to fine-tune cellular processes in accordance with the metabolic status of the cell (Wellen et al, 2009). The study of acetylation would have important implications in the context of tumorigenesis, as cancer cells are characterised by reprogrammed metabolism and elevated genome instability.

In this study, we investigated the role of KAT2A/B-mediated lysine acetylation in regulating PALB2. We found that KAT2A/B acetylate a cluster of seven lysine residues (the 7K-patch) within the PALB2 ChAM. ChAM acetylation enhanced its association with nucleosomes, while restricting its non-specific binding to naked DNA. Notably, DNA damage triggered rapid ChAM deacetylation and a concomitant increase in PALB2 mobility. Importantly, lysine to glutamine substitutions in the 7K-patch impeded PALB2 association with chromatin and its function in the DDR. On the basis of these observations, we propose that PALB2 chromatin association is actively regulated by KAT2A/B, which hence play a significant role in the maintenance of genome stability.

## Materials and Methods

### Cell culture and cell lines

All cells were grown at 37°C in an incubator with a humidified atmosphere with 5% CO_2_. HEK293T cells were grown in Dulbecco’s Modified Eagle’s Medium (DMEM, Sigma) supplemented with 10% fetal bovine serum (FBS), 100 U/mL penicillin, and 0.1 mg/mL streptomycin. U2OS Flp-In T-REx P2shRNA cell line (Bleuyard et al, 2017b), carrying a doxycycline-inducible shRNA targeting the endogenous *PALB2* 3’ UTR, referred to as U2OS-shPALB2, were used to generate stable isogenic cell lines with constitutive expression of N3xFLAG-PALB2 variants.

U2OS-shPALB2 were co-transfected with pOG44 and pcDNA5/FRT GW/N3×FLAG-PALB2 (7Q or 7R), and resultant stable cell lines were selected in DMEM supplemented with 10% FBS, 100 U/mL penicillin, 0.1 mg/mL streptomycin, 10 µg/mL blasticidin, 200 μg/mL hygromycin B (100 µg/mL to maintain the cell lines), and 1 µg/mL puromycin. Stable U2OS-shPALB2 lines carrying the empty pcDNA5/FRT GW/N3×FLAG vector or the pcDNA5/FRT GW/N3×FLAG-PALB2 WT vector have been previously described (Bleuyard et al, 2017b).

### Antibodies

The primary antibodies used (with their respective working dilutions) were: anti-FLAG (Sigma F1804, mouse, WB: 1/1000), anti-pan-acetyl lysine (AcK) (Cell Signaling Technology 9441S, rabbit, WB: 1/1000), anti-PALB2 (Bethyl A301-246A, rabbit, WB: 1/500), anti-BRCA2 (Millipore OP95, mouse, WB: 1/1000), anti-RAD51 (Yata et al, 2014) (rabbit, 7946 IF: 1/1000), anti-lamin A (Sigma L1293, rabbit, WB: 1/2000), anti-γ-H2AX (Millipore 05-636, mouse, WB: 1/1000), anti-GFP (Sigma G1544, mouse, WB: 1/1000), anti-histone H3 (Bethyl A300-823A, rabbit, WB: 1/1000), anti-GST (Santa Cruz Biotechnology sc-138, mouse, WB:1/1000), biotin-HRP conjugated (Sigma A0185, mouse, WB:1/1000), anti-KAT2A/GCN5 (Cell Signaling Technology 3305, rabbit, WB:1/1000), anti-α-tubulin (Cell Signaling Technology 3873, mouse, WB: 1/2000). Secondary antibodies coupled with horseradish peroxidase (HRP): goat anti-mouse (Dako P0447, WB: 1/1000), goat anti-rabbit (Dako P0448, WB: 1/1000).

### DNA damage and cell treatment

For ionising radiation-induced DNA damage, cells were exposed to 4 Gy γ-rays using a ^137^Cs source, delivering a dose rate of 1.68 Gy/min (Gravatom). For genotoxic drug treatments, cells were incubated for 2 h at 37°C with either 1 mM MMC (Sigma M0503) or 2.5 μM olaparib (Enzo Life Sciences LKT-O4402). DMSO was used as negative vehicle control. KDAC inhibition was performed by treating cells with a cocktail of 5 mM sodium butyrate (NaB, Sigma 303410), 5 μM trichostatin (TSA, Sigma T8552) and 0.5 mM nicotinamide (NaM) for 2 h at 37°C.

### siRNA treatment

For KAT2A/GCN5 and KAT2B/PCAF knockdowns, U2OS cells at 30% confluence were transfected using DharmaFECT (Dharmacon) according to the manufacturer’s instructions, with 50 pmol each of ON-Targetplus SMARTpools siRNAs targeting KAT2A (Dharmacon L-009722-02-0005) and KAT2B (Dharmacon L-005055-00-0005) in serum-free DMEM. As a negative control, 100 pmol of ON-TARGETplus non-targeting pool siRNAs (Dharmacon D001810-10-05) were used. The serum-free medium was replaced with DMEM supplemented with 10% FBS at 24 h after transfection, and after further 48 h incubation, the cells were collected by trypsinisation (total of 72 h siRNA exposure).

### Fluorescence recovery after photobleaching (FRAP)

For FRAP experiments, cells were plated into CELLview cell culture dishes (Greiner Bio-One) and analysed in phenol red-free Leibovitz’s L15 medium (Gibco). FRAP experiments were performed on a spinning-disk confocal microscope (Ultra-View Vox, Perkin Elmer) mounted on an IX81 Olympus microscope with an Olympus 60x 1.4 oil PlanApo objective, in a controlled chamber at 37°C and 5% CO_2_ (TOKAI HIT stage top incubator). The fluorescence signal was detected using an EMCCD camera (ImagEM, Hamamatsu C9100-13). Cells were bleached in the GFP channel at maximum laser power with a single pulse for 20 ms, within a square region of interest of 5 µm^2^. After bleaching, GFP fluorescence recovery was monitored within the bleached area every second for 40 s. FRAP parameters were controlled using Volocity software 6.0 (QuorumTechnologies). FRAP data were fitted and normalised using the FRAP plugins in ImageJ/Fiji (Cardona et al, 2012). From the FRAP curve fitting, half-time recovery time values after photobleaching (t_1/2_) were extracted and plotted in Prism 7.02, in which statistical analyses were performed. Normalised fluorescence intensity after photobleaching was calculated by dividing the fluorescence intensity at the bleached area by the whole cellular fluorescence intensity.

### Protein purification

FLAG-KAT2A, FLAG-KAT2A catalytic mutant, and FLAG-KAT2B proteins were purified as described previously (Fournier et al, 2016). GST-PALB2 full-length and fragments were purified from 1 L of ArcticExpress cells (Agilent Technologies), grown at 37°C in LB broth medium containing 50 µg/mL ampicillin and 25 µg/mL gentamycin. Protein expression was induced by 0.1 mM IPTG exposure for 24 h at 13°C. Cells were collected by centrifugation for 15 min at 1,400 × *g* at 4°C and washed with ice-cold phosphate-buffered saline (PBS). Cells lysis was performed by 30 min incubation on ice in 15 mL of ice-cold extraction buffer (50 mM Tris-HCl pH 8.0, 150 mM KCl, 1 mM EDTA, 2 mM DTT, 10% glycerol, and protease inhibitor cocktail (PIC, Sigma P2714) supplemented with 2 mg/mL Lysozyme (Sigma) and 0.2% Triton X-100), followed by sonication. Cell lysates were collected after 45 min centrifugation at 35,000 × *g*, 4°C. GST-fusion proteins were pull-down with glutathione Sepharose 4B beads (GE Healthcare), pre-washed with ice-cold PBS. After overnight incubation at 4°C on a rotating wheel, beads were washed three times with ice-cold extraction buffer, three times with 5 mL of ice-cold ATP-Mg buffer (50 mM Tris-HCl pH 7.5, 500 mM KCl, 2 mM DTT, 20 mM MgCl_2_, 5 mM ATP, and 10% glycerol) and three times with ice-cold equilibration buffer (50 mM Tris-HCl pH 8.8, 150 mM KCl, 2 mM DTT, and 10% glycerol). Proteins were eluted from beads in ice-cold elution buffer (50 mM Tris-HCl pH 8.8, 150 mM KCl, 2 mM DTT, 0.1% Triton X-100, 25 mM L-glutathione, and 10% glycerol).

For GFP-ChAM purification for mass spectrometry analysis, HEK293T cells (3 × 10^7^ cells) transiently expressing GFP-ChAM were collected by centrifugation for 5 min at 500 × *g*, 4°C and washed once with ice-cold PBS. Cells were further resuspended in 5 mL ice-cold sucrose buffer (10 mM Tris-HCl pH 8.0, 20 mM KCl, 250 mM sucrose, 2.5 mM MgCl_2_, 10 mM Benz-HCl and PIC). After addition of 10% Triton X-100 (Sigma) to a final concentration of 0.3%, the cell suspension was vortexed four times for 10 s with 1 min intervals. The intact nuclei were collected by centrifugation for 5 min at 500 × *g*, 4°C, and the supernatant was discarded. The nuclear pellet was washed once with ice-cold sucrose buffer and resuspended in ice-cold NETN250 buffer (50 mM Tris-HCl pH 8.0, 250 mM NaCl, 2 mM EDTA, 0.5% NP-40, 10 mM Benz-HCl and PIC). After 30 min incubation on ice, the chromatin fraction was collected by centrifugation for 5 min at 500 × *g*, 4°C, washed once with 5 mL ice-cold NETN250 buffer and lysed for 15 min at room temperature (RT) in ice-cold NETN250 buffer supplemented 5 mM MgCl_2_ and 125 U/mL benzonase (Novagen 71206-3). After addition of EDTA and EGTA to respective final concentrations of 5 mM and 2 mM to inactivate the benzonase and centrifugation for 30 min at 16,100 × *g*, 4°C, the supernatant was collected as the chromatin-enriched fraction. GFP-ChAM were pull-down using 15 µL GFP-Trap Agarose (Chromotek), pre-washed three times with ice-cold NETN250 buffer and blocked for 3 h at 4°C on a rotating wheel with 500 µL ice-cold NETN250 buffer supplemented with 2 mg/mL bovine serum albumin (BSA, Sigma). After 3 h protein binding at 4 °C on a rotating wheel, the GFP-Trap beads were collected by centrifugation for 5 min at 1,000 × *g*, 4°C and washed four times with ice-cold NETN250 buffer.

For the analysis of ChAM acetylation upon DNA damage, a GFP-ChAM fusion was transiently expressed from pDEST53-GFP-ChAM for 24 h in HEK293T cells. Whole-cell extracts (WCE) were prepared from ∼1.5 × 10^7^ cells resuspended in NETN150 buffer (50 mM Tris-HCl pH 8.0, 150 mM NaCl, 2 mM EDTA and 0.5% NP-40 alternative (NP-40 hereafter) (Millipore 492018)) supplemented with 1 mM DTT, PIC, lysine deacetylase inhibitor (5 mM NaB), 1 mM MgCl_2_ and 125 U/mL benzonase. After 30 min incubation on ice, cell debris was removed by 30 min centrifugation at 4°C, and the supernatant was collected as WCE. WCE was then incubated with 15 µl of GFP-Trap Agarose for GFP-pull down. After 1 h protein binding at 4°C on a rotating wheel, GFP-Trap beads were collected by 5 min centrifugation at 500 × *g* at 4°C and washed three times with NET150 buffer (50 mM Tris-HCl pH 8.0, 150 mM NaCl and 2 mM EDTA) supplemented with 0.1% NP-40, 1 mM DTT, PIC, 5 mM NaB and 1 mM MgCl_2_. Proteins were eluted from beads by heating at 85°C for 10 min in Laemmli buffer supplemented with 10 mM DTT. The proteins were separated by SDS-PAGE and analysed by western blot.

For the nucleosome pull-down assays, GFP-ChAM variants were affinity-purified from HEK293T cells (3 × 10^7^ cells) following transient expression. After collecting cells by centrifugation for 5 min at 500 × *g*, 4°C, the cell pellet was washed twice with ice-cold PBS and resuspended in ice-cold NETN150 buffer supplemented with 10 mM benzamidine hydrochloride (Benz-HCl) and PIC. After 30 min incubation on ice, the chromatin was pelleted by centrifugation for 5 min at 500 × *g*, 4°C, and the supernatant was collected as NETN150 soluble fraction and centrifuged for 30 min at 16,100 × *g*, 4°C to remove cell debris and insoluble material. For each sample, 10 µL of GFP-Trap Agarose were washed three times with 500 µL ice-cold NETN150 buffer. NETN150 soluble proteins (2.5 mg) in a total volume of 1 mL ice-cold NETN150 buffer were incubated with the GFP-Trap beads to perform a GFP pull-down. After 2 h incubation at 4°C on a rotating wheel, the GFP-Trap beads were collected by centrifugation for 5 min at 500 × *g*, 4°C and washed four times with ice-cold NETN150 buffer. Human nucleosomes were partially purified from HEK293T cells (4 × 10^7^ cells), collected by centrifugation for 5 min at 500 × *g*, 4°C, washed twice with ice-cold PBS and lysed in ice-cold NETN150 buffer supplemented with 10 mM Benz-HCl and PIC. After 30 min of incubation on ice, the chromatin was pelleted by centrifugation for 5 min at 500 × *g*, 4°C, washed once with ice-cold NETN150 buffer and digested for 12 min at 37 °C with 50 gel units of micrococcal nuclease (NEB) per milligram of DNA in NETN150 buffer containing 5 mM CaCl_2_, using 200 µL buffer per mg of DNA. The reaction was stopped with 5 mM EGTA and the nucleosomes suspension cleared by centrifugation for 30 min at 16,100 × *g*, 4°C.

### Acetyltransferase assays

Radioactive ^14^C-acetyltransferase assays on recombinant proteins were performed by incubating purified GST-PALB2 (full length and fragments) or purified RAD51 with purified FLAG-KAT2B in the presence of ^14^C-labeled acetyl-CoA in the reaction buffer (50 mM Tris-HCl pH 8.0, 10% glycerol, 100 mM EDTA, 50 mM KCl, 0.1 M NaB, PIC, and 5 mM DTT) for 1 h at 30°C. The reactions were stopped by addition of Laemmli buffer containing 10% beta-mercaptoethanol, boiled for 5 min, resolved by SDS-PAGE and stained using Coomassie blue to reveal overall protein distribution. The acrylamide gel was then dried and exposed to phosphorimager to reveal ^14^C-labeled proteins. Non-radioactive acetyltransferase assays were performed as described above using cold acetyl-CoA instead. After 1 h incubation at 30°C, the reactions were stopped by addition of Laemmli buffer containing 10 mM DTT, boiled for 5 min, resolved by SDS-PAGE, and after Ponceau S staining of the membrane to reveal overall protein distribution, analysed by western blot using anti-acetyl lysine antibody. Acetyltransferase assays performed for mass spectrometry analyses were performed as previously described (Fournier et al, 2016)

### Nucleosome pull-down assay

Nucleosome pull-down assays were performed by mixing 250 µg of partially purified nucleosomes and GFP-ChAM variants immobilised on GFP Trap beads in NETN150 buffer supplemented with 2 mg/mL BSA, followed by 30 min incubation at RT, then 1.5 h incubation at 4 °C, on a rotating wheel. GFP-Trap beads were further washed four times with NETN150 buffer, and samples were analysed by SDS-PAGE and western blot.

### DNA binding assays

For ChAM peptides binding to DNA, proteins or peptides were incubated in NDTN150 buffer (50 mM Tris HCl pH 7.5, 150 mM NaCl, 0.5% NP-40 and 2 mM DTT) with 300 ng pBluescript plasmid DNA for 1 h at room temperature (RT). Reactions were loaded onto a 1% agarose Tris-Acetate-EDTA (TAE) gel and run at 100 V for 60 min. The gel was stained for 1 h at RT with 0.5 µg/mL ethidium bromide in Tris-Borate-EDTA (TBE) buffer. Binding of ChAM peptide to histone H3 was performed by incubating for 1 h at 4°C biotinylated ChAM peptides and histone H3 protein in NDTN250 buffer (50 mM Tris HCl pH 7.5, 250 mM NaCl, 0.5% NP-40, and 2 mM DTT) supplemented with 5 mg/mL BSA. Dynabeads MyOne T1 blocked with BSA in NDTN250 buffer were added to the peptide-Histone H3 mixture and incubated for 1 h at 4°C. Immobilised complexes were washed three times with NDTN250 buffer and proteins were visualised by western blotting using the indicated antibodies.

### Immunofluorescence microscopy

For γ-H2AX foci analysis, cells were grown on coverslips and washed with PBS before pre-extraction with 0.1% Triton in PBS for 30 s at RT. Cells were then fixed twice with 4% PFA in PBS, first for 10 min on ice and then for 10 min at RT. After permeabilisation in 0.5% Triton X-100 in PBS for 10 min at RT, cells were blocked with 5% BSA in PBS supplemented with 0.1% Tween 20 solution (PBS-T-0.1) and incubated with anti-γ-H2AX antibody for 3 h at RT. After washing with PBS-T-0.1 for 5 min at RT, cells were incubated with secondary antibodies coupled with a fluorophore, washed with PBS-T-0.1 for 5 min at RT, and mounted on slides using a DAPI-containing solution. For RAD51 foci analysis, cells were grown on coverslips and washed twice with PBS before fixation with 4% PFA in PBS for 15 min at RT. After washing twice with PBS, cells were permeabilised with 0.5% Triton X-100 in PBS for 5 min at RT and blocked with 3% BSA in PBS supplemented with 0.1% Triton X-100 before incubation with anti-RAD51 antibody for 2 h at RT. After washing three times with PBS supplemented with 0.5% Tween 20 (PBS-T-0.5), cells were incubated with secondary antibody coupled with a fluorophore, washed three times with PBS-T-0.5 and mounted on slides using a DAPI-containing solution. Cells were analysed on a spinning-disk confocal microscope (Ultra-View Vox, Perkin Elmer) mounted on an IX81 Olympus microscope, with a 40x 1.3 oil UPlan FL objective. The fluorescence signal was detected using an EMCCD camera (ImagEM, Hamamatsu C9100-13). Images were processed in Image J (https://imagej.nih.gov/ij/) (Schneider et al, 2012).

### Sequence alignment analysis

Sequences of PALB2 orthologues from 12 species were retrieved from the Ensembl (http://www.ensembl.org) and NCBI (https://www.ncbi.nlm.nih.gov) and aligned using MUSCLE (http://www.ebi.ac.uk/Tools/msa/muscle/).

## Results

### PALB2 is acetylated within a 7K-patch in its ChAM domain

Our previous shotgun mass spectrometry (MS) study of the endogenous KAT2A/B-acetylome in HeLa cells identified a number of lysine residues within the central region of PALB2 as acetylation acceptors (Fournier et al, 2016) (**Fig. 1A**, highlighted in blue). To confirm the direct acetylation of PALB2 by KAT2A/B, we performed *in vitro* acetylation assays using either recombinant full-length PALB2 or a series of PALB2 fragments (**Fig, 1A**) with purified KAT2A or KAT2B.

**Figure 1.**
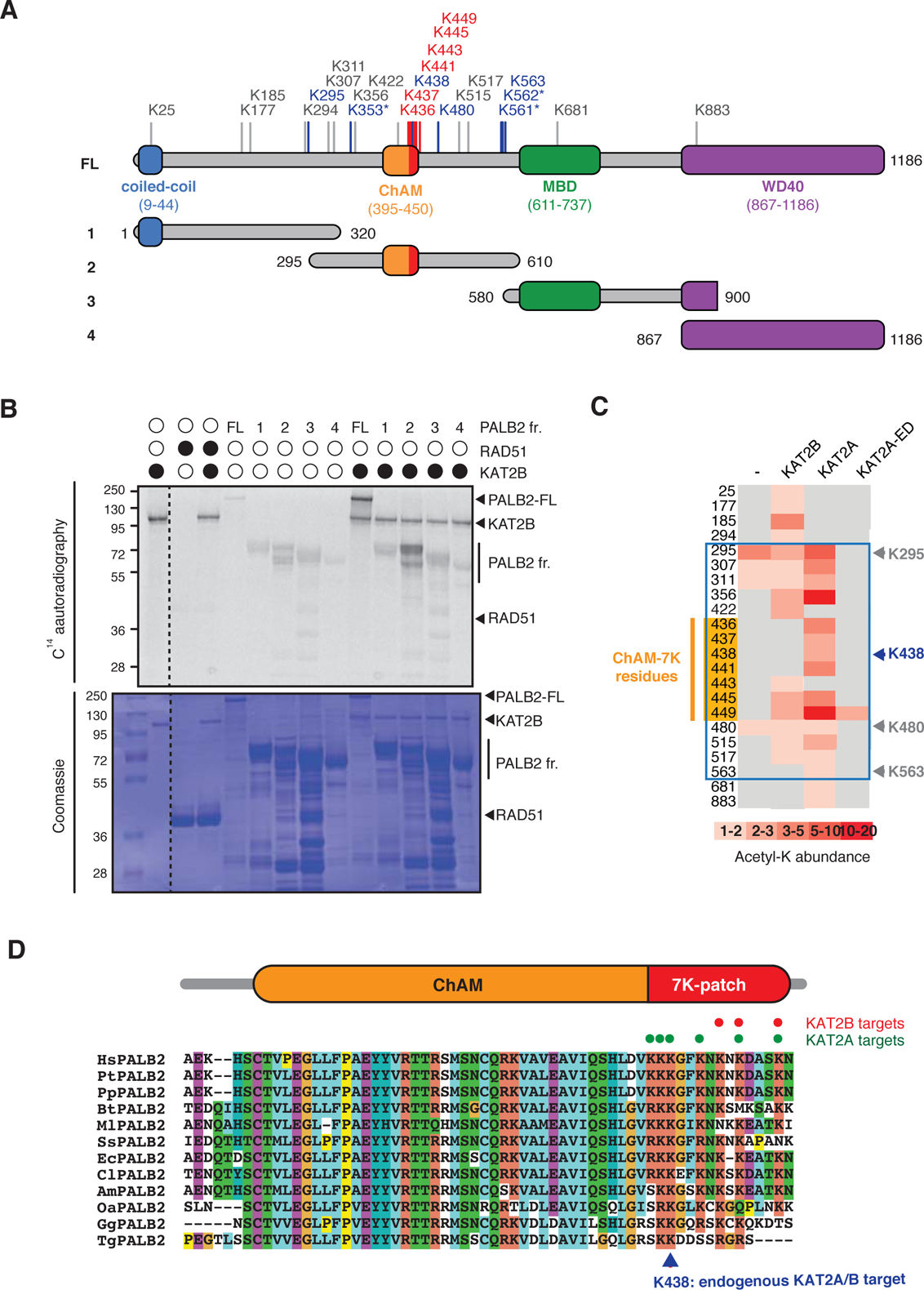
KAT2A/B acetylates PALB2 within a 7K-patch in its ChAM domain. ***A.*** PALB2 lysine residues identified as acceptors of KAT2A/B-dependent acetylation *in vivo* and *in vitro* by tandem MS analyses are shown in blue and black, respectively. The acetylated lysine residues within the ChAM are highlighted in red. Full-length PALB2 (FL, 200kDa) and fragments 1 to 4 used for *in vitro* acetylation assays are also shown. 1: PALB2 1-320 (61.2kDa), 2: 295-610 (60.6kDa), 3: 580-900 (61.2 kDa) and 4: 867-1186 (61.1kDa). ***B*.** PALB2 FL and fragments 1 to 4 were acetylated *in vitro* by KAT2B with ^14^C-labelled acetyl-CoA, and following SDS-PAGE, acetylated proteins and total proteins were detected by ^14^C-autoradiography and Coomassie blue staining, respectively. RAD51 was not identified as a target of KAT2A/B *in vivo* (Fournier et al, 2016) and was used as a negative control. ***C*.** A heat map of acetylated lysine residues, as identified by quantitative MS analysis of *in vitro* acetylated PALB2 FL or fragment 2 by KAT2B, KAT2A or a catalytically inactive KAT2A (KAT2A-ED). The abundance of each acetyl-lysine was evaluated as previously described (Fournier et al, 2016). ***D.*** ChAM protein sequences from twelve PALB2 orthologues were aligned using MUSCLE and visualised using ClustalW default colour-coding. Lysine residues acetylated by KAT2A and KAT2B *in vitro* are respectively highlighted by red and green circles. Hs (*Homo sapiens*, human), Pt (*Pan troglodytes*, chimpanzee), Pp (*Pan paniscus*, bonobo), Bt (*Bos taurus*, cow), Ml (*Myotis lucifugus*, little brown bat), Ss (*Sus scrofa*, wild boar), Ec (*Equus caballus*, horse), Cl (*Canis lupus familiaris*, dog), Am (*Ailuropoda melanoleuca*, giant panda), Oa (*Ovis aries*, sheep), Gg (*Gallus gallus*, red junglefowl), Tg (*Taeniopygia guttata*, zebra finch).

Lysine acetylation of full-length PALB2 and fragment 2, which encompasses the DNA/chromatin binding region of PALB2 (residues 295-610), was clearly visible by ^14^C-autoradiography (**Fig. 1B**) or western blot (WB) against acetylated lysine (pan-AcK) following *in vitro* acetylation with KAT2A or KAT2B, but not with a catalytically inactive mutant of KAT2A (KAT2A-ED) (**Fig. S1*A* and *B***). Our MS analyses of these products identified acetylated lysine residues in the PALB2 chromatin association motif (ChAM) within the C-terminal cluster of seven lysine residues composed of K436, K437, K438, K441, K443, K445, and K449 (**Fig. 1C**, highlighted in orange), denoted the 7K-patch, which is common to PALB2 orthologs (Bleuyard et al, 2012) (**Fig. 1D**). A cluster of lysine residues is often found in KAT2A/B substrates (Fournier et al, 2016), corroborating our finding that the ChAM is favorably targeted for KAT2A/B-mediated acetylation.

### The 7K-patch promotes ChAM nucleosome association

To evaluate the impact of the 7K-patch on ChAM chromatin association property, we generated a series of GFP-tagged ChAM truncations (**Fig. 2A**). Consistent with our previous observations, the full-length ChAM fragment (PALB2 residues 395-450) was highly enriched in the chromatin fraction in HEK293T cells (**Figs. 2*A*, *B* and S2*A***). Further analysis of ChAM truncations revealed that the region containing the 7K-patch was essential for its chromatin association (**Fig. 2A** and **B**). Considering the possibility that the deletion of 7K-patch might lead to aberrant subcellular localisation and have contributed to the reduced chromatin enrichment, we also affinity purified the corresponding fragments from the soluble fraction of HEK293T, and assessed their interaction with separately prepared nucleosomes (**Fig. S2*A***). Again, the pull-down assay showed that deletion of the 7K-patch impaired ChAM interaction with nucleosomes (**Fig. 2A** and **C**).

**Figure 2.**
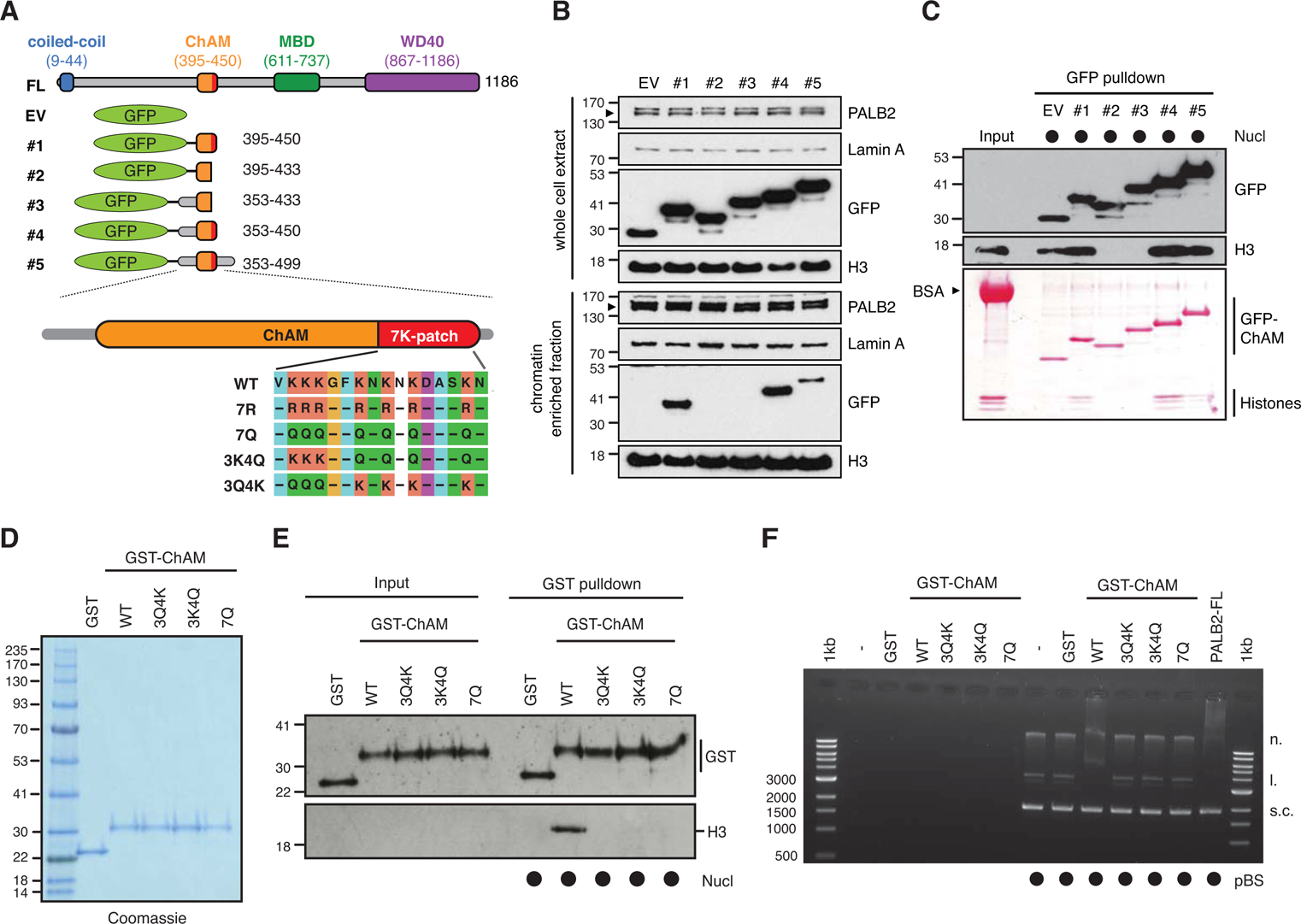
The 7K-patch promotes ChAM nucleosome association. ***A*.** Diagram showing the full-length (FL) PALB2, GFP-ChAM fragments #1 to #5 and GST-ChAM variants used in this study. ChAM C-terminal lysine-rich patch (7K-patch, residues 436-449) is highlighted in red. ***B.*** Western blotting assessing the chromatin enrichment of GFP-ChAM fragments transiently expressed in HEK293T cells. Empty GFP vector (EV) was used as a negative control. Whole cell extract was also prepared to compare GFP-ChAM fragments expression levels. Lamin A and histone H3 were used as controls for the cellular fractionation. ***C***. *In vitro* nucleosome binding assay using GFP and GFP-ChAM fragments. Partially purified human nucleosomes captured by GFP and GFP-ChAM fragments immobilised on beads were detected by anti-histone H3 western blot and Ponceau S staining. ***D.*** Coomassie blue staining of GST and GST-ChAM variants purified from *E.coli* and separated by SDS-PAGE. ***E.*** *In vitro* binding assays using immobilised GST-ChAM variants and purified HeLa mononucleosomes. GST was used as a negative control. The nucleosome binding efficiency was assessed by anti-histone H3 western blot. ***F.*** Electrophoretic mobility shift assay (EMSA) using pBluescript plasmid DNA (pBS) and purified GST-ChAM variants. Three DNA conformations are indicated to the right: supercoiled (s.c.), linear (l.), and nicked (n.).

To further assess the direct role of the 7K-patch acetylation on nucleosome binding, we purified recombinant GST-fusions of ChAM WT and mutants harboring glutamine substitutions at all seven lysine residues (7Q), three conserved lysine residues, K436, K437 and K438 (3Q4K), or the four remaining lysine residues, K441, K443, K445, and K449 (3K4Q) within the 7K-patch (**Fig. 2A** and **D**). K to Q substitutions nullify the positive charge of lysine and are therefore commonly considered to mimic acetyl-lysine. Significantly, *in vitro* nucleosome pull-down assays showed that all variants with Q substitutions abolished ChAM interaction with nucleosomes (**Fig. 2E**). As anticipated from the change in electrostatic charge, all ChAM variants with K to Q substitutions also exhibited an impaired affinity for DNA, as determined by native electrophoretic mobility shift assay (EMSA) (**Fig. 2F**). Collectively, these results confirmed that Q substitutions of the 7K-patch perturbed the ability of ChAM to bind both nucleosomes and naked DNA; this variant is hence considered to behave as a ChAM null.

### Acetylation within the 7K-patch enhances ChAM nucleosome association

KAT2A/B associate with chromatin and facilitate transcription by acetylating histones (Nagy et al, 2010). Consistently, we found that the level of acetylation in the GFP-ChAM fragment affinity purified from the chromatin-enriched fraction was notably higher than that from the cytoplasmic or the nuclear soluble fraction (**Fig. S2*A* and *B***). Our MS analysis of the chromatin-associated GFP-ChAM fragment identified acetylation of all seven lysine residues within the 7K-patch (**Fig. 3A**, marked with arrows). This observation appeared to be at odds with our observation that acetyl-mimetic Q substitutions disabled ChAM binding to nucleosomes (**Fig. 2D** and **E**). Since the side chain of glutamine is significantly shorter than the corresponding acetyl-lysine side chain (**Fig. S2*C***), we questioned whether the K to Q substitutions accurately mimic the properties of acetylated ChAM. Hence, synthetic peptides corresponding the minimal ChAM (PALB2 residues 395-450) containing acetyl-lysine at the evolutionarily conserved K436, K437 and/or K438, were generated and their biochemical properties were characterised. Nucleosome pull-down assays revealed a modest increase in nucleosome binding with acetylated ChAM peptides compared to their non-acetylated counterpart (**Fig. 3B**). Importantly, when salmon sperm DNA was supplemented in excess to outcompete the electrostatic ChAM interaction with DNA, only acetylated ChAM, but not non-acetylated ChAM, maintained its ability to bind nucleosomes. This effect was most evident with the ChAM peptide containing acetyl-K438, which was detected in our original MS study of the endogenous HeLa KAT2A/B-acetylome (Fournier et al, 2016). As anticipated, lysine acetylation, which neutralises the positive charge on the lysine side chain, conferred reduced affinity for negatively charged DNA (**Fig. 3C** and ***D***). Together, these results demonstrate that acetylation within the 7K-patch enhances ChAM nucleosome binding while limiting its non-specific affinity for DNA.

**Figure 3.**
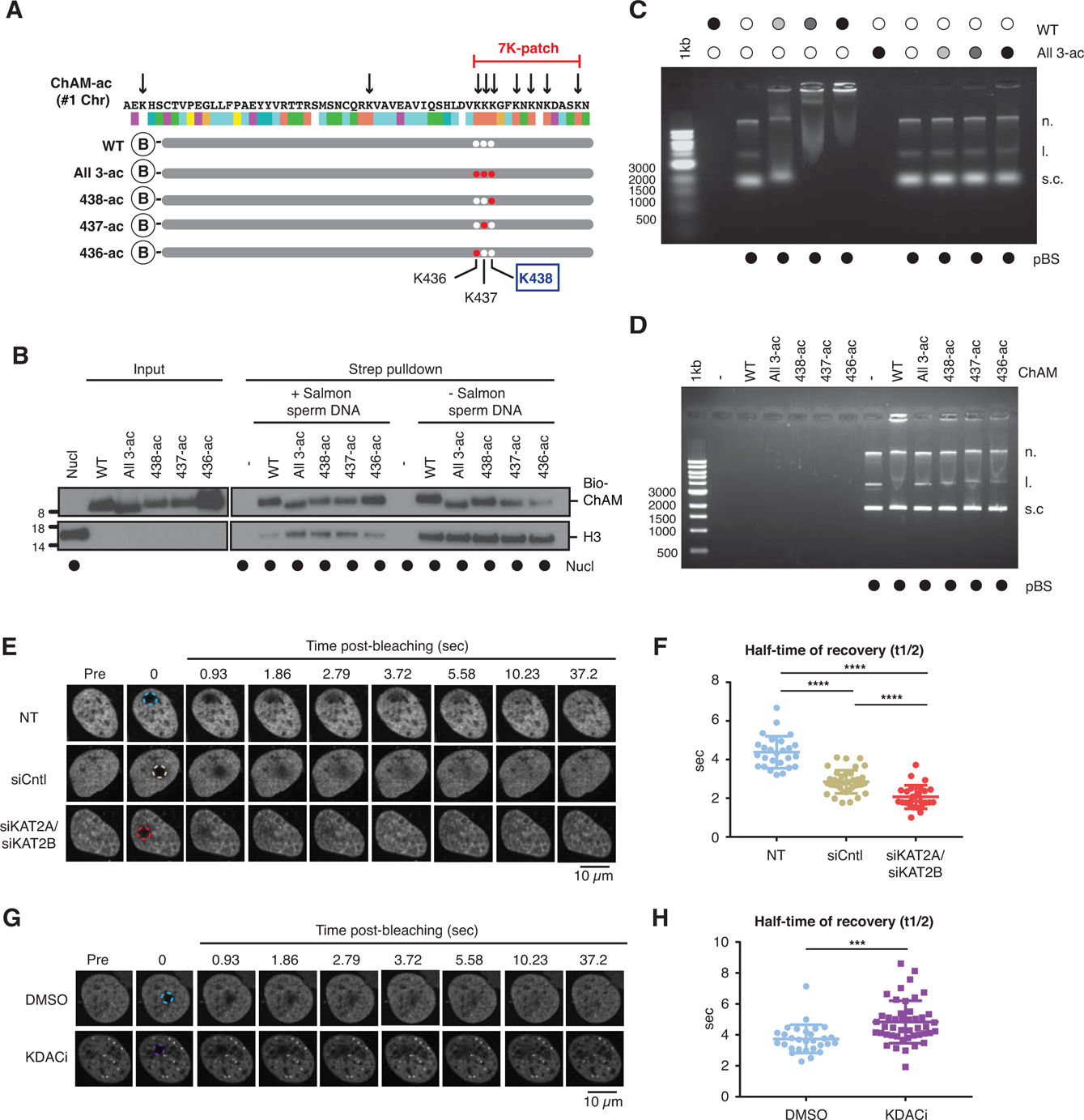
Acetylation within the 7K-patch enhances ChAM nucleosome association. ***A*.** *In vivo* acetylated lysine residues in chromatin-associated ChAM, detected by tandem MS analysis, are highlighted by arrows. Synthetic biotin-ChAM peptides non-acetylated (WT) or acetylated at single lysine residues 436 (436-ac), 437 (437-ac), and 438 (438-ac), or all three lysine residues (All 3-ac) are shown. ***B.*** *In vitro* nucleosome binding assays for the synthetic biotin-ChAM peptides. After incubation with purified HeLa polynucleosomes in the absence or presence of 5 µg salmon sperm DNA, biotin-ChAM peptides were pulled-down by streptavidin beads (Strep pulldown), and associated nucleosome was detected by anti-histone H3 western blot. ***C.*** EMSA using 300 ng pBluescript plasmid DNA and increasing amount of indicated biotin-ChAM peptides, i.e. 2.97 µM (light gray circle), 5.94 µM (dark gray circle) and 29.7 µM (black circle). Three DNA conformations are indicated: supercoiled (s.c.), linear (l.), and nicked (n.). ***D.*** As in *C*, except using the indicated biotin-ChAM peptides. ***E-F*.** FRAP assay of FE-PALB2 in U2OS cells in which KAT2A and KAT2B were depleted by siRNA (siKAT2A/siKAT2B). Non-treated (NT) and negative control siRNA (siCntl) treated cells were used as controls. ***G-H***. As in *E-F*, except cells were treated with a cocktail of lysine deacetylase inhibitors (KDACi, 5 mM sodium butyrate, 5 µM trichostatin A and 5 mM nicotinamide) or DMSO as a control. For each condition, representative images of live cells before bleaching (pre-bleaching) and during a 37.5 s recovery period (post-bleaching) are shown together with half-recovery time plots for at least 20 cells. Dashed circles indicate bleached areas. Statistical analyses were performed using GraphPad Prism 7.02 and *p*-values are for the unpaired Student’s *t*-test (** *p* < 0.0035, *** *p* < 0.0006, and **** *p* < 0.0001).

### PALB2 mobility increases upon deacetylation

Encouraged by the *in vitro* results using acetylated ChAM peptides, we further assessed whether lysine acetylation might equally facilitate PALB2 chromatin association *in vivo*. To evaluate the impact of native PALB2 acetylation, we down-regulated KAT2A/B and assessed PALB2 mobility as a surrogate measure of its chromatin association. To this end, a tandem FLAG- and EGFP-tagged full-length wild-type (WT) PALB2 (FE-PALB2) was conditionally expressed in a U2OS cell line in which endogenous *PALB2* can be down-regulated by a doxycycline-inducible short hairpin RNA (U2OS-shPALB2) **(Fig. S3*A*)**, and its mobility was assessed using fluorescence recovery after photobleaching (FRAP). This analysis revealed that KAT2A/B siRNA treatment indeed led to an increase in FE-PALB2 diffusion rate (reduced FRAP t_1/2_; **Figs. 3*E*, *F*, S3*B* and *C***), concomitant with reduced levels of global acetylation (Fournier et al, 2016). Conversely, the inhibition of lysine deacetylases (KDAC), which increased the global level of lysine acetylation, decreased the diffusion rate of FE-PALB2 (increased FRAP t_1/2_; **Figs. 3*G*, *H*, S3*D* and *E***). These results support the notion that PALB2 chromatin association is enhanced by KAT2A/B-mediated acetylation events.

### PALB2 ChAM 7K-patch acetylation is required for the protection of genomic DNA during DNA replication

To directly investigate the role of the 7K-patch acetylation *in vivo*, we generated full-length PALB2 variants in which lysine residues within the 7K-patch were substituted by Q or arginine (R). Contrary to the K to Q null-substitutions, which nullifies the positive charge of lysine, K to R substitutions maintain the charge yet are unable to accept acetylation and hence mimic constitutively non-acetyl-lysine (**Fig. S2*C***). These PALB2 variants, named 7Q or 7R, as well as PALB2 WT or an empty vector (EV), were stably expressed as FLAG-fusions at equivalent levels in U2OS-shPALB2 cells (**Figs. 4*A* and S4*A***). Our subcellular fractionation analyses showed a reduced level of PALB2 7Q in the chromatin-enriched fraction compared to PALB2 WT and 7R (**Fig. 4B** and **C**). In line with these observations, our FRAP analysis of the PALB2 variants, conditionally expressed as FE-fusions in U2OS-shPALB2, revealed an increase in PALB2 7Q diffusion kinetics compared to those of PALB2 WT and 7R (**Fig. 4D** and **E**). A PALB2 variant with an internal deletion of ChAM, which abolishes its chromatin association (Bleuyard et al, 2012), similarly exhibited a significant increase in PALB2 diffusion rate (**Fig. S4*B*-*D***). Hence, we concluded that the increase in PALB2 mobility reflected the chromatin-free fraction of PALB2, which is controlled by its ChAM 7K-patch.

**Figure 4.**
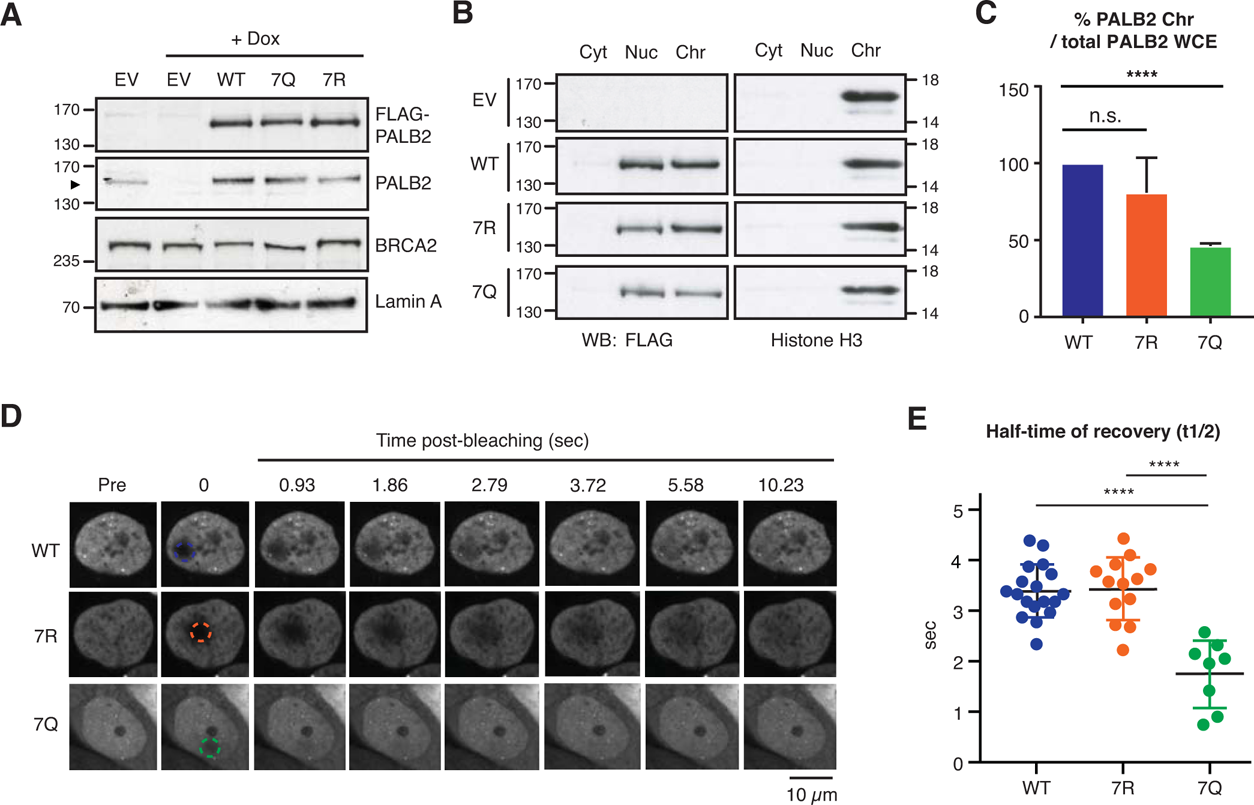
PALB2 ChAM 7K-patch mediate global chromatin association. ***A*.** Western blot analysis of U2OS-shPALB2 cells constitutively expressing FLAG (EV) or FLAG-PALB2 variants. Where indicated, cells were treated with 2 µg/mL doxycycline (+dox) to deplete endogenous PALB2. Lamin A was used as a loading control. ***B-C***. Subcellular distribution of FLAG-PALB2 variants in cytoplasmic (Cyt), nuclear soluble (Nuc) and chromatin-enriched (Chr) fractions upon doxycycline-induced PALB2 depletion. Histone H3 was used as a control for cellular fractionation. Protein levels, detected by western blotting, were quantified by ImageJ. PALB2 levels in the chromatin fraction were normalised against PALB2 levels in whole-cell extracts shown in *A* and expressed as a percentage of the WT in *C*. Statistical analyses were performed using GraphPad Prism 7.02, and *p*-values are for the unpaired Student’s *t*-test (**** *p* < 0.0001, ns = non-specific). ***D-E***. FRAP analysis of FE-PALB2 WT, 7R and 7Q conditionally expressed in U2OS-shPALB2 cells in which endogenous PALB2 was depleted *via* doxycycline exposure. Representative images before bleaching (pre-bleaching) and during the recovery period (post-bleaching) are shown in *D*, where dashed circles indicate bleached areas. Half-recovery time plots for at least 20 cells per cell line are shown in *E*. Statistical analyses were performed using GraphPad Prism 7.02 and *p*-values are for the unpaired Student’s *t*-test (**** *p* < 0.0001).

In light of our previous finding that steady-state PALB2 chromatin association is important for the protection of actively transcribed genes during DNA replication (Bleuyard et al, 2017a), we assessed cell cycle progression of U2OS-shPALB2 cells complemented with either PALB2 WT, 7Q and 7R. To this end, cells were arrested at G1/S boundary by double thymidine block, in the presence of doxycycline throughout (43 h), and were synchronously released into S phase. To our surprise, not only PALB2 7Q but also 7R showed slow progression of S-phase compared to WT complemented cells (**Fig. 5A, B** and ***C***). In line with this observation, PALB2 7Q or 7R-complemented cells displayed increased levels of the well-established DDR marker, serine 139 phosphorylated histone variant H2AX (γ-H2AX), upon doxycycline exposure (**Fig. 5D** and **E**). While PALB2 7R appeared to maintain global chromatin association equivalent to PALB2 WT (**Fig. 4B** and **C**), PALB2 function in gene protection requires its enriched association with a subset of actively transcribed genes (Bleuyard et al, 2017a). Markedly, our ChIP-qPCR revealed reduced enrichment of both PALB2 7Q and 7R at known PALB2-bound loci, namely the *ACTB, TCOF1* and *WEE1* genes (**Fig. S5**). Together, these results suggest that ChAM acetylation promotes PALB2 enrichment at active genes, which in turn protects these regions from DNA damage during DNA replication.

**Figure 5.**
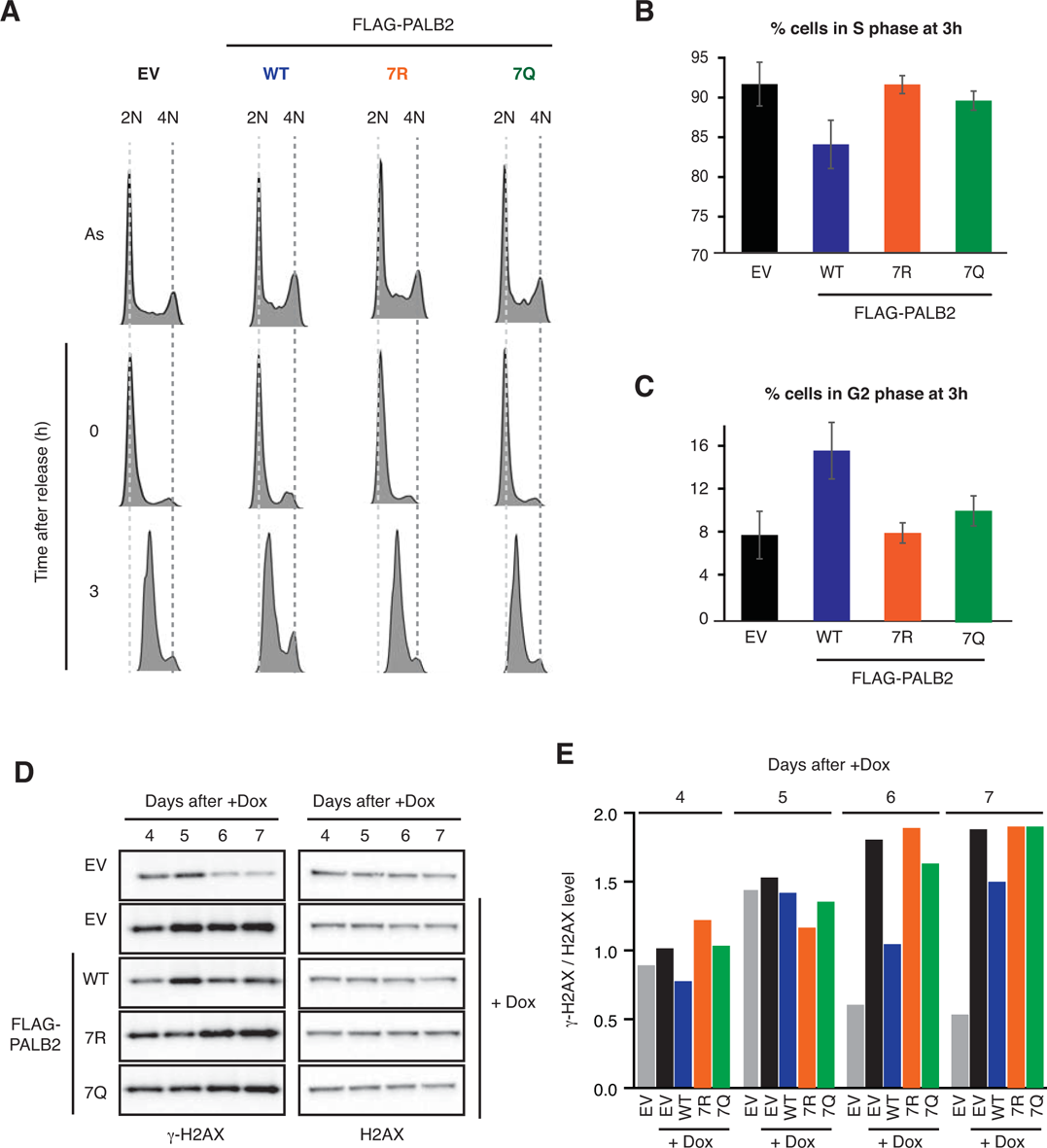
PALB2 ChAM 7K-patch acetylation is required to maintain genome stability during replication. ***A.*** DNA content of U2OS-shPALB2 cell lines stably expressing FLAG (EV) and FLAG-PALB2 variants, as assessed by flow cytometry (FACS) analysis of cells stained with propidium iodide (PI). Samples were prepared under three different conditions: asynchronous (As), synchronised at G1/S by double thymidine block (0 h) and at 3 hour after thymidine release (3 h). ***B-C.*** Histogram showing the percentage of U2OS-shPALB2 EV, FLAG-PALB2 WT, 7Q and 7R cells in S (DNA content between 2N and 4N) and G2 (DNA content 4N) phases of the cell cycle, based on the quantification of FACS profiles shown in *A*. ***D.*** γ-H2AX levels in U2OS-shPALB2 expressing the FLAG (EV) or FLAG-PALB2 WT, 7Q and 7R upon doxycycline (dox)-induced endogenous PALB2 depletion were detected by western blot. Total H2AX was used as a loading control. ***E.*** Quantification of γ-H2AX and H2AX levels shown in *D*, performed by densitometry analysis in ImageJ. Histogram shows the ratios of γ-H2AX versus H2AX levels.

### DNA damage triggers ChAM deacetylation and an increase in PALB2 mobility

So far, our analyses showed that ChAM acetylation promotes PALB2 association with nucleosomes/chromatin (**Fig. 6A**), whereas PALB2 dissociates from active genes upon CPT treatment (Bleuyard et al, 2017b). These observations prompted us to further investigate whether ChAM acetylation is dynamically controlled during the DDR. Significantly, rapid deacetylation of ChAM, which was expressed exogenously in HEK293 cells, was detectable within 15 min following ionizing radiation (IR) and persisted for at least 2 h (**Fig. 6B** and ***C***). Furthermore, using the FRAP approach, we observed clear increases in diffusion rates of FE-PALB2 following DNA damage induced by IR, MMC or olaparib treatment, where DNA damaging efficiency was confirmed by γ-H2AX staining (**Figs. 6*D*-*G*, S6*A*-*D***). Compared to untreated control conditions, IR, MMC and olaparib treatment decreased FRAP t_1/2_ by 22%, 36%, and 18%, respectively. Together, these results indicate that DNA damage triggers ChAM deacetylation and induces PALB2 mobilisation.

**Figure 6.**
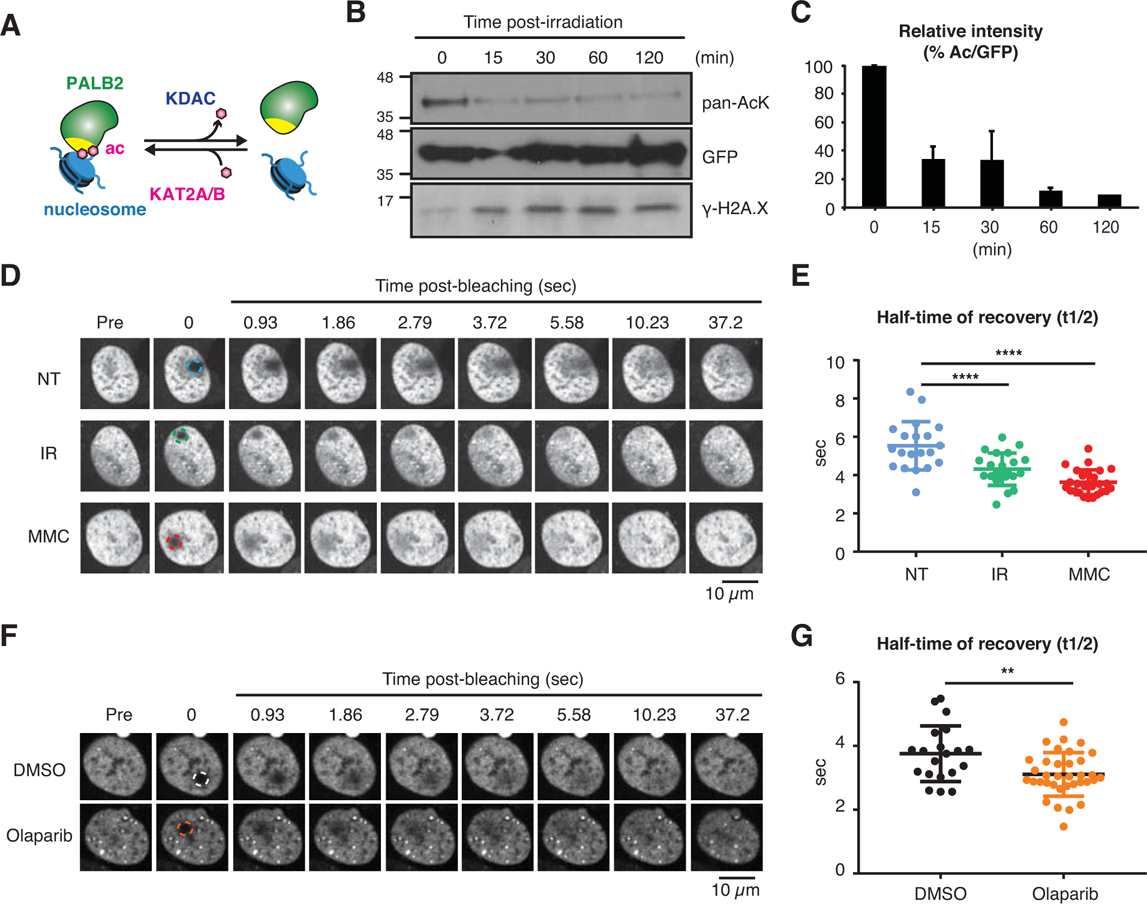
DNA damage triggers ChAM deacetylation and PALB2 mobilization. ***A.*** Depiction of PALB2 acetylation and nucleosome binding, controlled by KAT2A/B and KDACs. ***B*.** HEK293 cells transiently expressing GFP-tagged ChAM were treated with 4 Gy IR, and affinity-purified GFP-ChAM was assessed using anti-GFP and anti-acetyl-lysine (pan-AcK) antibodies. γ-H2AX signal in whole cell lysate was detected to monitor DNA damage. ***C*.** Histogram showing the relative level of ChAM acetylation following irradiation. The levels of acetylated and total GFP-ChAM were quantified by ImageJ, and the ratio of acetyl/total GFP-ChAM was normalised against that of the pre-irradiation sample (time 0). Error bars indicate standard deviation of three independent experiments (n=3). ***D-E*.** The mobility of FE-PALB2 in U2OS-shPALB2 was monitored by FRAP. Cells were examined under non-treated conditions (NT) or treated with 4 Gy IR or 1 mM MMC. ***F-G*.** As in *D-E*, except cells were treated with the PARP inhibitor olaparib or DMSO as a control. For each condition, representative images of live cells before bleaching (pre-bleaching) and during a 37.5 s recovery period (post-bleaching) are shown together with half-recovery time plots for at least 20 cells. Dashed circles indicate bleached areas. Statistical analyses were performed using GraphPad Prism 7.02 and *p*-values are for the unpaired Student’s *t*-test (** *p* < 0.0035, *** *p* < 0.0006, and **** *p* < 0.0001).

### 7K-patch-mediated PALB2 chromatin association is important for HR repair

Given that DNA damage triggers ChAM deacetylation and PALB2 mobilisation, we further evaluated the impact of the 7K-patch in the DDR. As expected, the depletion of endogenous PALB2 conferred impaired RAD51 foci formation both in undamaged and upon MMS or IR exposure (**Fig. S7*A***), which was rescued by PALB2 WT complementation (**Fig. 7A** and **B**). Markedly, cells complemented with PALB2 7R were similarly able to restore MMS-induced RAD51 foci formation, but the 7Q variant failed to do so (**Fig. 7A** and **B**). Consistent with this observation, PALB2 7Q-complemented cells also exhibited increased olaparib sensitivity compared to those complemented with PALB2 WT or 7R (**Fig. 7C**). Given that increased sensitivity to PARP inhibition is a hallmark of HR deficiency, this finding suggested that the 7Q substitutions conferred HR defects. It is noteworthy that all PALB2 variants exhibited equivalent levels of interaction with BRCA2 or RAD51 (**Fig. S7*B***) and were recruited to sites of DNA damage with similar efficiency (**Fig. S7*C* and *D***). These findings indicated that lysine residues within the ChAM 7K-patch are indispensable for PALB2 function in HR, but that this mechanism is independent of PALB2 binding to BRCA2 or RAD51 or its recruitment to sites of DNA damage.

**Figure 7.**
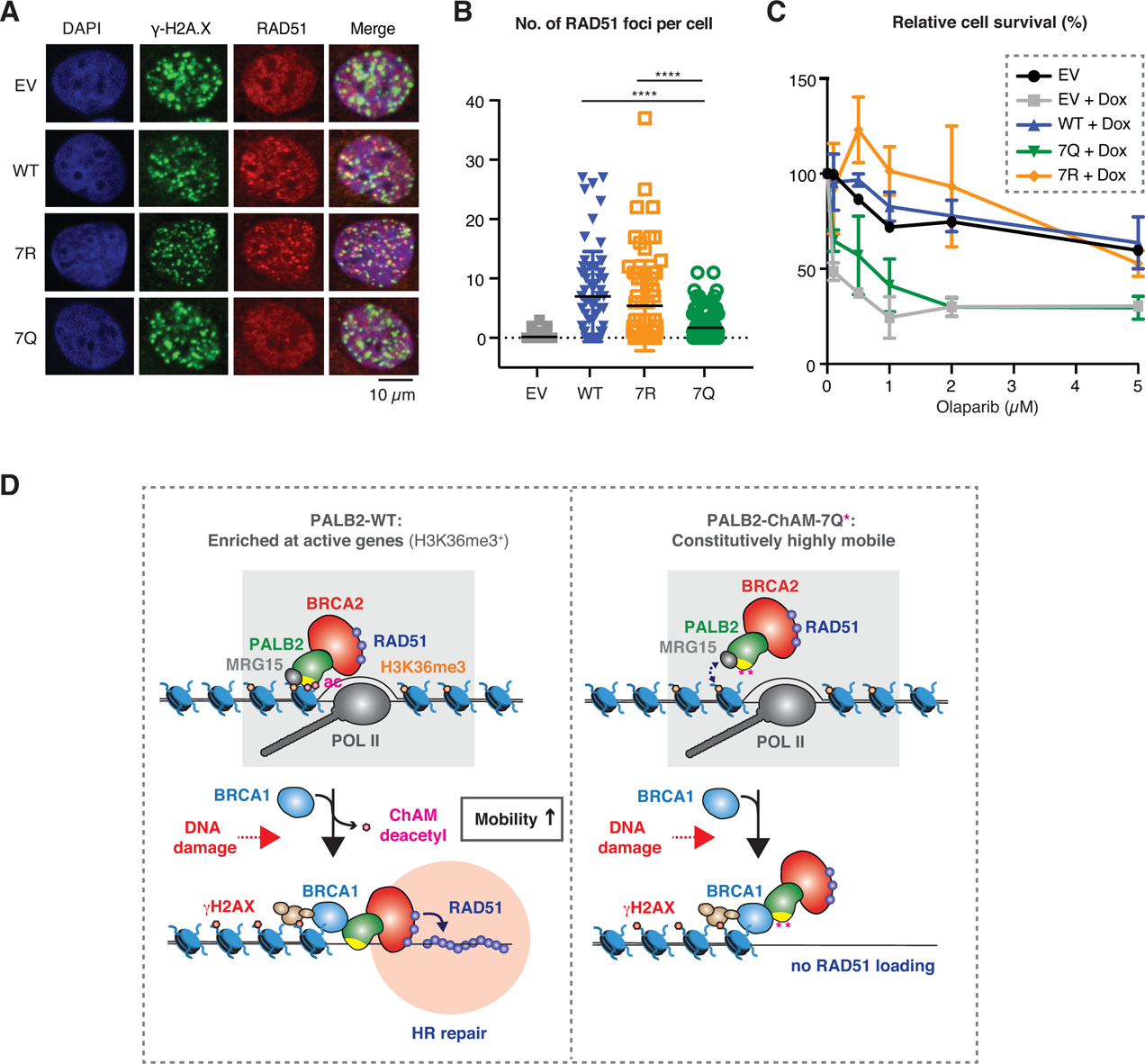
7K-patch-mediated PALB2 chromatin association is important for HR repair. ***A.*** Representative images of IR-induced RAD51 foci in U2OS-shPALB2 cells stably expressing FLAG (EV) and FLAG-PALB2 variants, following doxycycline-induced endogenous PALB2 depletion and 4 Gy IR treatment. DAPI was used for nuclear staining and γ-H2AX as a DNA damage marker. ***B.*** Quantification of the number of RAD51 foci per cell, shown in *A* (n ≥ 62). Statistical analyses were performed using GraphPad Prism 7.02 and *p*-values are for the unpaired Student’s *t-*test (**** *p* < 0.0001). ***C.*** Cell survival upon olaparib treatment of U2OS-shPALB2 cells stably expressing FLAG (EV) or FLAG-PALB2 variants, as assessed by WST-1 assay after five days of olaparib exposure. Where indicated, endogenous PALB2 was depleted by doxycycline exposure. ***D***. Model for PALB2 acetylation function in the maintenance of genome stability. *Left*: MRG15 and KAT2A/B-mediated ChAM acetylation jointly promote PALB2 enrichment at undamaged transcriptionally active chromatin. DNA damage triggers ChAM deacetylation and releases PALB2 from chromatin. This allows PALB2 to interact with BRCA1, which in turn recruits the entire HR complex to sites of DNA damage. ChAM binding to naked DNA through the 7K-patch may promote RAD51 loading and HR repair. *Right*: The 7K-patch null mutant with K to Q substitutions is more freely diffusible, and although it can be recruited to sites of DNA damage, it fails to engage with damaged DNA, leading to defective HR.

## Discussion

In this study, we have demonstrated that reversible lysine acetylation controls the mode of PALB2 chromatin association: the 7K-patch in the C-terminal part of ChAM was acetylated by KAT2A/B in undamaged cells, and was important for normal damage-induced RAD51 recruitment and olaparib resistance. Crucially, ChAM acetylation enhances PALB2 binding to nucleosomes and limits non-specific interactions with DNA. DNA damage, however, triggers PALB2 deacetylation and increases the overall mobility of PALB2. Considering all our findings collectively, we propose the model shown in **Fig. 7D**: KAT2A/B-mediated ChAM acetylation, jointly with MRG15, enforces PALB2 enrichment at undamaged transcriptionally active chromatin and hence protects these regions during DNA replication. DNA damage triggers ChAM deacetylation and a concomitant reduction in PALB2-MRG15 interaction (Bleuyard et al, 2017b), which releases PALB2 from chromatin and increases its mobility. In this way, PALB2 may become available to form functional HR complexes with BRCA1, which is otherwise less stably associated with PALB2 (Bleuyard et al, 2017b). Following recruitment of the entire HR complex to sites of DNA damage, deacetylated ChAM binding to exposed naked DNA may allow appropriate engagement of this complex with damaged DNA and hence promote RAD51 loading and HR repair.

The increase in PALB2 mobility during the DDR shown in this study is reminiscent of previous reports showing increased RAD51 mobility upon replication stress (Srivastava et al, 2009; Yu et al, 2003). Yu *et al*. showed that a fraction of RAD51, which is in complex with BRCA2, is immobile in unchallenged cells, but becomes mobile upon hydroxyurea-induced replicative stress (Yu et al, 2003). Notably, their data suggested that such increased RAD51 mobility was unlikely to be a consequence of altered interaction with BRCA2 upon replicative stress. Similarly, Jeyasekharan *et al*. reported increased BRCA2 mobility upon IR-induced DNA damage, which was induced in a manner dependent on the DNA damage signalling kinases ATM and ATR and coincided with active HR events (Jeyasekharan et al, 2010). Given that BRCA2 and PALB2 form a stable complex (B2P2) together with a fraction of RAD51 and that their interaction is important for HR repair, we suggest that the B2P2 complex and associated RAD51 are mobilised together upon genotoxic stress.

The mechanisms that regulate the mobilisation and localisation of HR factors remain incompletely understood but undoubtedly involve PTMs. We propose that variable modes of PTM-mediated chromatin association synergistically govern the recruitment of repair factors to defined regions of chromatin. In undamaged cells, PALB2 is enriched at actively transcribed chromatin through KAT2A/B-mediated ChAM acetylation and H3K36me^3^-mediated MRG15 interaction, while maintaining transient chromatin interaction. This transient mode of association may allow a severance mechanism, in which PALB2 continuously monitors the presence of DNA damage across these regions. Indeed, cells expressing PALB2 7R exhibited an increase in the basal level of γ−H2AX compared to those expressing PALB2 WT, underlining the importance of PALB2 acetylation in unperturbed cells. Upon DNA damage, however, ChAM is deacetylated and, as a consequence, PALB2 affinity for undamaged chromatin is reduced. We suggest that damage-induced ChAM deacetylation allows its recruitment to sites of DNA damage through interaction with BRCA1, fulfilling its function in promoting HR repair. Intriguingly, cells expressing the non-acetylatable PALB2 7R variant exhibited no detectable HR defects, which is in agreement with our finding that DNA damage triggers ChAM deacetylation. These observations highlight the importance of dynamic regulation of PALB2, where both acetylated and non-acetylated forms of PALB2 play critical roles in genome integrity control. Our work also suggests that caution should be exercised in the use of K to Q substitutions for functional studies of lysine acetylation. Robust *in vivo* assessment of acetylation events would ideally involve the development of a chemically modifiable residue that exhibits improved similarity to acetylated lysine in a reversible manner.

More broadly, lysine acetylation is known to control biological processes, such as transcription, based on the metabolic status of the cell (Lee et al, 2014). Therefore, a better appreciation of the role of lysine acetylation in the DDR could expand the scope of studies aiming to understand how reprogrammed metabolism could increase genome instability in cancer cells (Nowell, 1976). Since PTMs are essential for physiological homeostasis and metabolites are necessary cofactors for the deposition of these PTMs, we envision that reprogrammed metabolism in cancer cells could alter the PTM landscape of DDR proteins and hence contribute to genome instability. Considering that glucose concentration in the growth medium can influence the level of protein acetylation in cells (Lee et al, 2014), it will be important to understand whether PALB2 function could be affected by cellular metabolic status. Indeed, our preliminary results demonstrated that extracellular glucose concentration affect PALB2 dynamics, as reducing glucose level in the growth medium increases PALB2 dynamics. It is therefore reasonable to speculate that, in tumour cells with reprogrammed metabolism, protein properties such as chromatin association could be detrimentally altered. Future studies assessing aberrant PALB2 lysine acetylation in cancer tissues and whether KDAC or bromodomain inhibitors, currently in clinical trials for cancer therapy, or alteration of glucose metabolic pathways would modulate PALB2 acetylation may offer therapeutic potentials. Furthermore, deciphering the metabolic regulation of the DDR and DNA repair could highlight additional cellular pathways that might be targeted for cancer therapy, possibly in combination with existing therapies that otherwise may give rise to resistance.

## Author contributions

M.F. and F.E. conceived the study and designed the experimental strategies; M.F., J-Y.B, A.M.C., J.E., and S.H. performed the experiments; J-Y.B, S.S., and L.T. contributed to reagents and analytical tools; M.F. and J.E. analysed data; M.F., J.Y.B, A.M.C., S.S., and F.E. contributed to text and figures; M.F. and F.E. wrote the manuscript.

## Acknowledgements

FE is supported by a Wellcome Trust Senior Research Fellowship in Basic Biomedical Science (101009/Z/13/Z) and is thankful for support from the Edward P Abraham Research Fund. SJS is supported by the Francis Crick Institute which receives its core funding from Cancer Research UK (FC001156), the UK Medical Research Council (FC001156), and the Wellcome Trust (FC001156). LT receives funding from the European Research Council (ERC) Advanced Grant (ERC-2013-340551, Birtoaction). We are also grateful to Nicola O’Reilly (Francis Crick Institute Peptide Chemistry Technology Platform) for synthesis of ChAM peptides, Andrew Jefferson and Carina Mónico (Micron Oxford Advanced Bioimaging) for assistance with microscopy, Christine Ralf (Sir William Dunn School) for technical support in generating constructs, and Chris Norbury (Sir William Dunn School) for critical reading of the manuscript.

## Competing interests

The authors declare no financial and non-financial competing interests on this study.

## Data availability

The mass spectrometry proteomics data have been deposited to the ProteomeXchange Consortium via the PRIDE (Perez-Riverol et al, 2019) partner repository with the dataset identifier PXD014678 and PXD014681.

## Supplemental Material and Methods

### Plasmids

For bacterial expression, full-length PALB2 and fragments 1 to 4 were PCR amplified using primer pairs, numbered 1 and 2 (Fr. 1), 3 and 4 (Fr. 2), 5 and 6 (Fr. 3), or 7 and 8 (Fr. 4) listed in Table 1, from pCMV-SPORT6-PALB2 (IMAGE clone 6045564, Source BioSciences) and cloned into the BamHI/NotI sites of the pGEX-6P-1 vector (GE Healthcare). For mammalian expression, ChAM fragments of varied length were first PCR amplified using primer pairs, numbered 9 and 11 (#1), 9 and 12 (#2), 10 and 12 (#3), 10 and 11 (#4), or 10 and 13 (#5) listed in Table 1, and cloned into the BamHI/XhoI sites of the pENTR3C Gateway entry vector (Thermo Fisher Scientific). To generate GFP-fusion or GST-fusion expression vectors, ChAM fragments were then transferred from pENTR3C to pcDNA-DEST53 (GFP-fusions) or pDEST15 (GST-fusions) (Invitrogen) using Gateway cloning. PALB2 Q and R mutations were introduced in the pENTR3C-ChAM#1 or pENTR3C-PALB2 vectors by inverse PCR (5’-phosphorylated oligonucleotides containing the desired mutations were used to create blunt-ended products, which were then recircularised by intramolecular ligation). GST-ChAM missense variants 7Q, 3Q4K and 3K4Q were respectively generated using primer pairs, numbered 18 and 20 (7Q), 18 and 15 (3Q4K), or 14 and 20 (3K4Q) listed in Table 1. PALB2 7Q and 7R full length missense variants were respectively generated using primer pairs, numbered 18 and 19 (7Q), or 16 and 17 (7R) listed in Table 1. To generate N3xFLAG-fusion expression vectors, PALB2 variants were then transferred from pENTR3C to pcDNA5/FRT-GW/N3×FLAG (Bleuyard et al, 2017) using Gateway cloning.

**Table 1.**
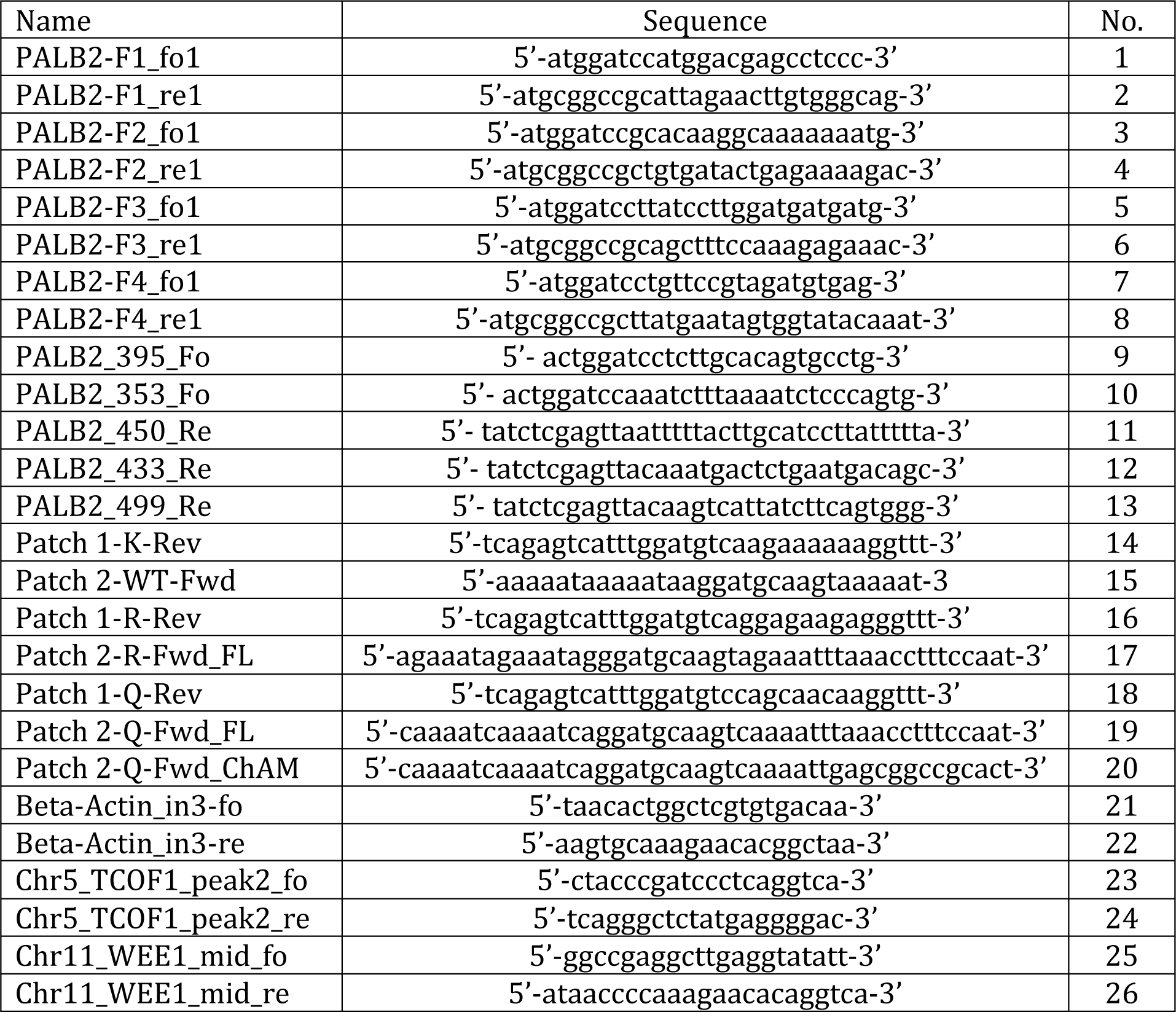
List of oligonucleotides used in this study

### Chemical cell fractionation and whole-cell extract

HEK293T cells transiently expressing GFP-ChAM variants were collected using TrypLE Express reagent (Gibco), washed twice with ice-cold PBS and resuspended in Sucrose buffer (10 mM Tris-HCl pH 7.5, 20 mM KCl, 250 mM Sucrose, 2.5 mM MgCl_2_, 10 mM benzamidine hydrochloride and P2714 protease inhibitor cocktail from Sigma-Aldrich), using 1 mL buffer per 100 mg of weighed cell pellet. After addition of Triton X-100 (Sigma) to a final concentration of 0.3% w/v, the cell suspensions were vortexed three times for 10 s with 1 min intervals. The intact nuclei were collected by centrifugation for 5 min at 500 x g, 4°C, and the supernatant collected as cytoplasmic fraction. The nuclei pellet was washed once with ice-cold sucrose buffer and resuspended in ice-cold NETN150 buffer (50 mM Tris-HCl pH 8.0, 150 mM NaCl, 2 mM EDTA, 0.5% NP-40, 10 mM benzamidine hydrochloride and P2714 protease inhibitor cocktail from Sigma), using 400 µL buffer per 100 mg of initial weighed cell pellet. After 30 min incubation on ice, the chromatin fraction was collected by centrifugation for 5 min at 500 x g, 4°C and the supernatant collected as nuclear soluble fraction. The chromatin pellet was washed once with ice-cold NETN150 buffer and finally solubilised for 1 h on ice in NETN150 buffer containing 2 mM MgCl_2_ and 125 U/mL Benzonase nuclease (Merck Millipore), using 250 µL buffer per 100 mg of initial weighed cell pellet. Cytoplasmic, nuclear soluble and chromatin-enriched fractions were centrifuged for 30 min at 16,100 x g, 4°C to remove cell debris and insoluble material. For whole-cell extract, cells were directly lysed in NETN150 buffer containing 2 mM MgCl_2_ and 125 U/mL Benzonase for 1 h on ice and centrifuged for 30 min at 16,100 x g, 4°C to remove cell debris and insoluble material.

### Cell survival

In 96-wells plates, EUFA1341 cells complemented with FLAG-PALB2 WT and variants were exposed to increasing concentration of Olaparib (0-2 μM). After 4 days of incubation at 37°C, cell proliferation was measured using WST-1 reagent (Roche Applied Science) following manufacturer protocol.

### Laser damage

To perform laser damage experiments, cells were plated onto cell view cell culture dishes (3.5mm) and analysed in phenol red-free medium (Leibovitz L15-medium, Gibco life technologies). Images analysis were performed on a spinning-disk confocal microscope (Ultra-View Vox, Perkin Elmer) mounted on an IX81 Olympus microscope with an Olympus 60x 1.4 oil PlanApo objective, in a controlled chamber at 37°C and 5% CO2 (TOKAI HIT stage top incubator). The fluorescence signal was detected using an EMCCD camera (ImagEM, Hamamatsu C9100-13). To perform laser damage experiments, cells were irradiated in the DAPI channel at maximum laser power with a single pulse for 90 ms, using a straight line crossing the whole cellular surface. After bleaching, GFP fluorescence recovery was monitored within the bleached area every 30 s for 5 min. Laser damage parameters were controlled using the Volocity software 6.0 (QuorumTechnologies). PALB2 recruitment at sites of laser damage was quantified in Fiji/Image G (Image / Stacks/ Plot Z-axis Profile) by measuring the EGFP fluorescence intensity within a region of interest at site of damage over fifteen minutes after laser-induced damage. Images were recorded every 30 s during this period. Fluorescence intensity at the damaged site over the time course experiment was corrected for background and overall loss of fluorescence intensity in the nucleus (non-damaged region of interest) and normalised to the pre-damaged value.

### ChIP-qPCR

ChIP experiments were performed from 20×10^6^ cells as described previously (Bleuyard et al, 2017). FLAG antibody used to perform ChIP experiments is described in the antibodies section of Material and Methods. Quantitative real-time PCR (qPCR) experiments have been performed on a Rotorgene Q Real-Time PCR system (Qiagen) using the SensiFAST SYBR No-Rox kit (Bioline). Oligonucleotides used to perform qPCR against coding regions of *ACTB*, *TCOF1* and *WEE1* genes are listed in Table 1 (numbered 21 to 26).

### Cell synchronisation

Cell synchronisation experiments were performed by double thymidine block as previously described (Esashi et al, 2005). Briefly, cells were first blocked in G1/S phase with 2 mM thymidine treatment for 18 hr, release in fresh media for 9 h and blocked again at G1/S phase with 2 mM thymidine treatment for 16 h. Subsequently, cells were release in fresh media and collected at 0 and 3 h after release for DNA content analysis by flow cytometry (FACS).

### Cell cycle analysis by flow cytometry

Cell cycle analysis was performed by DNA content measurement by propidium iodide (PI) staining followed by FACS analysis. After cell collection by centrifugation at 500 x g at 4°C, cells were fixed in 70% ethanol. After cell permeabilisation for 5 min on ice with 0.1% Triton contained in 0.1% BSA resuspended in PBS, DNA was stained for 3 h in the dark in a PI solution containing 0.1% BSA, 0.1 mg/ml of RNase and 2 ug/ml of PI. DNA content was analysed from 10000 cells on a FACS calibur machine (BD Calibur) and FACS data was analysed using FlowJo software V10.

### Mass spectrometry

*Protein sample digestion*: proteins contained in solution were urea-denaturated in 4 M urea dissolved into 50 mM triethylammonium bicarbonate (TEAB), reduced with 10 mM tris-(2-carboxyethyl)phosphine (TCEP) for 30 min at room temperature, alkylated with 50 mM chloroacetamide for 30 min at room temperature in the dark, digested with endoproteinase Lys-C (Roche) for 2 h at 37°C, followed by trypsin digestion for 16 h at 37°C. Before trypsin digestion, the urea concentration was diluted down to 1 M into 50 mM TEAB and calcium chloride was added at 1 mM final. The digestion steps were performed in a Thermomixer compact (Eppendorf) shaking at 650 rpm. Trypsin digestion was stopped by adding 1% Trifluoroacetic acid (TFA) final. Digest samples were centrifuged for 30 min at 4°C to remove aggregates. Peptides mixtures were further desalted onto hand-made C18 tips, as following: the C18 tip was first washed twice with 100% acetonitrile after 5 min centrifugation at 2000rpms at room temperature. Peptides were further loaded onto the C18 tip by centrifugation at 4,000 rpm at room temperature. After two washing steps into 0.1% TFA solution, desalted peptides were eluted into 50% acetonitrile/0.1% TFA solution and dried using a SpeedVac before LC-MS/MS analysis. All MS/MS spectra for acetylated PALB2 peptides are provided in the supplementary material.

Acetylated lysine abundance values were calculated as following: PALB2 acetylated peptides specifically identified in KAT2A, KAT2B and KAT2A mutant conditions were extracted and compared. Spectral counts for similar acetylated peptides identified across these three conditions were divided by the total PALB2 spectral counts identified in each condition, to normalise for experimental variation encountered between independent runs. This calculation was performed for each acetylated peptide identified, and normalised values were then summed up to define the acetylation abundance factor of PALB2 in each condition tested.

**Figure S1.**
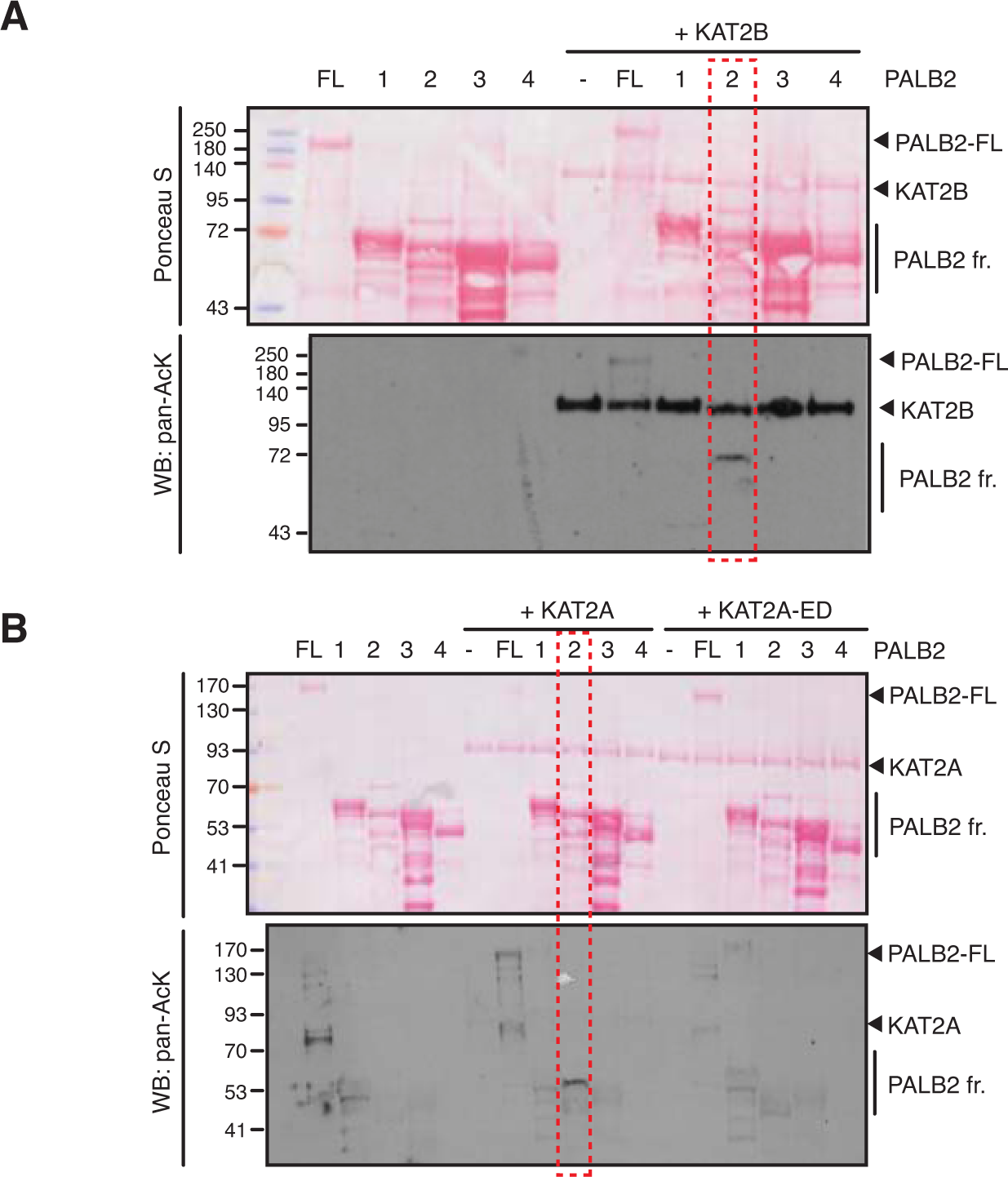
PALB2 is directly acetylated by KAT2A and 2B. ***A-B.*** *In vitro* acetylation of PALB2 by KAT2B (*A*) or KAT2A and a catalytically inactive KAT2A-ED (*B*). Purified PALB2 full-length and fragments 1 to 4 (Fig. 1A) were incubated with either purified KAT2B, KAT2A or KAT2A-ED in the presence of acetyl-CoA, followed by SDS-PAGE. Total and acetylated proteins were visualized by Ponceau S staining and anti-acetyl lysine (pan-AcK) western blot, respectively. Acetylated PALB2 fragment 2 by KAT2B or KAT2A is highlighted with red dashed boxes.

**Figure S2.**
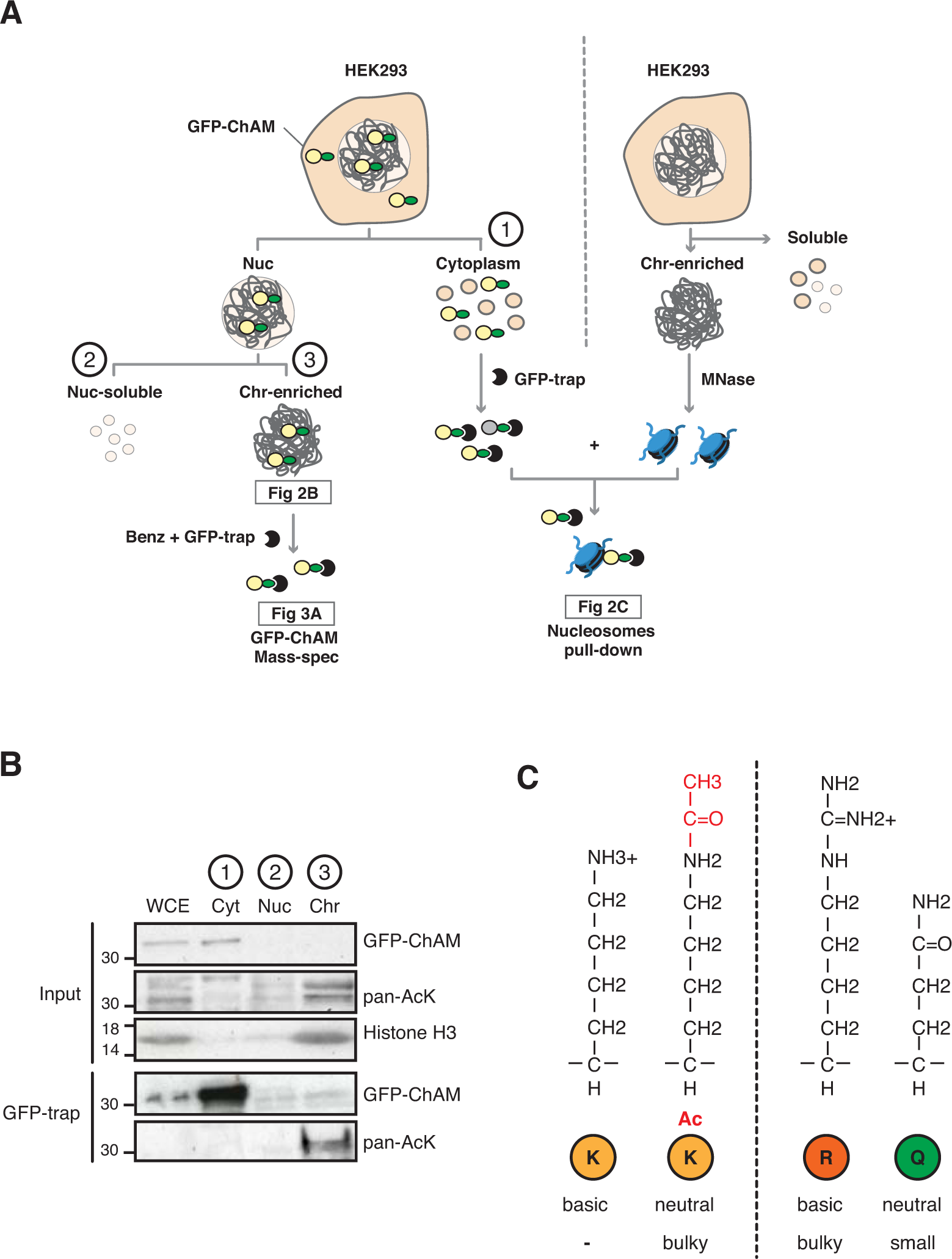
Chromatin-associated ChAM is highly acetylated *in vivo*. ***A.*** Depiction of experimental procedures for subcellular fractionation of HEK293 cells transiently expressing GFP-ChAM variants. Chromatin enriched GFP-ChAM was detected by WB (Fig. 2B), or following benzonase treatment, affinity purified using GFP-trap and was analysed for acetylation by MS (Fig. 3A). For nucleosome binding assay, shown in Fig 2C, nucleosomes were partially purified from separate, non-transfected HEK293, and incubated with GFP-ChAM purified from cytoplasmic fraction of transfected HEK293. ***B***. Subcellular distribution of GFP-ChAM overexpressed in HEK293 cells, in cytoplasmic (Cyt), nuclear soluble (Nuc) and chromatin-enriched (Chr) fractions. Histone H3 was used as a control for cellular fractionation. GFP-ChAM and acetylated GFP-ChAM were respectively detected by anti-GFP and anti-acetyl-lysine (pan-AcK) antibodies. ***C.*** Depiction of lysine (K), acetyl-lysine (K-ac), arginine (R) and glutamine (Q) structures, showing differences in size and charge for each residue.

**Figure S3.**
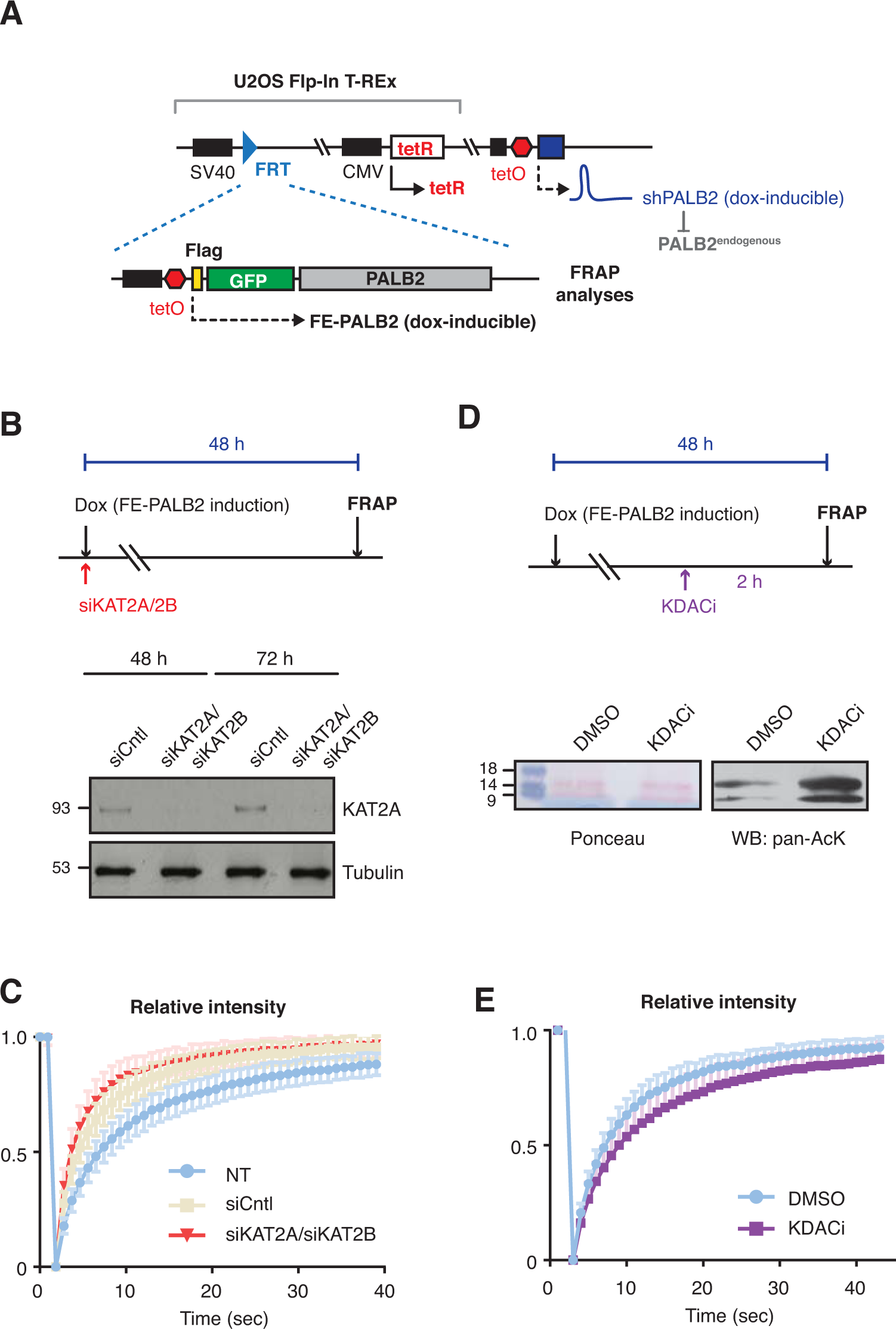
PALB2 mobility increases upon deacetylation. ***A.*** Depiction of cell lines used for FRAP assays of full-length PALB2. U2OS Flp-In T-REx, harboring inducible shRNA targeting 3’UTR of endogenous PALB2 (U2OS-shPALB2) was used as parental cells. FE-PALB2 under tetO promoter, integrated at a defined FRT site, was induced by doxycycline exposure. ***B.*** Diagram of experimental procedure, assessing the impact of KAT2A/B using FRAP against FE-PALB2. KAT2A protein levels were measured by western blot analysis of U2OS-shPALB2 FE-PALB2 cells treated with siRNA targeting KAT2A and KAT2B (siKAT2A/siKAT2B) or scramble small interfering RNA (siCntl) (negative control) for 48 or 72 h. Tubulin was used as a loading control. ***C***. Relative fluorescence intensity of FE-PALB2 in WT, siCntl, and siKAT2A/B treated cells, related to FRAP analysis shown in Fig. 3E and F. ***D***. Diagram of experimental procedure, assessing the impact of KDAC inhibition using FRAP against FE-PALB2. Level of pan-acetylated histones in U2OS-shPALB2 FE-PALB2 cells treated with DMSO or deacetylases inhibitors (KDACi) is shown. Ponceau S staining of proteins after transfer onto nitrocellulose membrane was used as a loading control. ***E***. Relative fluorescence intensity of FE-PALB2 in mock and KDACi treated cells, related to FRAP analysis shown in Fig. 3G and *H*.

**Figure S4.**
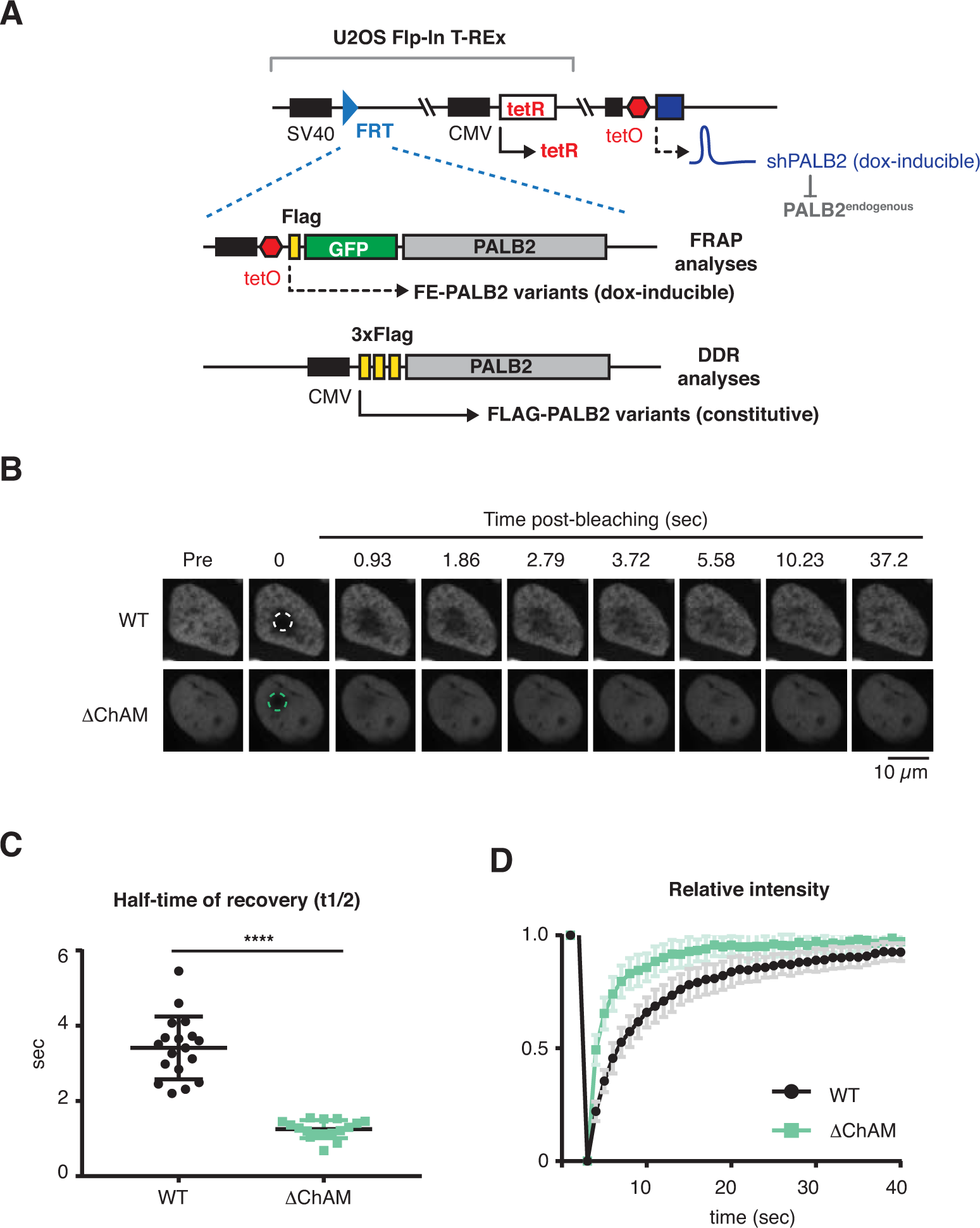
PALB2 ChAM 7K-patch is required to maintain genome stability. ***A.*** Depiction of cell lines used in this study. U2OS-shPALB2 was used as parental cells. FE-PALB2 variants under tetO promoter or FLAG-PALB2 variants under CMV promoter were integrated at a defined FRT site. ***B-D***. FRAP analysis of FE-PALB2 wild-type (WT) or harboring an internal deletion of the ChAM (ΔChAM) conditionally induced in U2OS-shPALB2. Images of FE-PALB2 in live cells before bleaching (pre-bleaching) and during the recovery time after bleaching, plots of half-time recovery times (s) after photobleaching and relative fluorescence intensity are shown in *C*, *D* and *E*, respectively.

**Figure S5.**
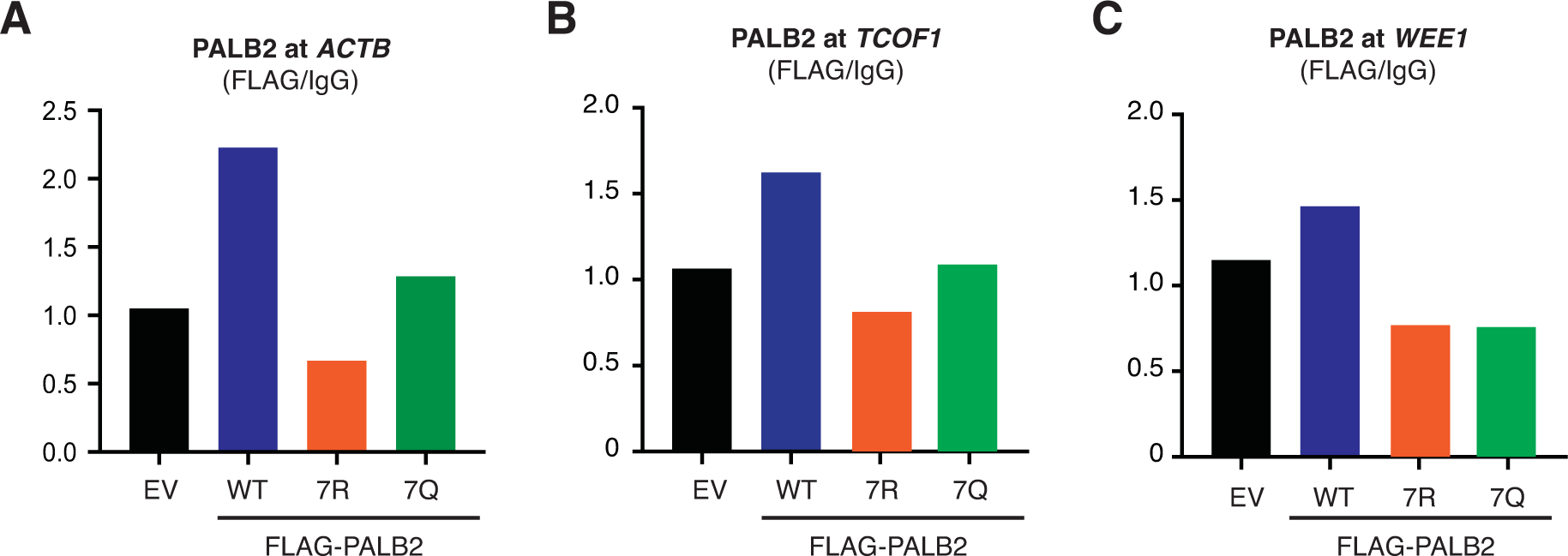
PALB2 ChAM 7K-patch is required to maintain genome stability. ChIP-qPCR analysis defecting FLAG-PALB2 WT, 7Q and 7R enrichment at known PALB2-bound loci (Bleuyard et al, 2017a), namely the coding regions of *ACTB (**A**)*, *TCOF1 (**B**)* and *WEE1* (***C***) genes.

**Figure S6.**
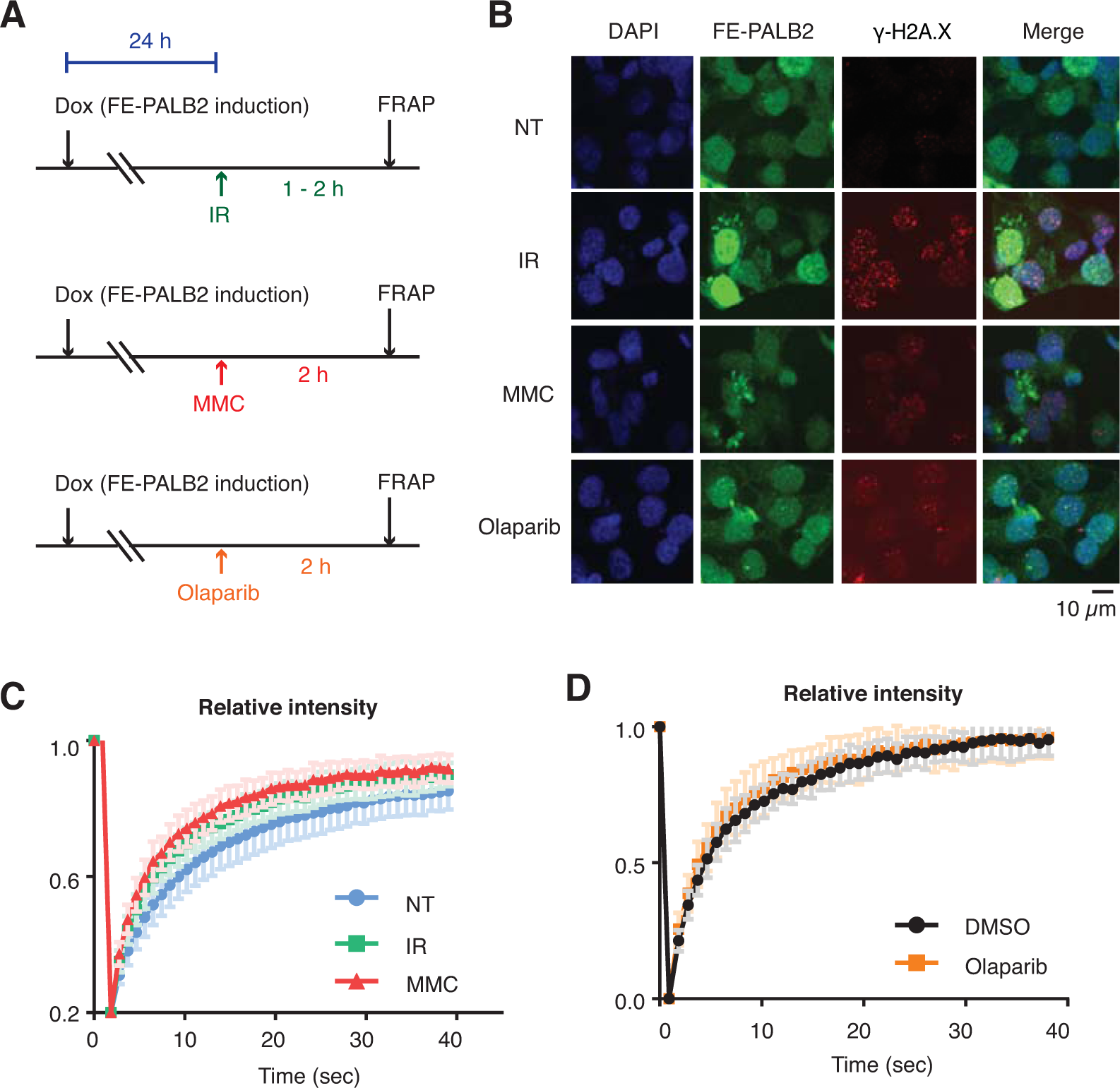
PALB2 mobility increases upon genotoxic stress. ***A***. Diagrams of experimental procedures, assessing the impact of IR, MMC and olaparib using FRAP against FE-PALB2. ***B***. γ-H2AX levels in untreated or damaged cells (4 Gy IR, 1 mM MMC, or 2.5 μM olaparib) as measured by anti-γ-H2AX immunofluorescence staining. ***C***. Relative fluorescence intensity of FE-PALB2 in either undamaged or DNA damaging conditions (4 Gy IR or 1 mM MMC) measured after photobleaching of cells shown in Fig. 6D and E. ***D***. As in C, except cells were treated with DMSO vehicle or 2.5 μM olaparib, related to Fig. 6F and G.

**Figure S7.**
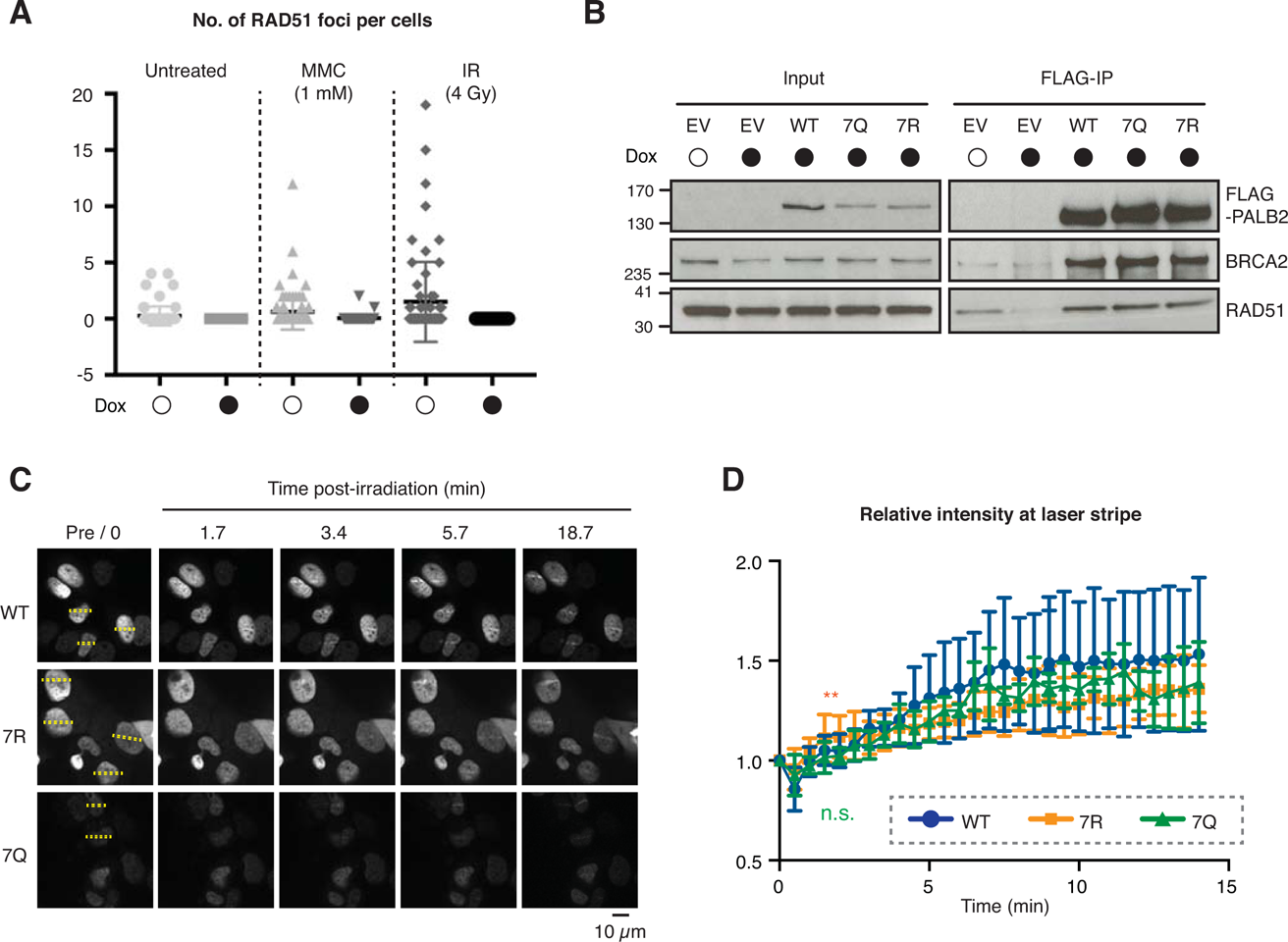
PALB2 ChAM 7K-patch promotes HR upon DNA damage. ***A*.** Number of RAD51 foci per cell in U2OS-shPALB2 without depletion of endogenous PALB2 (white circles) or upon depletion (black circles), in either untreated or DNA damaging conditions (1 mM MMC or 4 Gy IR). ***B.*** FLAG-PALB2 was pulled-down from U2OS-shPALB2 cells stably expressing either FLAG-tagged PALB2 variants, upon doxycycline-induced endogenous PALB2 depletion. Cells expressing only the FLAG-tag (EV) were used a negative control. BRCA2 and RAD51 interaction with PALB2 was assessed by western blot. ***C.*** Analysis of FE-PALB2 variants recruitment to DNA damage sites induced by laser micro-irradiation. Dashed boxes indicate irradiated areas. ***D.*** Quantitation of the amount of FE-PALB2 variants recruited to laser damage sites. The relative fluorescence intensity at laser damage sites was calculated by dividing the fluorescence intensity of the damaged area over the time course experiment by the fluorescence intensity of the non-damaged region of the nucleus and normalisation using the pre-damaged fluorescence value as a reference.

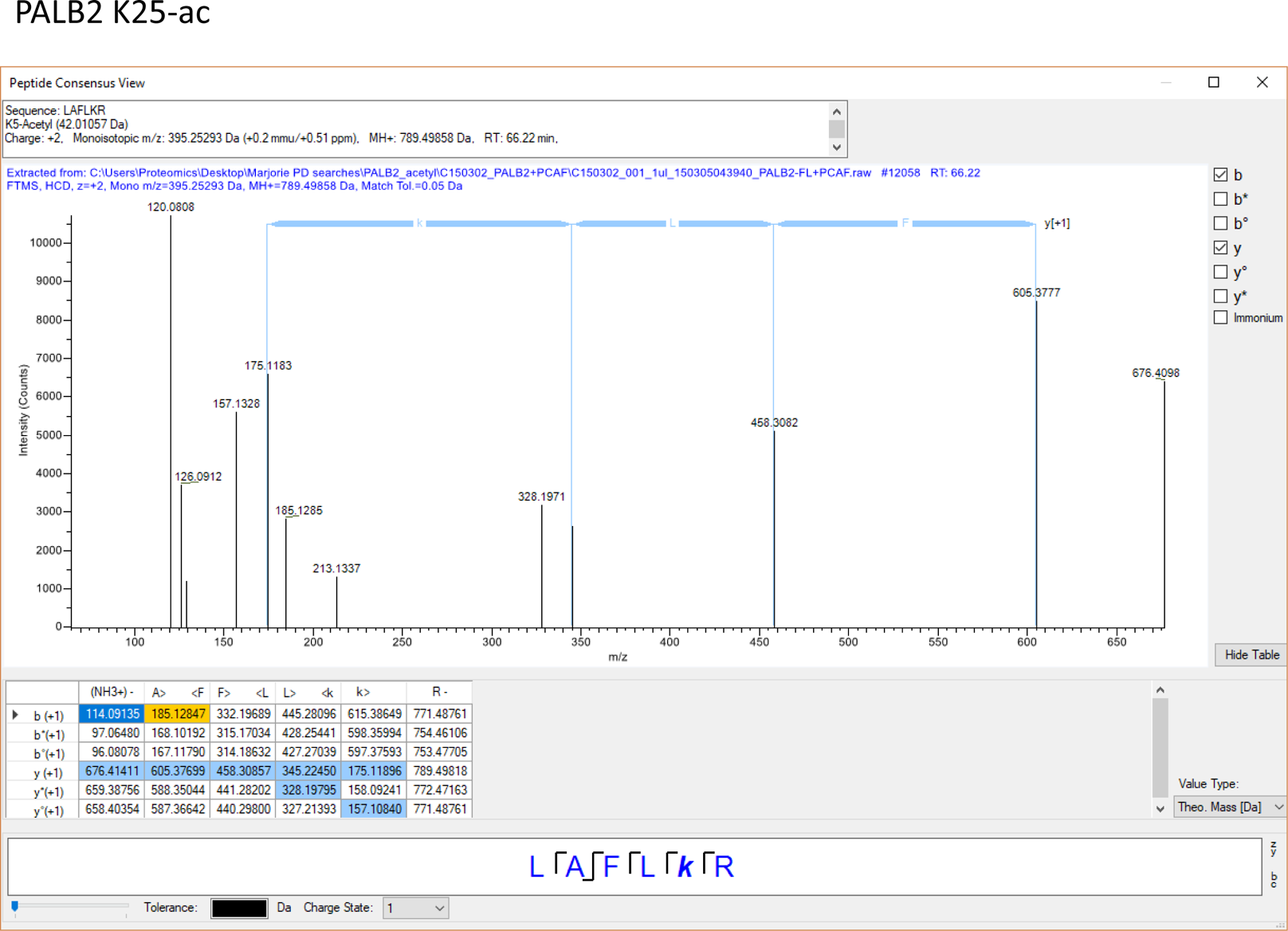

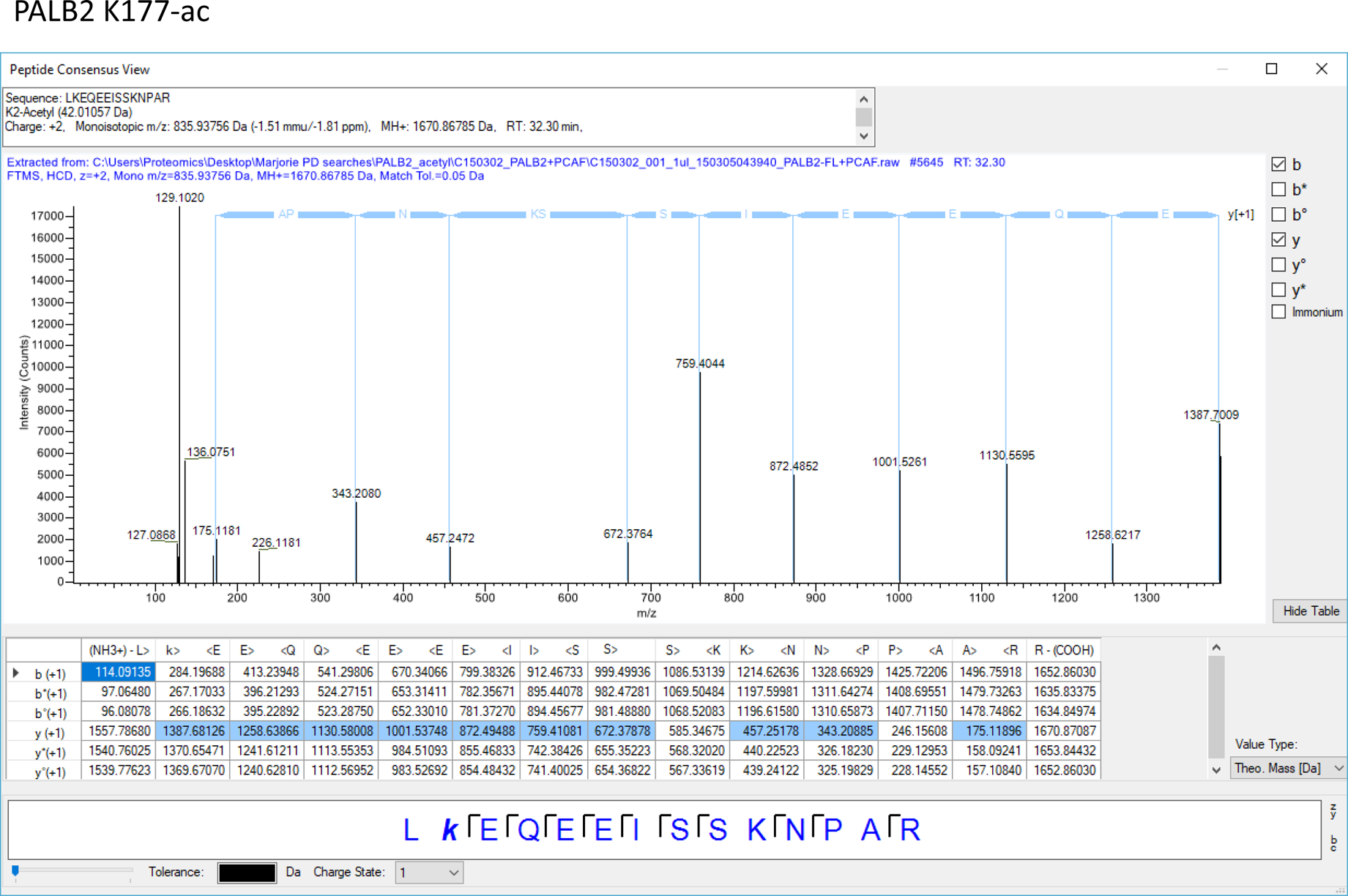

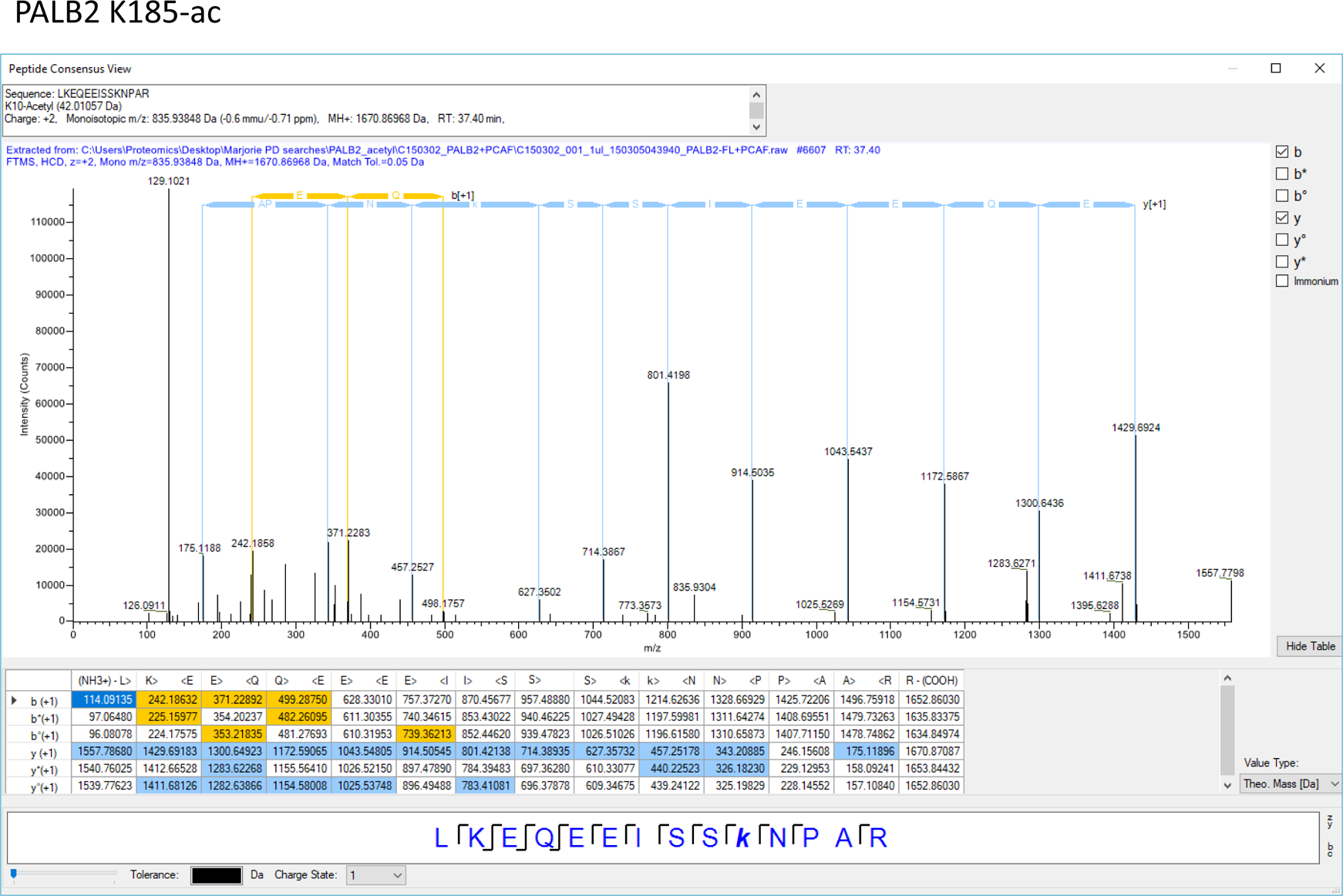

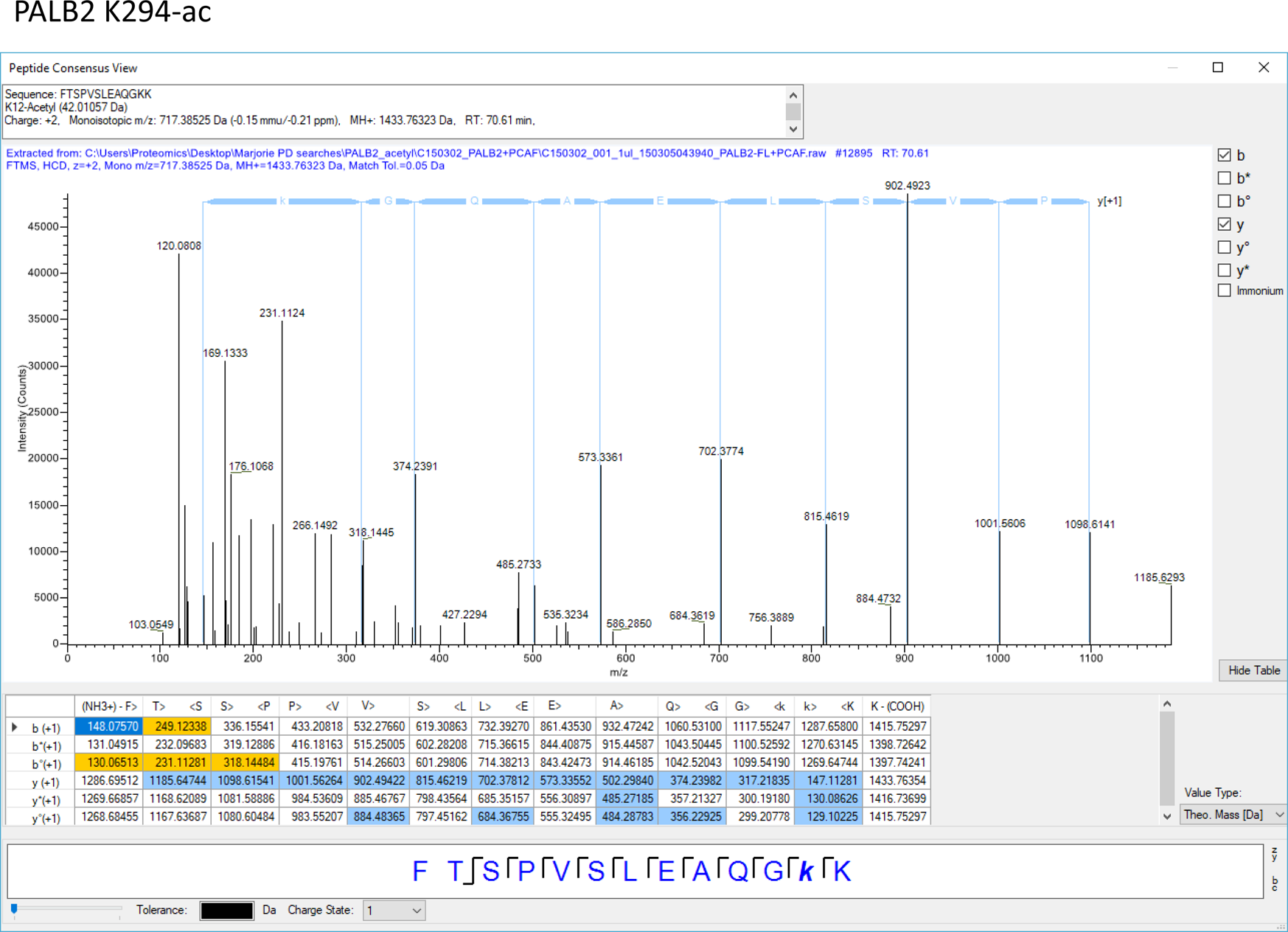

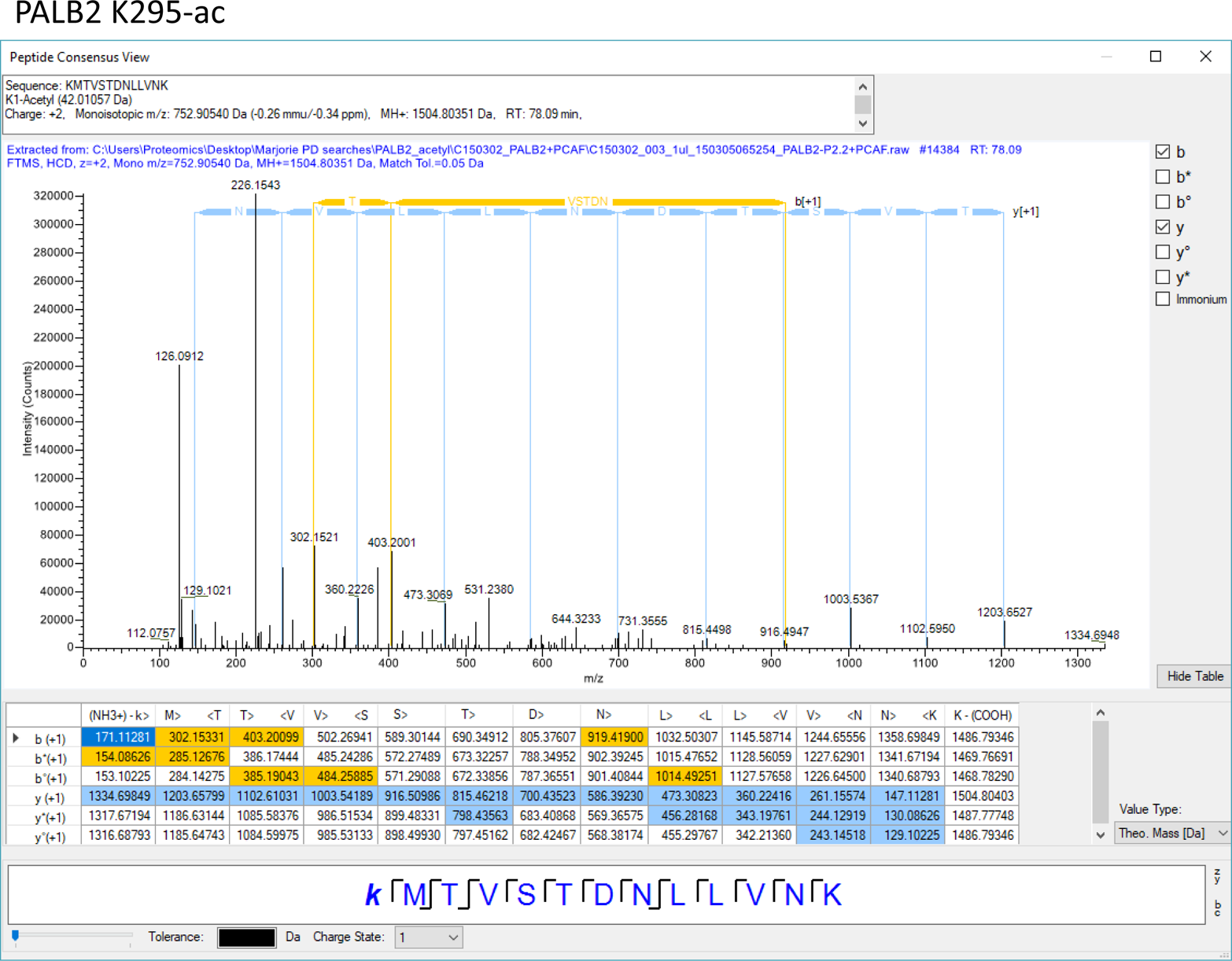

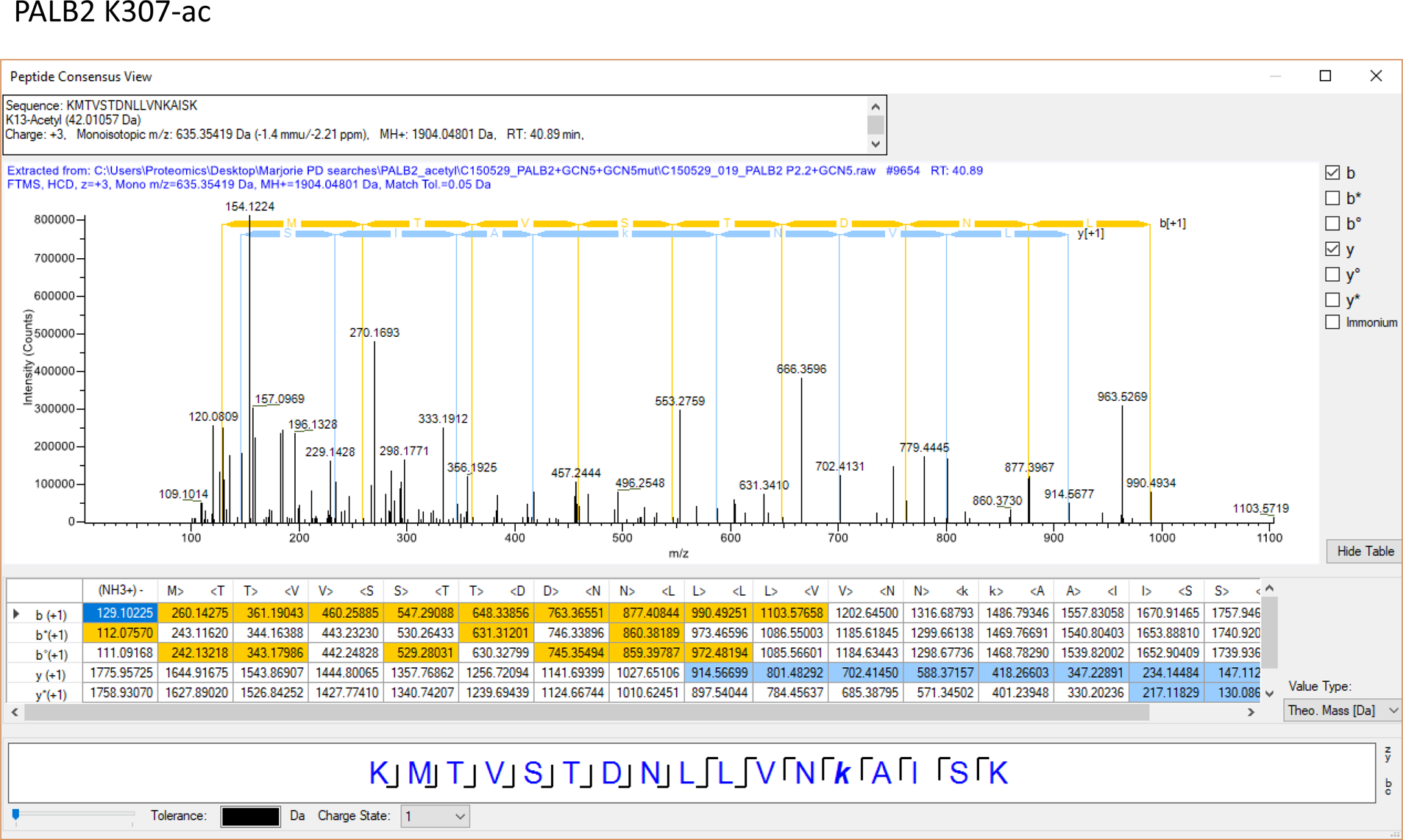

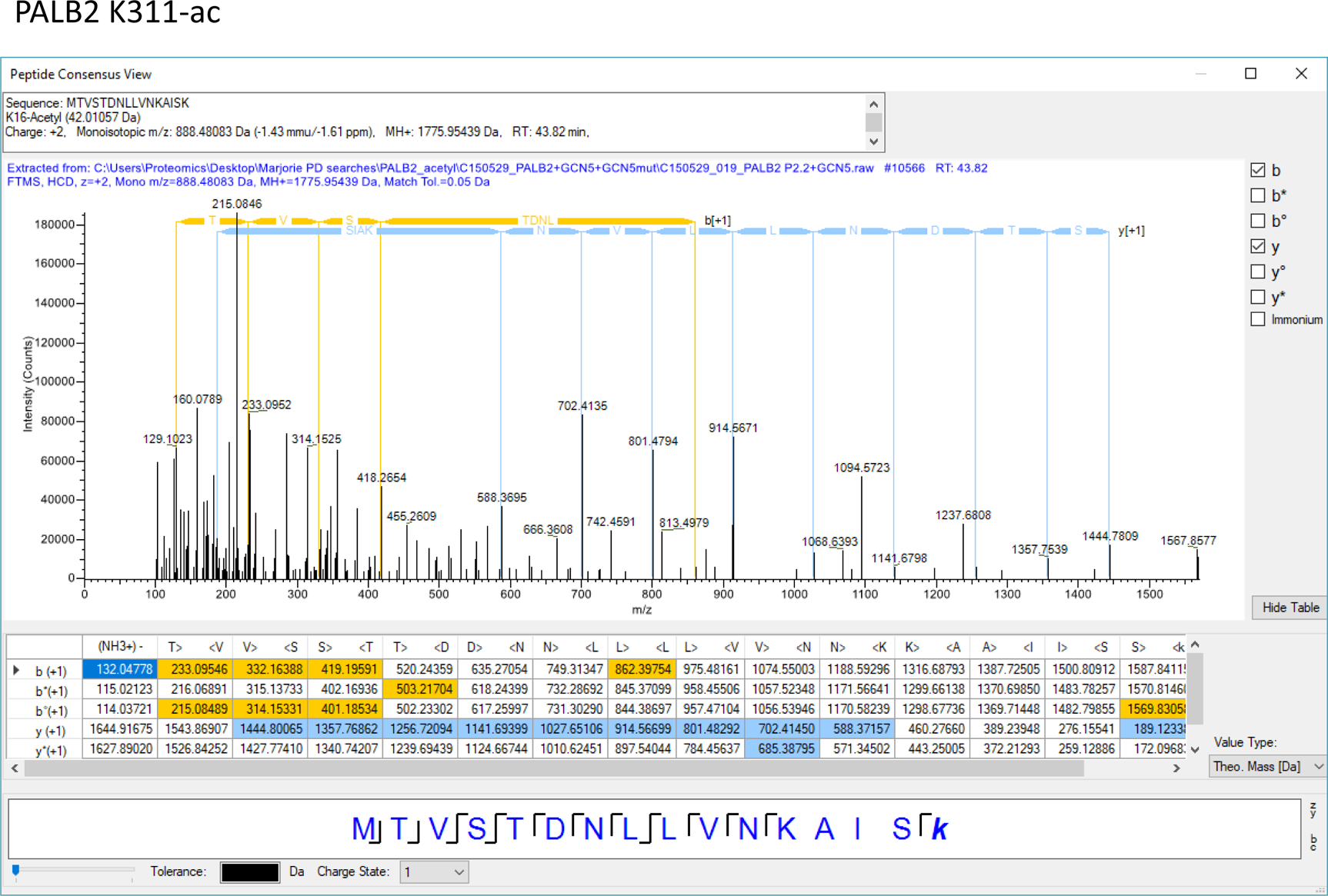

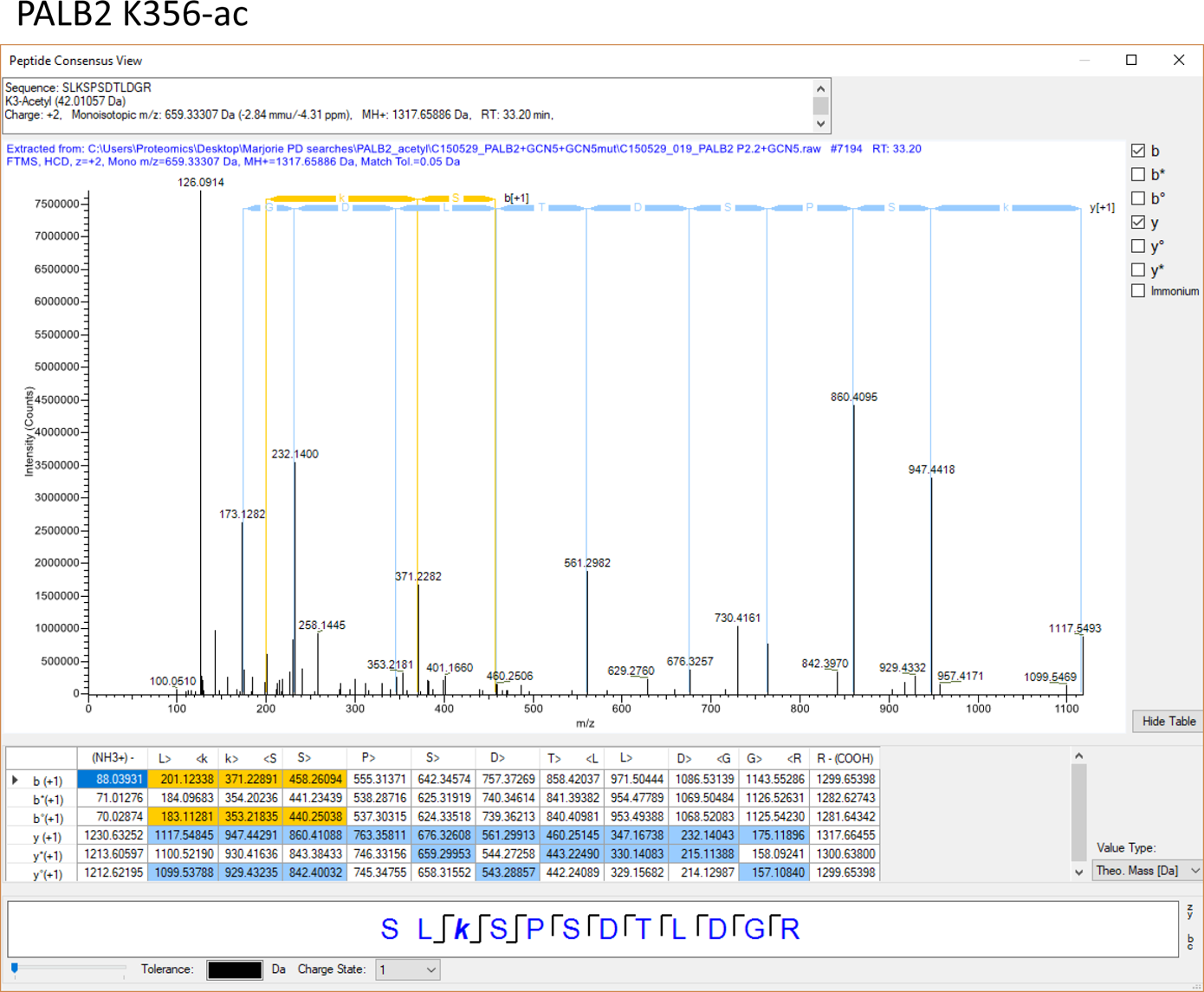

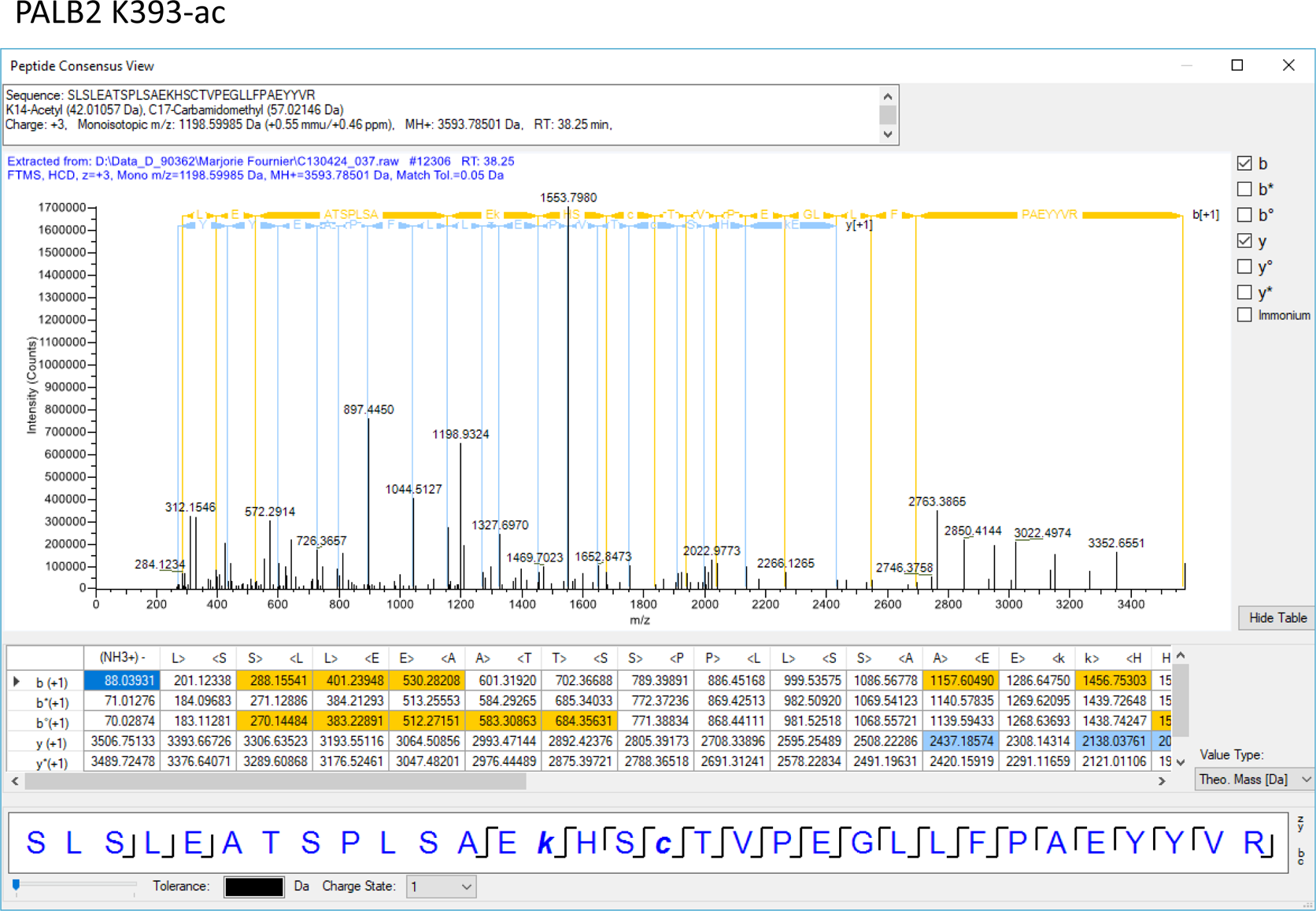

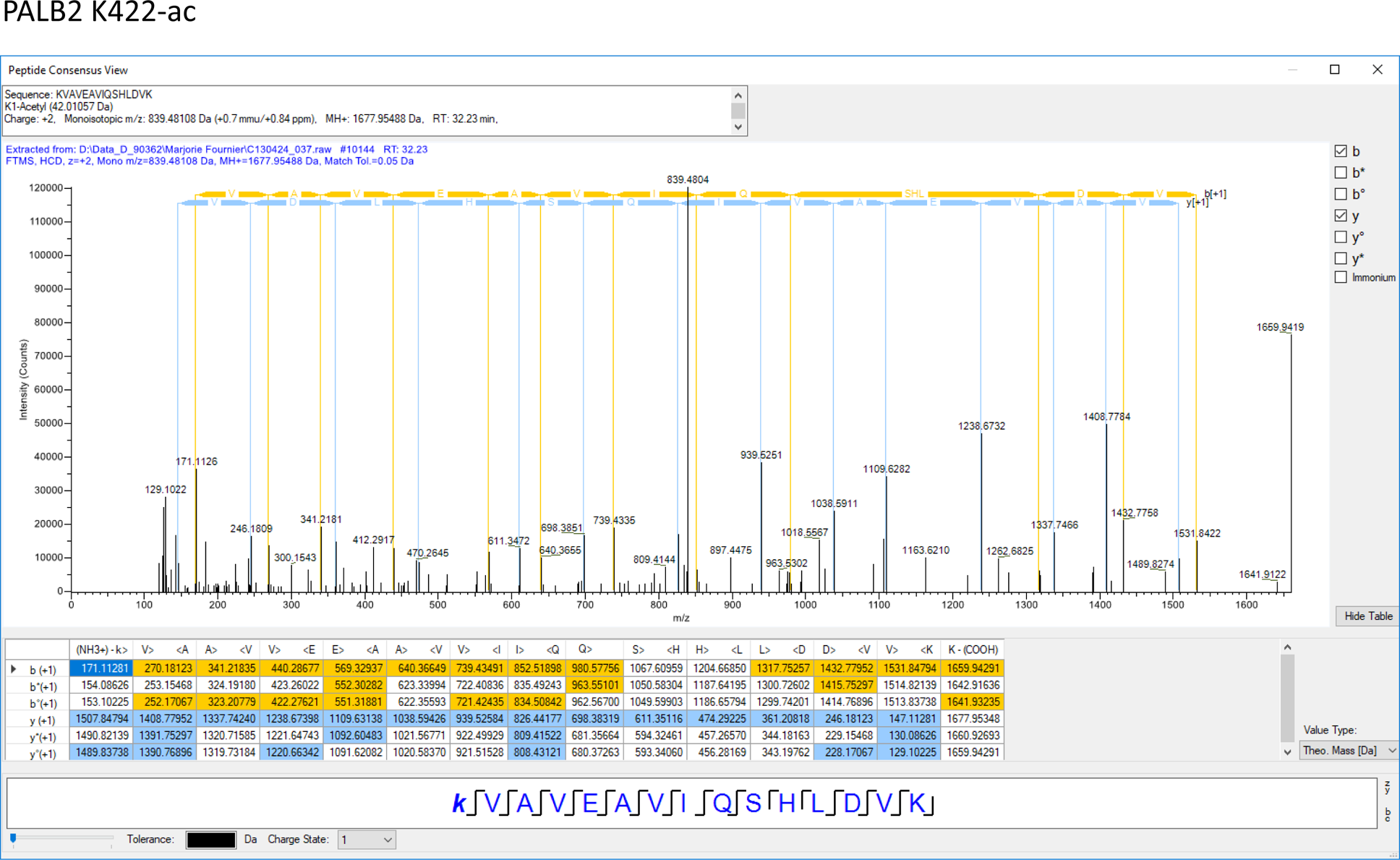

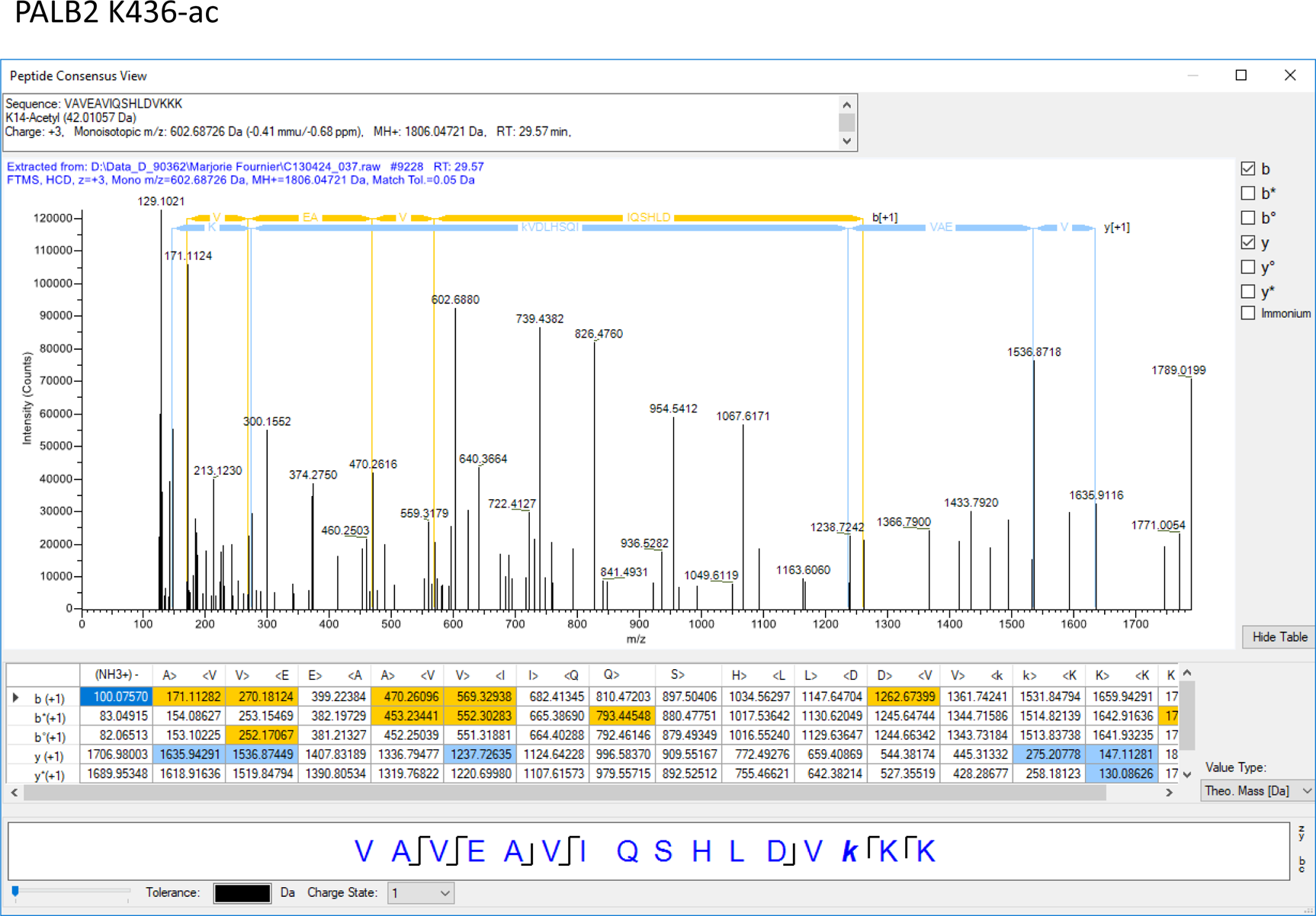

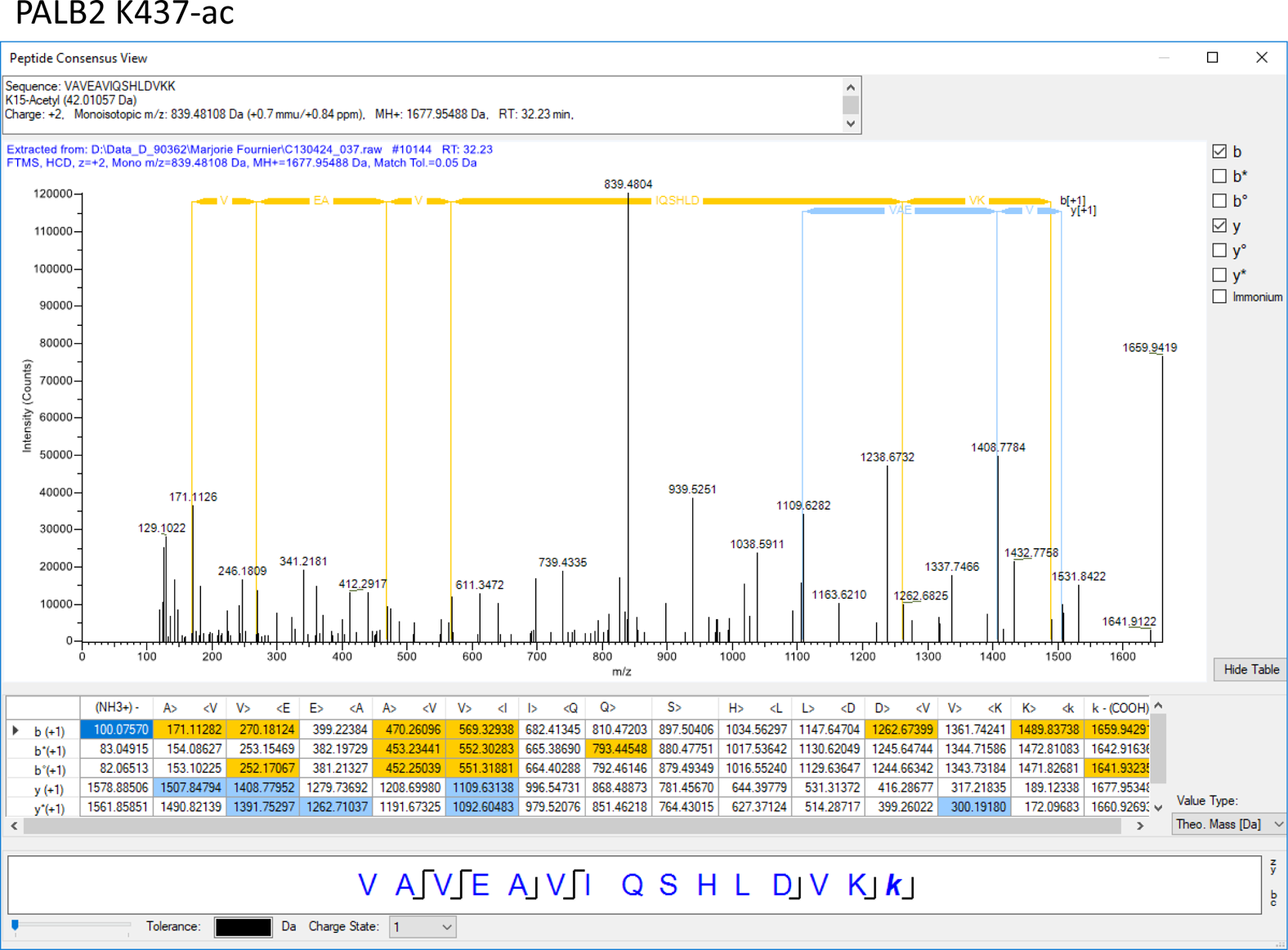

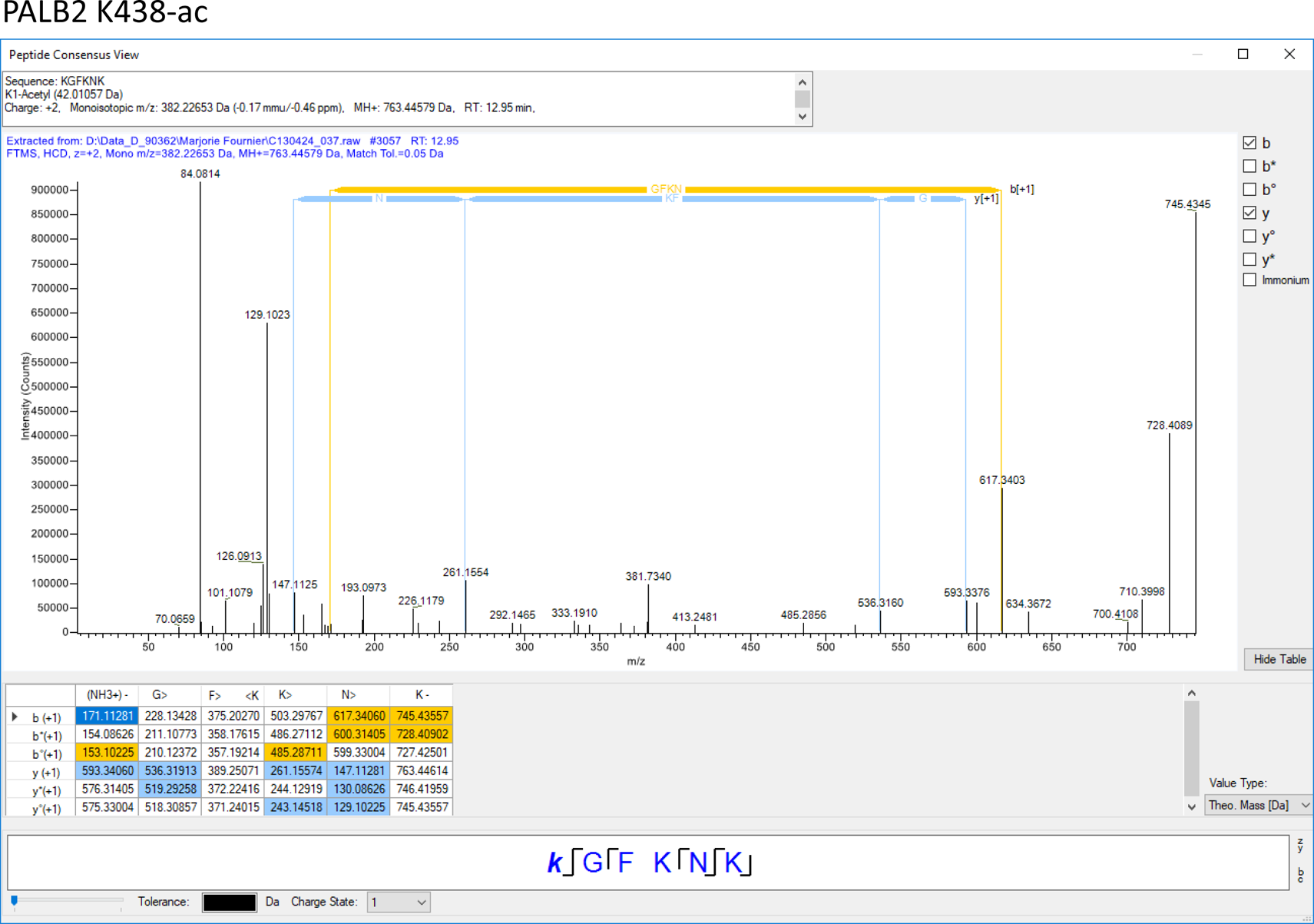

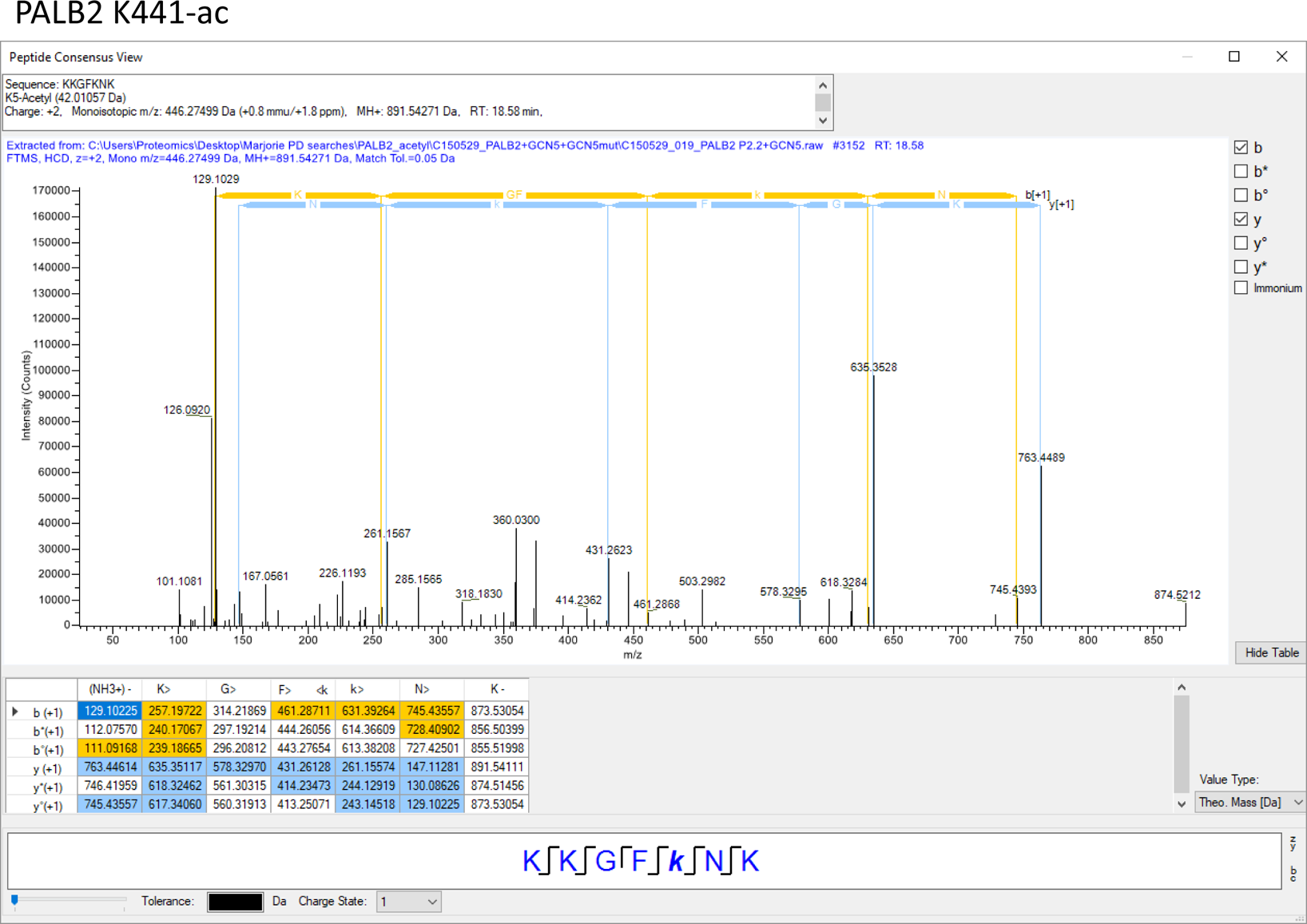

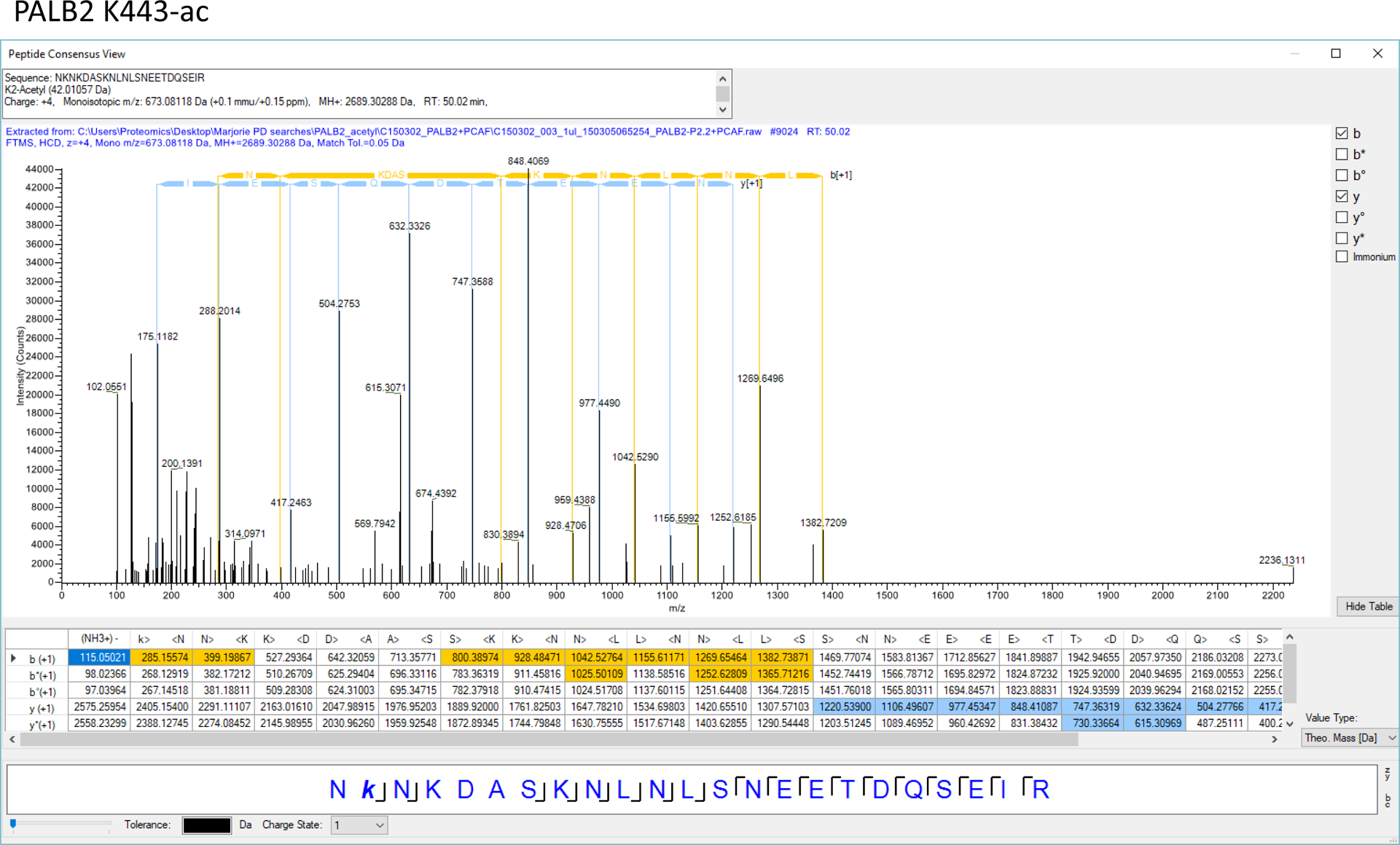

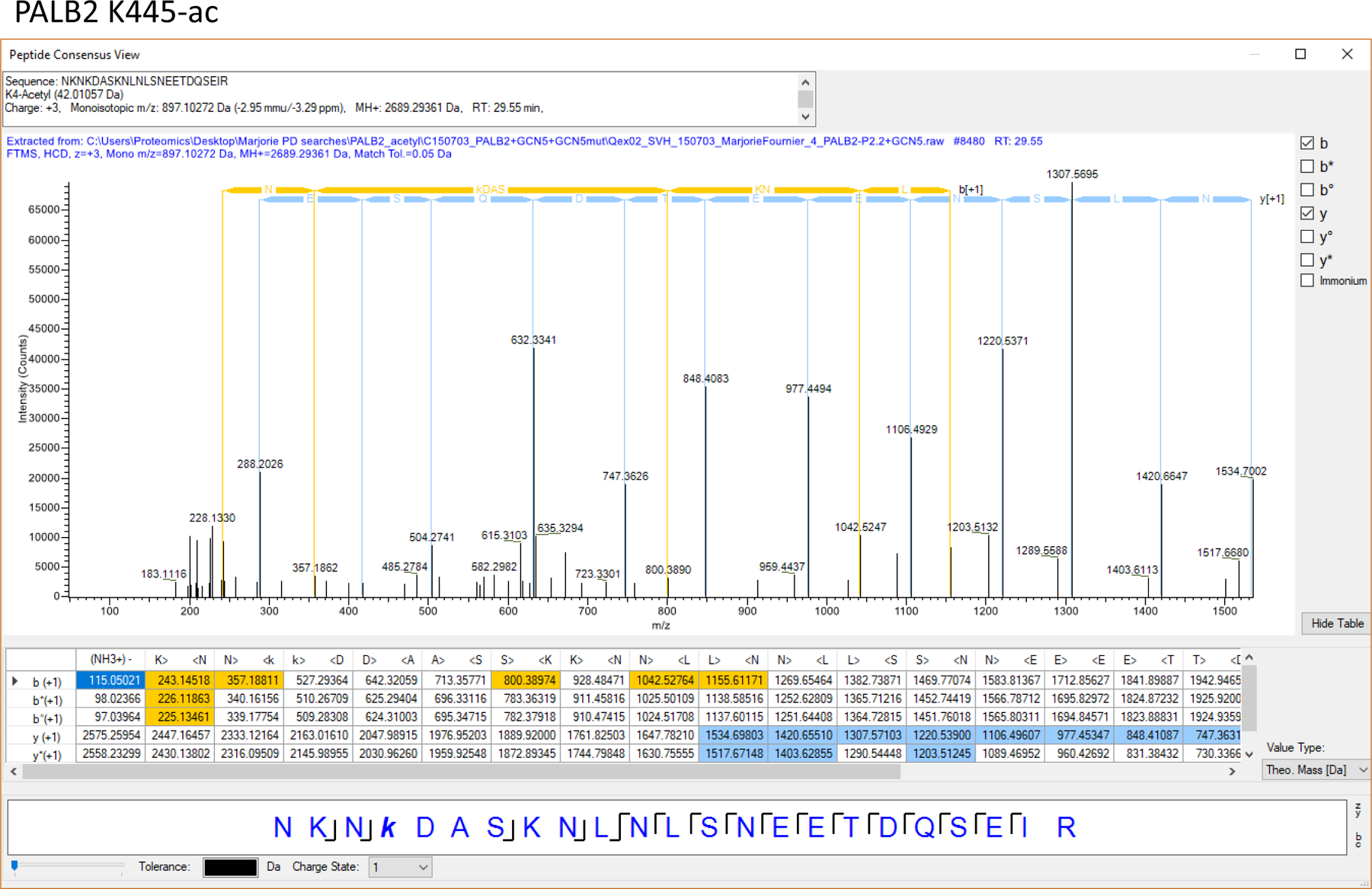

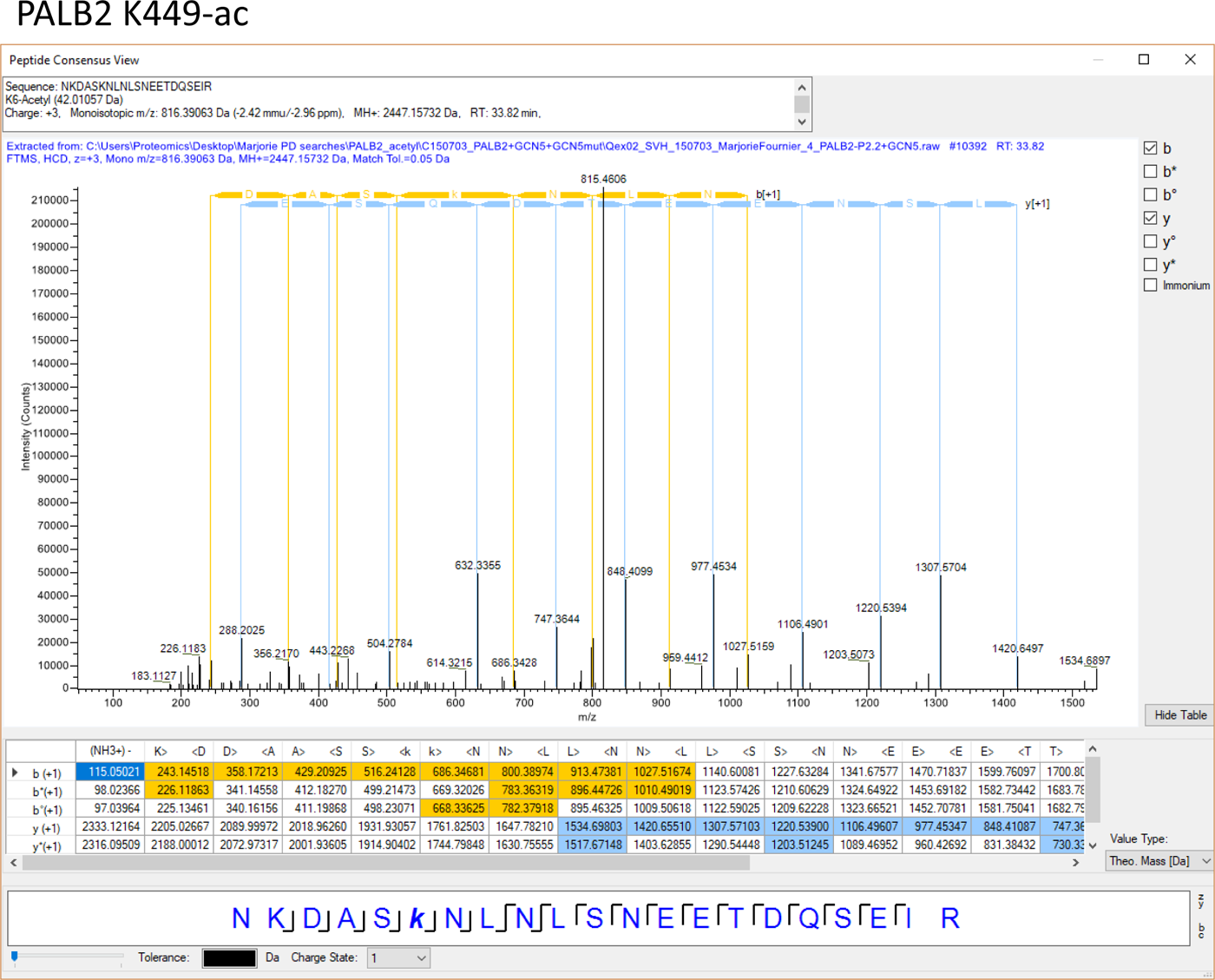

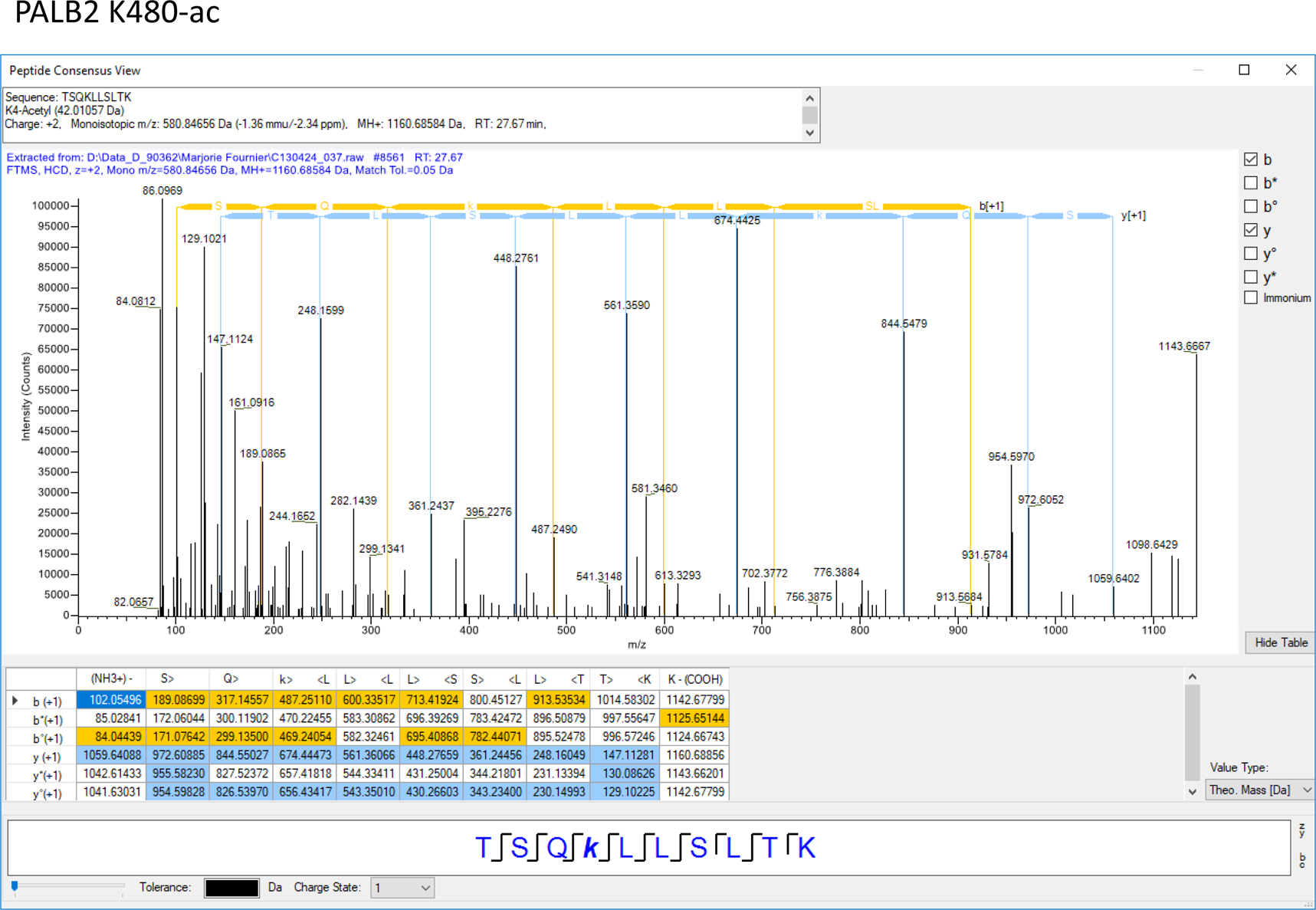

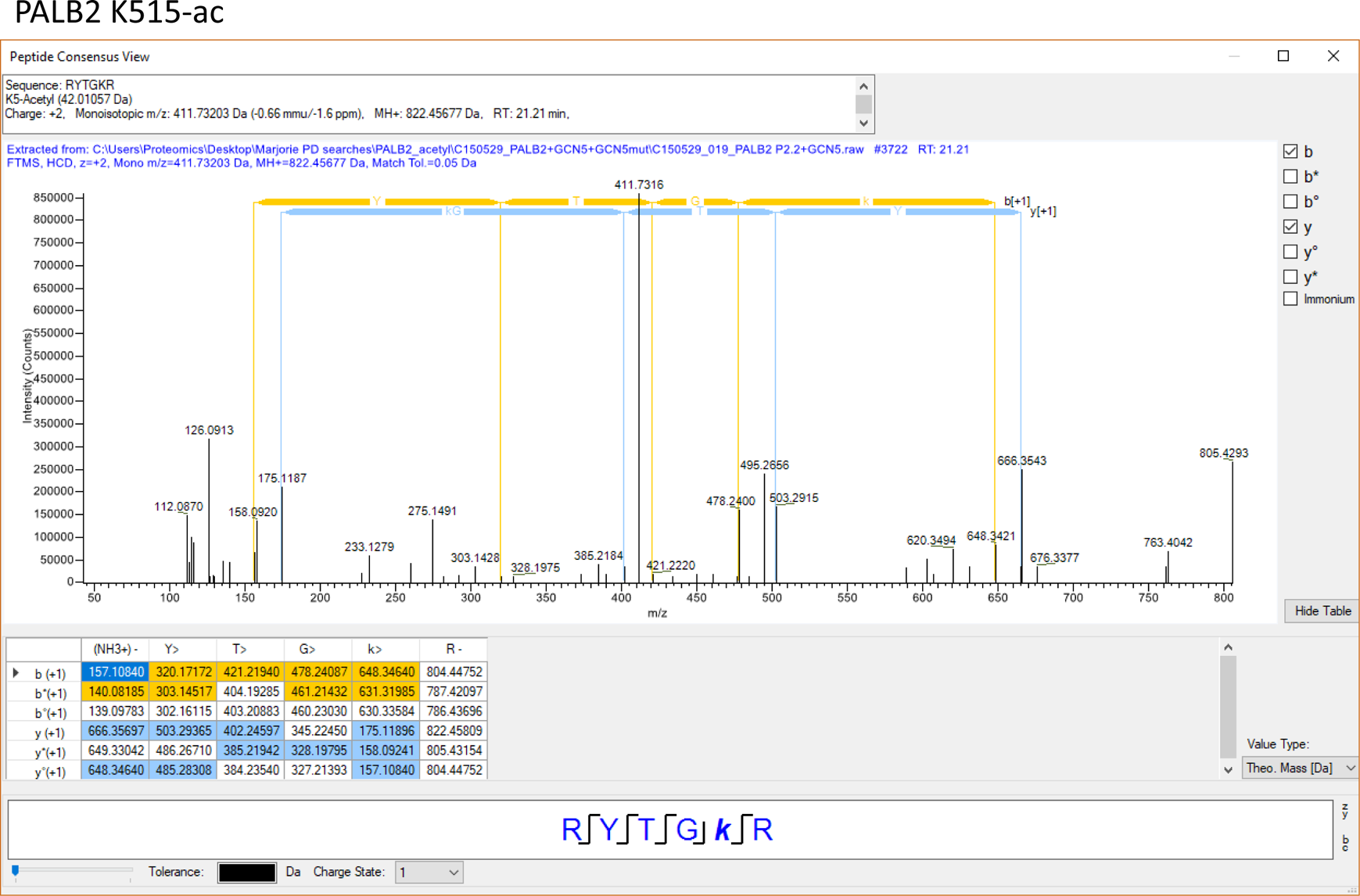

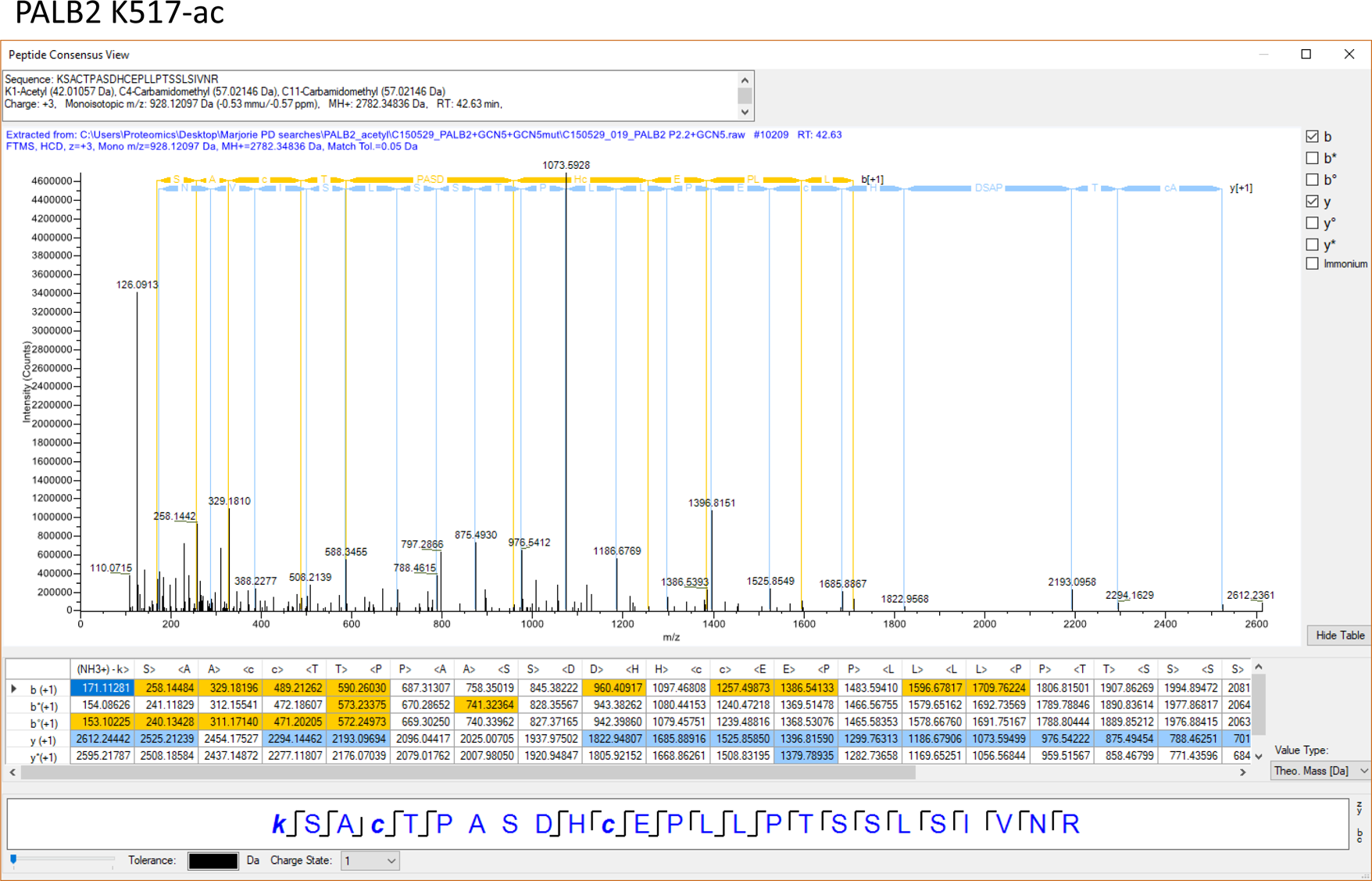

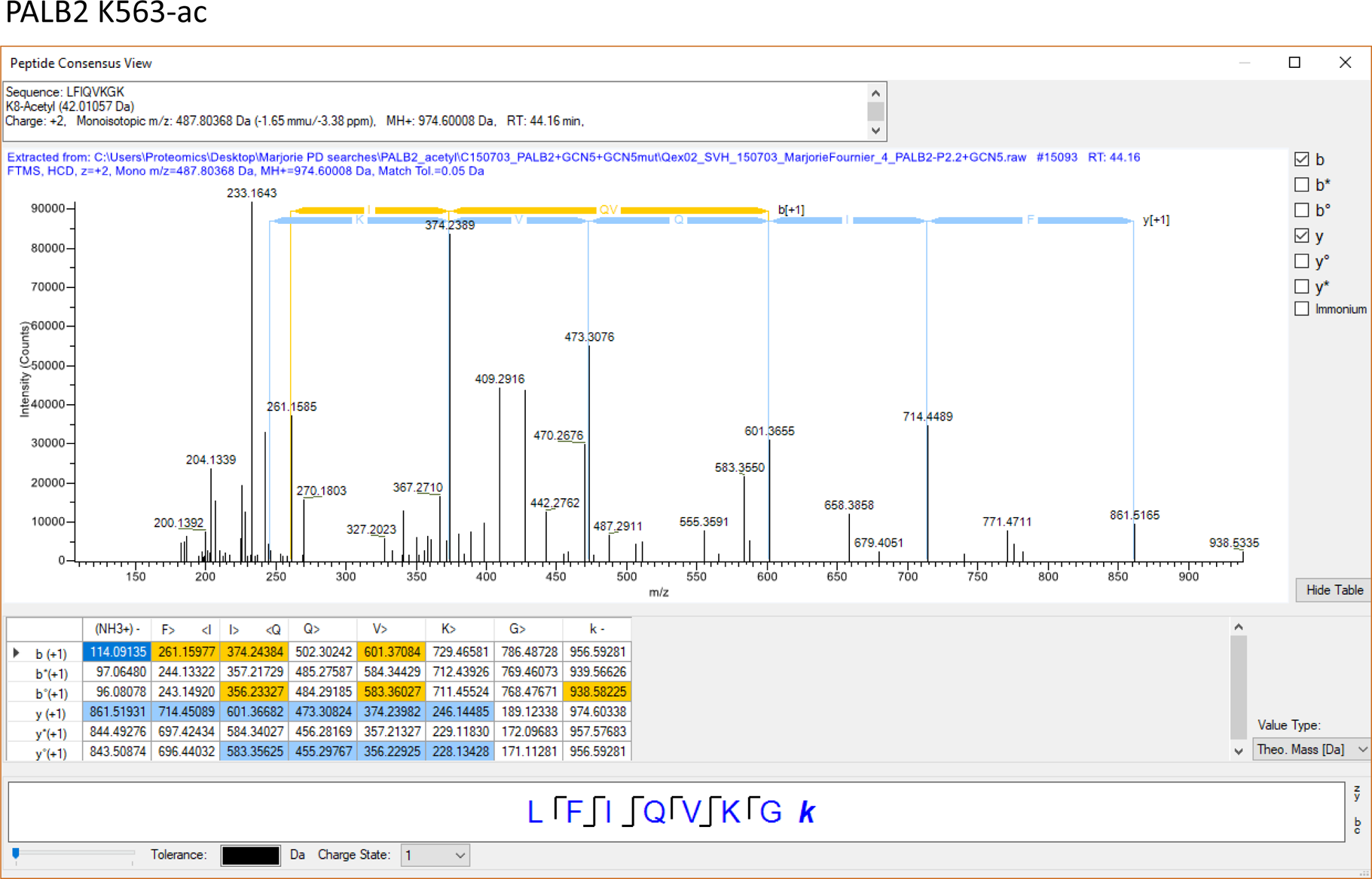

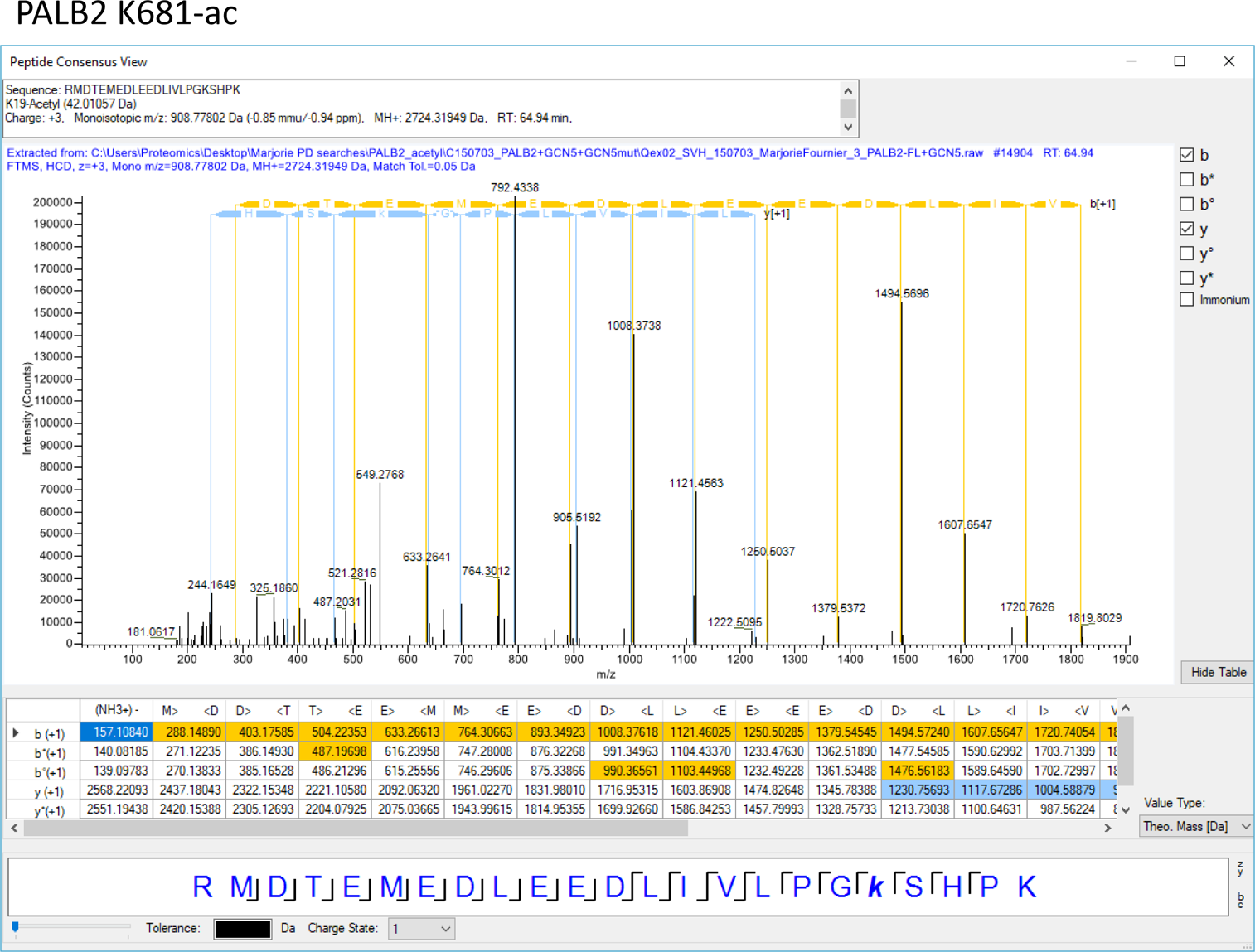

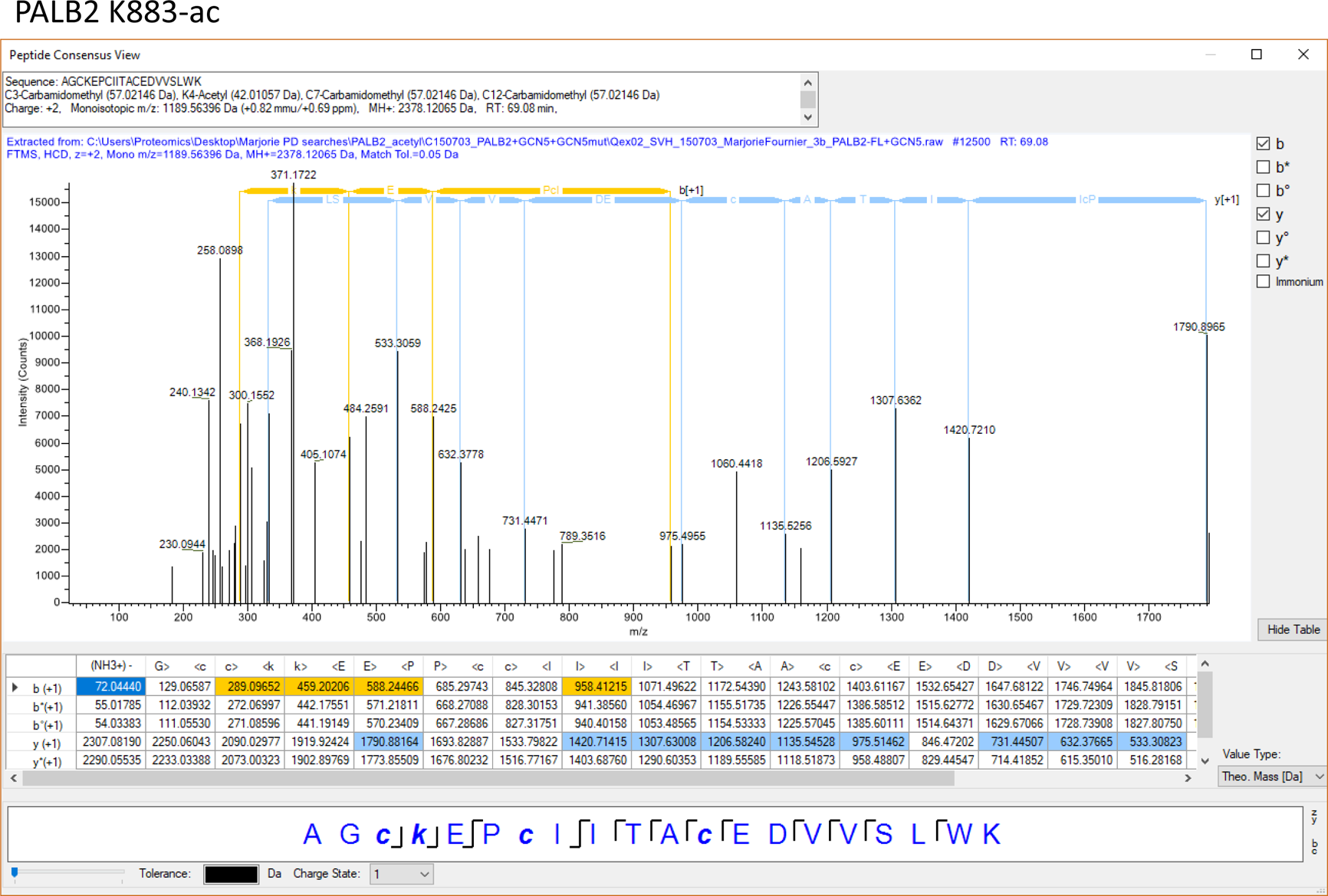

## Notes

#### Summary of Updates

The previous version demonstrates that PALB2 de-acetylation promotes HR repair. In the revised manuscript, we further included phenotypes of cells expressing non-acetylatable PALB2, showing that the acetylation is equally important for genome integrity control in unperturbed cells. These together highlight that the dynamic regulation of PALB2 acetylation is critical for safeguarding genome integrity, both in damaged and undamaged cells.

## References

1. Ahlskog JK, Larsen BD, Achanta K, Sorensen CS (2016) ATM/ATR-mediated phosphorylation of PALB2 promotes RAD51 function. EMBO Rep 17: 671–681

2. Bleuyard JY, Buisson R, Masson JY, Esashi F (2012) ChAM, a novel motif that mediates PALB2 intrinsic chromatin binding and facilitates DNA repair. EMBO Rep 13: 135–141

3. Bleuyard JY, Butler RM, Esashi F (2017a) Perturbation of PALB2 function by the T413S mutation found in small cell lung cancer. Wellcome Open Res 2: 110

4. Bleuyard JY, Fournier M, Nakato R, Couturier AM, Katou Y, Ralf C, Hester SS, Dominguez D, Rhodes D, Humphrey TC, Shirahige K, Esashi F (2017b) MRG15-mediated tethering of PALB2 to unperturbed chromatin protects active genes from genotoxic stress. Proc Natl Acad Sci U S A 114: 7671–7676

5. Buisson R, Niraj J, Rodrigue A, Ho CK, Kreuzer J, Foo TK, Hardy EJ, Dellaire G, Haas W, Xia B, Masson JY, Zou L (2017) Coupling of Homologous Recombination and the Checkpoint by ATR. Mol Cell 65: 336–346

6. Cardona A, Saalfeld S, Schindelin J, Arganda-Carreras I, Preibisch S, Longair M, Tomancak P, Hartenstein V, Douglas RJ (2012) TrakEM2 software for neural circuit reconstruction. PLoS One 7: e38011

7. Dantuma NP, van Attikum H (2016) Spatiotemporal regulation of posttranslational modifications in the DNA damage response. EMBO J 35: 6–23

8. Fournier M, Orpinell M, Grauffel C, Scheer E, Garnier JM, Ye T, Chavant V, Joint M, Esashi F, Dejaegere A, Gonczy P, Tora L (2016) KAT2A/KAT2B-targeted acetylome reveals a role for PLK4 acetylation in preventing centrosome amplification. Nature Comm 7: 13227

9. Guo Y, Feng W, Sy SM, Huen MS (2015) ATM-dependent Phosphorylation of the Fanconi Anemia Protein PALB2 Promotes the DNA Damage Response. J Biol Chem 290: 27545–27556

10. Hayakawa T, Zhang F, Hayakawa N, Ohtani Y, Shinmyozu K, Nakayama J, Andreassen PR (2010) MRG15 binds directly to PALB2 and stimulates homology-directed repair of chromosomal breaks. J Cell Sci 123: 1124–1130

11. Jeyasekharan AD, Ayoub N, Mahen R, Ries J, Esposito A, Rajendra E, Hattori H, Kulkarni RP, Venkitaraman AR (2010) DNA damage regulates the mobility of Brca2 within the nucleoplasm of living cells. Proc Natl Acad Sci U S A 107: 21937–21942

12. Jones S, Hruban RH, Kamiyama M, Borges M, Zhang X, Parsons DW, Lin JC, Palmisano E, Brune K, Jaffee EM, Iacobuzio-Donahue CA, Maitra A, Parmigiani G, Kern SE, Velculescu VE, Kinzler KW, Vogelstein B, Eshleman JR, Goggins M, Klein AP (2009) Exomic sequencing identifies PALB2 as a pancreatic cancer susceptibility gene. Science 324: 217

13. Lee JV, Carrer A, Shah S, Snyder NW, Wei S, Venneti S, Worth AJ, Yuan ZF, Lim HW, Liu S, Jackson E, Aiello NM, Haas NB, Rebbeck TR, Judkins A, Won KJ, Chodosh LA, Garcia BA, Stanger BZ, Feldman MD, Blair IA, Wellen KE (2014) Akt-dependent metabolic reprogramming regulates tumor cell histone acetylation. Cell Metab 20: 306–319

14. Nagy Z, Riss A, Fujiyama S, Krebs A, Orpinell M, Jansen P, Cohen A, Stunnenberg HG, Kato S, Tora L (2010) The metazoan ATAC and SAGA coactivator HAT complexes regulate different sets of inducible target genes. Cell Mol Life Sci 67: 611–628

15. Nowell PC (1976) The clonal evolution of tumor cell populations. Science 194: 23–28

16. O’Donovan PJ, Livingston DM (2010) BRCA1 and BRCA2: breast/ovarian cancer susceptibility gene products and participants in DNA double-strand break repair. Carcinogenesis 31: 961–967

17. Orthwein A, Noordermeer SM, Wilson MD, Landry S, Enchev RI, Sherker A, Munro M, Pinder J, Salsman J, Dellaire G, Xia B, Peter M, Durocher D (2015) A mechanism for the suppression of homologous recombination in G1 cells. Nature 528: 422–426

18. Perez-Riverol Y, Csordas A, Bai J, Bernal-Llinares M, Hewapathirana S, Kundu DJ, Inuganti A, Griss J, Mayer G, Eisenacher M, Perez E, Uszkoreit J, Pfeuffer J, Sachsenberg T, Yilmaz S, Tiwary S, Cox J, Audain E, Walzer M, Jarnuczak AF, Ternent T, Brazma A, Vizcaino JA (2019) The PRIDE database and related tools and resources in 2019: improving support for quantification data. Nucleic Acids Res 47: D442–D450

19. Rahman N, Seal S, Thompson D, Kelly P, Renwick A, Elliott A, Reid S, Spanova K, Barfoot R, Chagtai T, Jayatilake H, McGuffog L, Hanks S, Evans DG, Eccles D, Breast Cancer Susceptibility C, Easton DF, Stratton MR (2007) PALB2, which encodes a BRCA2-interacting protein, is a breast cancer susceptibility gene. Nat Genet 39: 165–167

20. Reid S, Schindler D, Hanenberg H, Barker K, Hanks S, Kalb R, Neveling K, Kelly P, Seal S, Freund M, Wurm M, Batish SD, Lach FP, Yetgin S, Neitzel H, Ariffin H, Tischkowitz M, Mathew CG, Auerbach AD, Rahman N (2007) Biallelic mutations in PALB2 cause Fanconi anemia subtype FA-N and predispose to childhood cancer. Nature Genet 39: 162–164

21. Schneider CA, Rasband WS, Eliceiri KW (2012) NIH Image to ImageJ: 25 years of image analysis. Nat Methods 9: 671–675

22. Srivastava V, Modi P, Tripathi V, Mudgal R, De S, Sengupta S (2009) BLM helicase stimulates the ATPase and chromatin-remodeling activities of RAD54. J Cell Sci 122: 3093–3103

23. Sy SM, Huen MS, Chen J (2009a) MRG15 is a novel PALB2-interacting factor involved in homologous recombination. J Biol Chem 284: 21127–21131

24. Sy SM, Huen MS, Chen J (2009b) PALB2 is an integral component of the BRCA complex required for homologous recombination repair. Proc Natl Acad Sci U S A 106: 7155–7160

25. Tanner KG, Langer MR, Denu JM (2000) Kinetic mechanism of human histone acetyltransferase P/CAF. Biochemistry 39: 11961–11969

26. Wellen KE, Hatzivassiliou G, Sachdeva UM, Bui TV, Cross JR, Thompson CB (2009) ATP-citrate lyase links cellular metabolism to histone acetylation. Science 324: 1076–1080

27. Xia B, Dorsman JC, Ameziane N, de Vries Y, Rooimans MA, Sheng Q, Pals G, Errami A, Gluckman E, Llera J, Wang W, Livingston DM, Joenje H, de Winter JP (2007) Fanconi anemia is associated with a defect in the BRCA2 partner PALB2. Nat Genet 39: 159–161

28. Xia B, Sheng Q, Nakanishi K, Ohashi A, Wu J, Christ N, Liu X, Jasin M, Couch FJ, Livingston DM (2006) Control of BRCA2 cellular and clinical functions by a nuclear partner, PALB2. Mol Cell 22: 719–729

29. Yata K, Bleuyard JY, Nakato R, Ralf C, Katou Y, Schwab RA, Niedzwiedz W, Shirahige K, Esashi F (2014) BRCA2 coordinates the activities of cell-cycle kinases to promote genome stability. Cell Rep 7: 1547–1559

30. Yu DS, Sonoda E, Takeda S, Huang CL, Pellegrini L, Blundell TL, Venkitaraman AR (2003) Dynamic control of Rad51 recombinase by self-association and interaction with BRCA2. Mol Cell 12: 1029–1041

## References

1. Bleuyard JY, Fournier M, Nakato R, Couturier AM, Katou Y, Ralf C, Hester SS, Dominguez D, Rhodes D, Humphrey TC, Shirahige K, Esashi F (2017) MRG15-mediated tethering of PALB2 to unperturbed chromatin protects active genes from genotoxic stress. Proc Natl Acad Sci U S A 114: 7671–7676

2. Esashi F, Christ N, Gannon J, Liu Y, Hunt T, Jasin M, West SC (2005) CDK-dependent phosphorylation of BRCA2 as a regulatory mechanism for recombinational repair. Nature 434: 598–604

